# Single-cell transcriptome reveals insights into the development and function of the zebrafish ovary

**DOI:** 10.1101/2021.12.01.470669

**Authors:** Yulong Liu, Michelle E. Kossack, Matthew E. McFaul, Lana Christensen, Stefan Siebert, Sydney R. Wyatt, Caramai Kamei, Samuel Horst, Nayeli Arroyo, Iain Drummond, Celina E. Juliano, Bruce W. Draper

## Abstract

Zebrafish are an established research organism that has made many contributions to our understanding of vertebrate tissue and organ development, yet there are still significant gaps in our understanding of the genes that regulate gonad development, sex, and reproduction. Unlike the development of many organs, such as the brain and heart that form during the first few days of development, zebrafish gonads do not begin to form until the larval stage (≥5 dpf). Thus, forward genetic screens have identified very few genes required for gonad development. In addition, bulk RNA sequencing studies which identify genes expressed in the gonads do not have the resolution necessary to define minor cell populations that may play significant roles in development and function of these organs. To overcome these limitations, we have used single-cell RNA sequencing to determine the transcriptomes of cells isolated from juvenile zebrafish ovaries. This resulted in the profiles of 10,658 germ cells and 14,431 somatic cells. Our germ cell data represents all developmental stages from germline stem cells to early meiotic oocytes. Our somatic cell data represents all known somatic cell types, including follicle cells, theca cells and interstitial stromal cells. Further analysis revealed an unexpected number of cell subpopulations within these broadly defined cell types. To further define their functional significance, we determined the location of these cell subpopulations within the ovary. Finally, for select examples, we used gene knockout experiments to determine the role of newly identified genes. Our results reveal novel insights into ovarian development and function and the sequencing information will provide a valuable resource for future studies.

## INTRODUCTION

Over the last several decades, the zebrafish has emerged as a model to study vertebrate gonad development and function. Zebrafish ovaries and testes contain homologs of most cell types present in mammalian gonads (e.g. Sertoli and Leydig cells in testes, and follicle and theca cells in ovaries). As in mammals, the zebrafish ovary and testis form from a gonad primordium that is initially bipotential (reviewed in Siegfried and Draper, 2020). The bipotential stage in zebrafish lasts until approximately 15-20 days post fertilization when sex is determined. Though the mechanism by which sex is determined in domesticated zebrafish is not known, zebrafish have orthologs of most genes that drive sex differentiation in mammals. In several cases mutational analysis has revealed these genes play conserved roles in sex determination and differentiation. Examples include the male promoting gene *dmrt1* and the female promoting genes *wnt4a*, *foxl2a* and *foxl2b* (Webster et al., 2018; Kossack et al., 2019; Yang et al., 2017). Though the embryonic origin of the bipotential gonad has yet to be determined in zebrafish, orthologs of genes that are expressed in, and required for, the development of the bipotential gonad in mammals, such as *Gata4* and *Wt1*, are also expressed in the bipotential gonad in zebrafish (Leerberg et al., 2017). Together these data argue that gonad development in vertebrates is regulated by a largely conserved genetic program.

One major difference between zebrafish and mammalian reproduction is that the mammalian ovary produced a finite number of oocytes only during embryogenesis, by contrast, in many teleost (bony fish), such as the zebrafish, adult females can produce new oocytes throughout their lifetime due to the presence of self-renewing germline stem cells (GSCs) (Beer and Draper, 2013, Cao et al., 2019a). Thus the zebrafish adult ovary contains germ cells at all stages of development, from premeiotic germ cells to mature eggs while the mammalian oocytes arrest in prophase of meiosis I (also called dictyate arrest) around the time of birth (Selman et al., 1993, Faddy et al., 1987). The ability to continuously produce new germ cells throughout their lifetime makes female zebrafish an excellent research organism to study female germline stem cells and the somatic cells that regulate their development.

The unipotent GSCs can divide both to replenish its numbers while also producing cells that differentiate into sex-specific gametes, either eggs or sperm. In organisms where GSCs have been identified, they localize to a special microenvironment called the GSC niche. This niche is crucial for maintaining most GSCs in an undifferentiated state while also allowing some to differentiate. GSCs and their niche have been previously identified in a limited number of organisms, such as *Drosophila*, *Caenorhabditis elegans* and male mice (Xie and Spradling, 2000). Identified GSC niches are maintained by specialized somatic cells that produce short-range signals that regulate GSC proliferation and differentiation. For example, in *Drosophila*, somatic cap cells in the ovary maintain GSCs by expressing specific ligands, such as *decapentaplegic* (*dpp,* mammalian *Bmp*), to prevent GSC differentiation (Xie and Spradling, 2000; Song et al., 2004); while in the mouse testis, Sertoli cells in the seminiferous tubules express Glial Derived Neurotrophic Factor (GDNF) which is required for GSC self-renewal (Meng et al., 2000).

GSCs have been identified in the zebrafish ovary, but little is known about the niche and the mechanism by which GSCs are maintained. In the adult ovary, early germ cells localize to a discrete region on the surface of the ovary, called the germinal zone, which has been proposed to be the GSC niche (Beer and Draper, 2013). The germinal zone contains mitotically dividing GSCs, oocyte progenitor cells, and early meiotic oocytes (Draper et al., 2007; Beer and Draper, 2013). However, neither the GSC-intrinsic gene expression landscape, nor the extrinsic GSC niche cells that likely regulate early GSC differentiation in zebrafish have been characterized.

The best characterized somatic cell type in the zebrafish ovary are the follicle cells. Teleost follicle cells are homologous to mammalian granulosa cells and thus are the major oocyte support cell. Follicle and granulosa cells produce signals that regulate oocyte development and maturation, and also produce nutrients that are transported into the oocyte cytoplasm via gap junctions (Su et al., 2009). Unlike granulosa cells, which form a multi-cell layed complex surrounding the mammalian oocyte, follicle cells in fish form only a single-cell layer around the oocytes. In mammals granulosa cells are the major source of ovarian estradiol (E2) production (Ryan, 1979), and this is likely to be the case in zebrafish. In addition to follicle cells, E2 production also requires steroidogenic theca cells which produce androstenedione, the precursor that follicle cells use for E2 synthesis (ref). Theca cells are found around and between oocytes but are otherwise not well characterized in zebrafish. The final major cell type in the ovary are the stromal cells, which are composed of all other somatic cell types present, including connective tissue, blood vessels, and immune cells. The fibroblast-like interstitial stromal cells play a largely structural role but the roles of the other stromal cell types in ovarian development and function are not well characterized in any vertebrates. Although these general cell populations have been observed in the zebrafish ovary, there are significant gaps in knowledge regarding the identification of the full diversity of cell types present in the ovary or the functional roles they play.

In this study, we characterize the complex cell environment in the zebrafish ovary using single-cell RNA sequencing. This data has allowed us to identify all known stages of germ cells, from germline stem cells to pre-follicle stage oocytes (Stage IA; Selman et al., 1993), as well as the somatic cell populations, including follicle, theca, and stromal cells. Using sub-cluster analysis, we have defined subpopulations within these broad cell-type classifications and validated these subpopulations by determining where they reside in the ovary using single-molecule fluorescent *in situ* hybridization (smFISH). Further, we demonstrated the accuracy of this dataset to identify developmentally relevant genes using gene knockout analysis. Our data provide strong support that orthologs of genes involved in mammalian ovary development and function play parallel roles in zebrafish, further supporting zebrafish as a relevant research animal to understand human ovarian development and diseases. This reference data set will serve as a resource to greatly enhance future studies of ovarian development and function in unprecedented detail.

## RESULTS

### Single-cell RNA sequencing identifies all cell types present in the zebrafish juvenile ovary

To identify all cell types and states in the zebrafish ovary, we collected single-cell transcriptomes from ∼25K cells isolated from 40 day post-fertilization (dpf) ovaries. We chose 40 dpf ovaries for three major reasons. First, by 30 dpf sex determination is complete and the initial bipotential gonad has committed to differentiate as either an ovary or testis (Kossack and Draper, 2019). Second, by 40 dpf all major ovarian somatic cell types are likely present (Rodriguez-Mari et al., 2005). Finally, as we were particularly interested in understanding how the developmental progression of germline stem cells is regulated, 40 dpf ovaries have a higher proportion of early-stage germ cells relative to adult ovaries (Draper, 2012).

We prepared three single-cell libraries from dissociated Tg(*piwil1:egfp*)*^uc02^* transgenic ovaries, where eGFP is expressed in all germ cells (Figure1-figure supplement 1; (Leu and Draper, 2010)). Two of these libraries were prepared using a dissociation method that favored isolation of single somatic cells. The third library was prepared from cells isolated using a milder dissociation method that favored germ cell survival, followed by fluorescence-activated cell sorting (FACS) to purify eGFP+ germ cells; (Figure 1-figure supplement 1; see methods for details). We used the 10X Genomics platform for library preparation and sequencing. Following quality control, data cleaning and the removal of red blood cells, doublets, and ambient RNA we recovered a total of 25,089 single-cell transcriptomes that comprise of 10,658 germ cells and 14,431 somatic cells (Figure 1-figure supplement 2; see Methods). We obtained germ cells with an average of 2,510 genes/cell, 11,294 transcripts/cell and somatic cells with an average of 854 genes/cell, 4,812 transcripts/cell.

Cell clustering analysis grouped cells into eight distinct populations using uniform manifold approximation and projection (UMAP; Figure 1, Figure 1-figure supplement 3A, and Table 1). We assigned these clusters provisional cell type identities using the expression of known cell type-specific genes, as follows: germ cells- *deadbox helicase 4* (*ddx4*; formerly *vasa*; (Yoon et al., 1997)); follicle cells- *gonadal soma derived factor* (*gsdf*; (Gautier et al., 2011)); theca cells- *cytochrome P450 cholesterol side-chain cleaving enzyme* (*cyp11a2*; (Parajes et al., 2013)); vasculature- *Fli-1 proto-oncogene* (*fli1a*; (Brown et al., 2000)); neutrophils- *myeloid-specific peroxidase* (*mpx*; (Lieschke et al., 2001)); macrophages- *macrophage-expressed gene 1* (*mpeg1.1*; (Zakrzewska et al., 2010)); NK cells- *NK-lysin tandem duplicate 2* (*nkl.2*; (Carmona et al., 2017; Tang et al., 2017)); Figure 1 and Figure 1-figure supplement 3B). Except for blood vessels and blood cells, the remaining stromal cells are the most ill-defined cell population in the teleost gonad, and no genes have previously been identified that specifically mark this population. Ovarian stromal cells are generally defined as the cells that are not components of the ovarian follicle (i.e. germ and associated follicle and theca cells) and play structural roles (Kinnear et al., 2020). We determined that expression of the gene encoding *collagen, type I, alpha 1a* (*col1a1a*; (Morvan-Dubois et al., 2003)), is highly enriched in this cell population. Collagen is well known for its structural role in tissues and therefore reasoned that the remaining unidentified population of cells in the UMAP graph represented the stromal cells (Figure 1B and Figure 1-figure supplement 3B). Thus, we were able to identify all previously observed and expected cell types in our dataset. A list of genes with enriched expression in each cluster can be found in Table 1.

**Figure 1.**
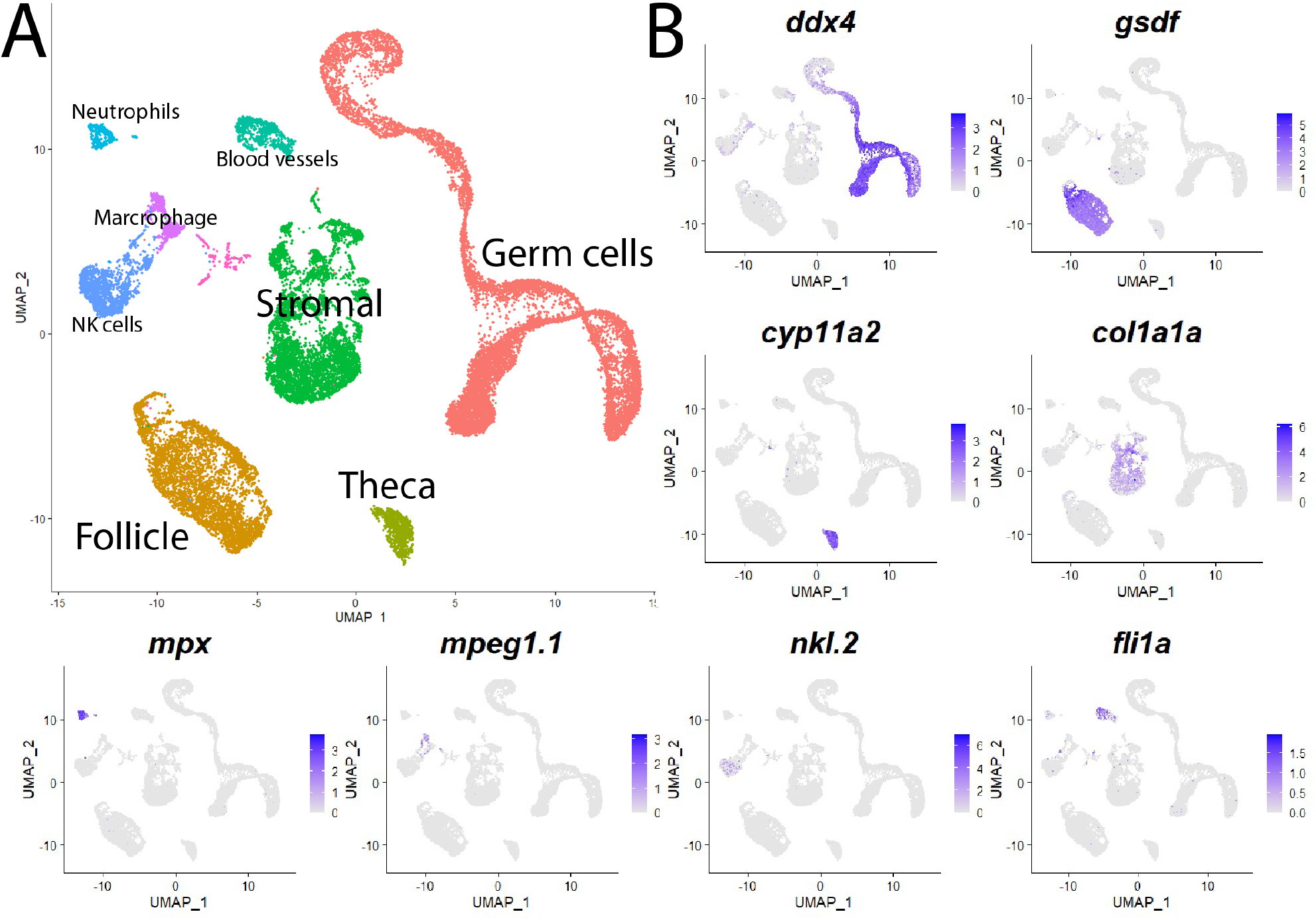
Single-cell RNA sequencing of 25,089 single cells isolated from 40-day old zebrafish ovaries. **A.** Single-cell UMAP plot of the 40-day old zebrafish ovary. Cells are color-coded by computationally determined cell clusters. **B.** Gene expression plots of known cell-specific marker genes identify the major cell type (labeled in **A**) that each cluster corresponds to. Cells expressing the indicated gene are colored purple, and the relative intensity indicates relative expression levels (intensity scale for each plot is on the right).

Our initial analysis revealed genes that are differentially expressed between major cell types but did not distinguish possible sub-populations. For example, *ddx4* and *gsdf* were correctly identified as genes expressed specifically in germ cells and follicle cells, respectively, however specific genes expressed in discrete developmental stages of these populations were not revealed. To gain a more refined view of each of the distinct cell populations we extracted and sub-clustered each population.

### Developmental trajectory of female germ cells

Unlike mammals, female zebrafish are able to produce new oocytes throughout their lifespan due to the presence of GSCs (Beer and Draper, 2013; Cao et al., 2019b). Zebrafish therefore provide a unique opportunity to study female GSCs in a vertebrate. The genes that regulate GSC maintenance or progression of progenitor cells toward differentiation are not well defined in zebrafish. To gain further insight into genes that regulate germ cell development, we performed sub-cluster analysis on the germ cell population (Figure 2A) and the differentiation trajectory was immediately evident in the UMAP. To determine the organization of the cells within the UMAP, we first asked which cells express known stage-specific germ cell markers. Using *nanos2, DNA meiotic recombinase 1* (*dmc1*) and *zona pellucida protein 3b* (*zp3b*), which are expressed in GSCs, early meiotic germ cells, and early oocytes, respectively (Beer and Draper, 2013; Onichtchouk et al., 2003; Yoshida et al., 1998). We found that *nanos2*-expression cells clustered to the left end of the UMAP, *dmc1*-expressing cells were in an intermediate location and *zp3b*-expression cells clustered to the right end of the UMAP. Thus, the cells are organized into what appears to be a developmental trajectory (Figure 2A,B). Because our library preparation method excluded cells larger than 40*μ*m in diameter, it is likely that only Stage IA (pre-follicle stage) or early Stage IB (follicle stage) oocytes are present in our scRNA-seq data set (Selman et al., 1993). To further assess this developmental trajectory, we performed a dedicated trajectory analysis and inferred pseudotime using Monocle 3 (Cao et al., 2019a). We found that the directionality and the sequential gene expression along pseudotime precisely correlated with the trajectory determined by our initial analysis (Figure 2-figure supplement 1). With the developmental trajectory of the germ cells constructed, we were able to identify genes with enriched expression in subsets of germ cells (Table 2). Notably missing from the known stage-specific genes listed above are genes known to be expressed in proliferating oocyte progenitor cells that are intermediate between the GSC and the cells that have entered meiosis (sub-cluster 2 in Figure 2A). In most organisms, oocyte progenitor cells undergo several rounds of mitotic amplifying division before entering meiosis (Pepling et al., 1999). A common feature of oocyte progenitor cells, including those in zebrafish ovaries, is that they divide with complete nuclear division but incomplete cytoplasmic division, thus creating a multi-cell cyst with synchronized developmental progression (Marlow and Mullins, 2008; Pepling et al., 1999). To identify novel markers of oocyte progenitor cells, we analyzed the genes with enriched expression in this sub-population (sub-cluster 2 in Figure 2A; Table 2). Of the top 100 enriched genes, we found only one gene whose expression was both germ cell-specific and restricted to this subpopulation, an unannotated gene, called *zgc:194189* (Table 2). Further sequence analysis revealed that this gene is the zebrafish ortholog of *forkhead-box protein L2 like* (*foxl2l*; Figure 2-figure supplement 2; (Ruzicka et al., 2019). FoxL2l is a paralog of FoxL2, and is present in most teleost genomes as well as in sharks, coelacanths, and spotted gar, but is absent in the genomes of land-dwelling vertebrates (Figure 2-figure supplement 2). Interestingly, medaka *foxl2l* (formerly *foxl3*; Figure 2-figure supplement 2) is expressed in oocyte progenitors and is required for progenitor cells to commit to the oocyte-fate (Nishimura et al., 2015). The *foxl2l* positive cells in our data set also express higher levels of *proliferating cell nuclear antigen* (*pcna*) gene relative to the GSC population (Figures 2B,C), suggesting they divide more rapidly than GSCs, providing further support that this sub-population are zebrafish oocyte progenitors.

**Figure 2.**
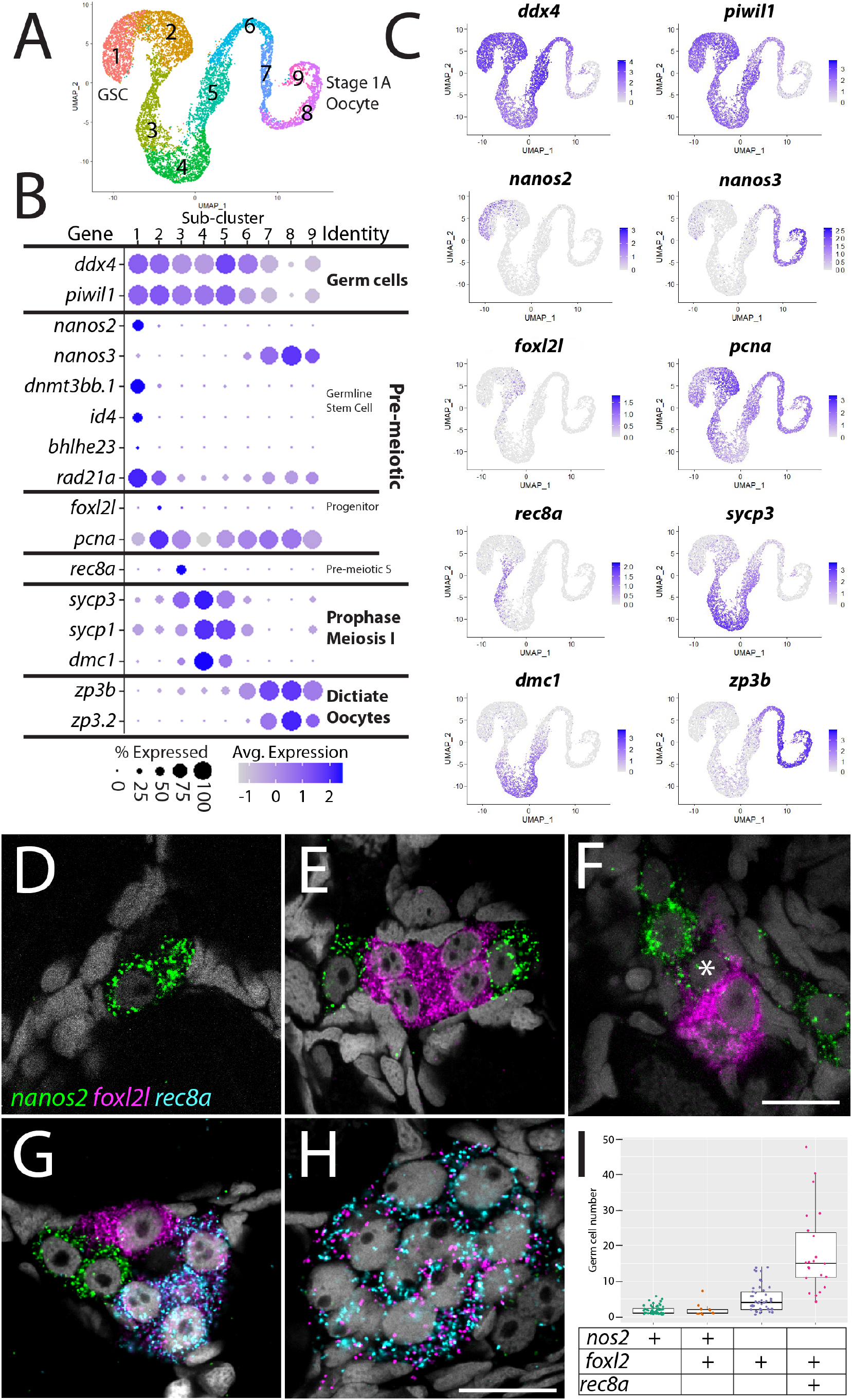
Germ cell sub-cluster analysis reveals developmental transitions of early germ cells. **A.** Germ cell sub-cluster UMAP plot, with cells color-coded by computationally determined cell subtypes. **B.** Dot-plot showing the relative expression of select genes in the germ cell sub-clusters. Some genes, like *ddx4* and *piwil1*, are expressed in all germ cells, while others, such as *nanos2* or *rec8a*, are only expressed in distinct subclusters. **C.** Gene expression UMAP plots of select genes. **D-H**. Triple single-molecule fluorescent *in situ* hybridization for *nanos2* (green), *foxl2l* (magenta) and *rec8a* (blue) in 40 days old zebrafish whole mount ovary. Asterix in **F** indicates a cell double-positive for *nanos2* and *foxl2l.* **I.** Cell number quantification of individual cysts that express the genes indicated on Y-axis. *n*= 70, N=3. Scale bar in F, for D-G, 10*μ*m; in H, 10*μ*m.

To test the hypothesis that *foxl2l*-expressing cells are oocyte progenitor cells, we used single molecule fluorescent *in situ* hybridization (smFISH) to determine where these cells are located in the 40 dpf ovary. Specifically, we determined the location of sub-cluster 2 cells relative to GSCs (sub-cluster #1) and to cells that are initiating meiosis (sub-cluster 3). We identified GSCs using *nanos2* expression and early meiotic cells using *meiotic recombination protein 8a* (*rec8a*) expression. *nanos2* encodes an RNA binding protein that is expressed specifically in germline stem cells (Beer and Draper, 2013) while *rec8a* encodes a meiosis-specific member of the Rad21 cohesin family that is loaded onto chromosomes during pre-meiotic S-phase (Crespo et al., 2019). Our trajectory analysis indicated that *rec8a* expressing cells would be more developmentally advanced than those that express *foxl2l*, as would be expected given its suspected role in pre-meiotic S-phase (Figures 2B,C). Indeed, we found that *foxl2l-* and *rec8a*-expressing cells localize within multi-cell clusters (Figures 2E-H), consistent with our hypothesis that *foxl2l* is expressed in oocyte progenitors, whereas *nanos2-*expressing cells are found predominantly as single cells or doublets, as previously shown (Figures 2D-G; (Beer and Draper, 2013). We found cells expressing *nanos2*, *foxl2l,* and *rec8a* often clustered near one another, (Figure 2G). Also consistent with our hypothesis, *foxl2l* positive cells were found as clusters of ≥4 cells and *rec8a* positive cells were found in clusters of ≥8 cells (Figures 2I). Interestingly, we rarely found cells that were double-positive for *nanos2* and *foxl2l*, and when we did, they had minimal expression of either gene, suggesting these cells were differentiating from GSC to oocyte progenitor (Figure 2F). Cells expressing higher levels of *foxl2l* could be divided into two populations, based on morphology and gene expression. The first consisted of cells within smaller clusters (2-4 cells) that expressed high levels of *foxl2l,* but not *rec8a.* These cells are likely early oocyte progenitors (Figures 2E,G). The second consisted of cells within larger clusters (≤8 cells), expressed relatively less *foxl2l* but also expressed *rec8a*. These cells were likely late oocyte progenitors, including those that are undergoing pre-meiotic S-phase (Figures 2G,H, I). Thus, we identified gene expression patterns that define the pre-meiotic germ cell populations in the ovary.

To determine the function of *foxl2l*, we used CRISPR/Cas9-mediated gene editing to recombine a viral-2A-eGFP insert in-frame into the *foxl2l* locus, resulting in the Tg(*foxl2l:foxl2l-2A-egfp*)*^uc91^* allele (Wierson et al., 2020). The resulting fusion protein encodes all but the last 27 of 239 amino acids of the Foxl2l protein (Figure 3A). We found that in heterozygous animals, GFP was expressed in a subset of premeiotic germ cells in the ovary (Figure 3B), identical to the pattern determined using smFISH (Figure 2E-H). To assess the role of *foxl2l* in germ cell development, we crossed heterozygous parents to produce homozygous mutant offspring. We found that heterozygous knock-in animals had normal sex ratios as adults (Figures 3C-E). By contrast, all homozygous knock-in mutant animals were fertile males as adults, indicating that *foxl2l* is required for female development (Figures 3C,F; *n=* 274, N=4). In zebrafish, the ability to produce oocytes is a requirement for female development, thus loss of oocyte production, which occurs in *foxl2l* mutants, results in all-male development (Rodriguez-Mari et al., 2010; Shive et al., 2010). This is homologous to the role of *foxl2l* in medaka, which is expressed in oocyte progenitor cells and is required for these cells to commit to oogenesis, and upon loss of *foxl2l* these cells instead commit to spermatogenesis (Nishimura et al., 2015).

**Figure 3.**
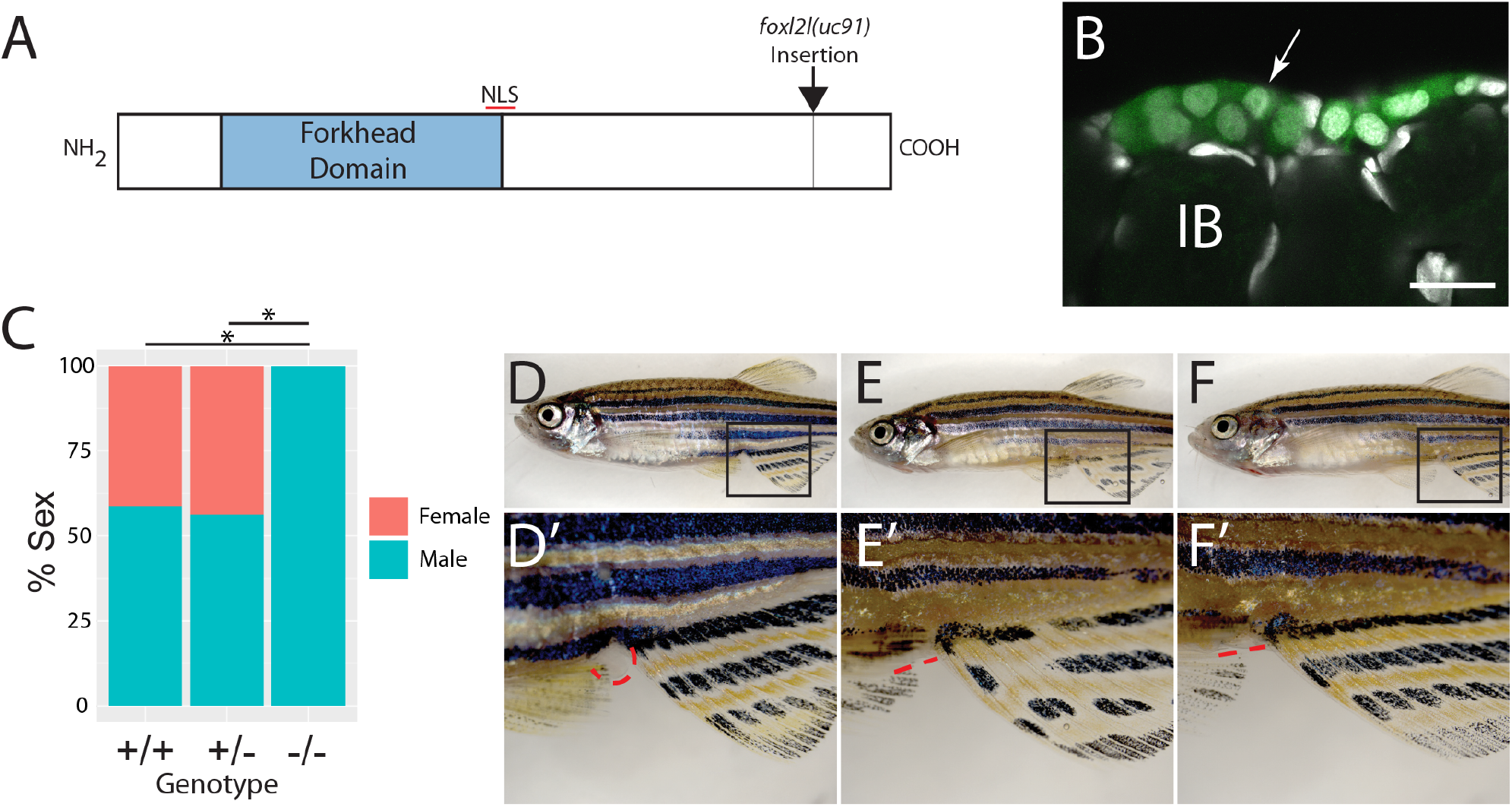
Mutational analysis of *foxl2l*. **A.** Schematic diagram of the Foxl2l protein showing the DNA binding Forkhead homology domain (blue), the location of the nuclear localization signal (NLS), and the viral-2A-*egfp* insertion site in the *foxl2l*(*uc91*) allele. **B.** GFP expression in germ cells from a *foxl2l*(*uc91*) knock-in allele heterozygote recapitulates endogenous *foxl2l* expression (compare to Figure 2E-F). **C.** Sex ratios of *foxl2l*(*uc91*) heterozygotes and homozygotes. **D-F** Representative light micrographs of fish examined in **C.** Wild-type adult female zebrafish (**D**), has characteristic light-yellow pigmentation on ventral belly and a prominent anal papilla (highlighted with red dashed lines) (**D’**). **E.** Wild-type adult male zebrafish (**E**) has dark yellow pigmentation on ventral belly and lacks an anal papilla (highlighted with red dashed lines) (**E’**). *foxl2l*(*uc91*) homozygous mutant (**F**) is phenotypically male. IB, stage IB oocyte.

Female germline stem cells have not been well characterized in any vertebrate. To date only two genes have been shown to play a cell autonomous role in female GSC development in zebrafish-*nanos2* and *nanos3* (Beer and Draper, 2013; Cao et al., 2019b; Draper et al., 2007). *nanos2* and *nanos3* encode conserved RNA binding proteins. In zebrafish, *nanos2* is expressed specifically in GSC in both male and females and loss of *nanos2* leads to sterile males (Beer and Draper, 2013; Cao et al., 2019b). By contrast, *nanos3* is only expressed in females where it plays two separate roles. First, *nanos3* (formally called *nanos1*) is a maternally expressed gene whose mRNA localizes to primordial germ cells (PGC) and loss of maternal function leads to loss of PGCs during early development (Draper et al., 2007; Koprunner et al., 2001). Second, *nanos3* mutant females fail to maintain oocyte production as adults due to loss of *nanos2*-expressing GSCs (Beer and Draper, 2013; Draper et al., 2007). While genetic mosaic analysis established that *nanos3* was required cell autonomously for GSC maintenance (Beer and Draper, 2013), to date *nanos3* expression in premeiotic germ cells in the ovary has not been demonstrated. To address this, we identified the *nanos3-*expressing cells in our data set. As previously reported, *nanos3* is expressed at high levels in early oocytes, but we also detected expression in the apparent GSC subpopulation, confirming that *nanos3* is expressed in GSCs as suggested by the *nanos3* mutant phenotype (Figures 2B,C).

In other organisms Nanos proteins function in complex with members of the Pumilio family of RNA binding proteins (De Keuckelaere et al., 2018; Forbes and Lehmann, 1998). The zebrafish genome encodes three orthologs of Pumilio, called Pum1-3, but it is not known which of these proteins function together with Nanos2 or Nanos3 to maintain GSC or PGC development. We found that *pum1* and *pum3*, but not *pum2* are expressed at significant levels in GSCs (Figure 2-figure supplement 3). In addition, *pum1* is predominantly expressed in GSCs while *pum3* is expressed more uniformly in early oocytes (Figure 2-figure supplement 3). This raises the possibility that Pum1 may partner with Nanos2 while Pum3 partners with Nanos3, though it is also possible that Nanos2 could form complexes with both Pum1 and Pum3. Further mutational analysis is required to test these relationships.

We detected the highest average number of unique transcripts in GSCs (3,517 unique transcripts/cell; Figure 2-figure supplement 4) relative to other premeiotic cells (e.g. 2,315 unique transcripts/cell in progenitor cells; Figure 1-figure supplement 4A). To identify potential regulators of GSC development, in addition to *nanos2*, we identified genes in our data set with enriched expression in this cell population (Table 2). They include *inhibitor of DNA binding 4* (*id4*), which encodes a negative regulator of transcription that is also expressed in mouse spermatogonial stem cells, *jagged canonical Notch ligand 2b* (*jag2b*), *dickkopf WNT signaling pathway inhibitor 3b* (*dkk3b*), *DNA methyltransferase protein 3bb.1* (*dnmt3bb.1*), *interferon regulatory factor 10* (*irf10*), and *chromogranin b* (*chgb*; (Gore et al., 2016; Haddon et al., 1998; Hsu et al., 2010; Oatley et al., 2011; Xie et al., 2008); Figure 2-figure supplement 4B). Using *id4* and *chgb* as representatives, we performed smRNA *in situ* to verify the expression pattern in ovaries. We found that all *id4-*expressing cells also express *nanos2* (Figure 2-figure supplement 4C), confirming that *id4* is expressed in GSCs in the 40 dpf ovary. Interestingly, not all *nanos2*-expressing cells were *id4+*. We obtained similar results with *cghb* (Figure 2-figure supplement 4D). Thus we identified several new genes whose expression is specific to GSC. In future studies it will be important to determine if these genes are expressed in both male and female GSC’s or, like *nanos3*, are female-specific.

### Identification of germ cell stage-specific gene modules and possible transcriptional regulators

We next aimed to identify the potential transcription factors that regulate the development of germ cells. As a first step we performed non-zero matrix factorization (NMF) to find gene modules, which are sets of genes that are co-expressed within cell clusters and sub-clusters and may therefore identify co-regulated genes (Figure 2-supplemental figure 5; (Brunet et al., 2004; Farrell et al., 2018; Siebert et al., 2019). We identified 32 gene modules (note: modules 8, 13, 19 and 29 were determined to be low quality gene modules and therefore not considered in our analysis; see methods), and many of the gene modules showed stage-specific enrichment during germ cell development. For example, genes within modules 25, 28, 10, and 9 were specifically enriched in the GSCs, oocyte progenitors, early meiosis, and oocyte clusters, respectively (Figure 2-figure supplement 5A, B). We next performed gene motif enrichment analysis to find potential transcription factor binding sites enriched within 2 Kb 5’ of the transcription start site of top 20% of genes within individual modules (see Methods). We then verified our gene motif enrichment using the *figla* transcription factor as a test case. Folliculogenesis-specific basic helix-loop-helix (bHLH) transcription factor (Figla) is known to regulate folliculogenesis-specific gene expression in mice and zebrafish (Liang et al., 1997; Qin et al., 2018; Soyal et al., 2000). We found *figla* is expressed in late meiotic germ cells and early oocytes (Figure 2-supplemental figure 5C). We searched for modules that contained genes with enriched putative Figla binding sites, and identified module 9, a module that contains genes with enriched expression in early oocytes (e.g. *zp3*; Table 3; Figure 2-figure supplement 5A,C). Therefore, we were confident that NMF and gene motif enrichment analysis could accurately identify important regulatory motifs in germ cells. To identify putative GSC-specific transcription factors, we focused our analysis on modules 7 and 25, which contained genes whose expression were specifically enriched in the GSC cluster (Figure 2-figure supplement 5). We found that genes within these two modules were enriched for putative binding motifs of transcription factors Bhlhe23, early growth response 4 (Egr4), and pre-B-cell leukemia homeobox 3b (Pbx3b) (Figure 2-figure supplement 5A,D). Importantly, GSCs also had enriched expression of Bhlhe23, Egr4, and Pbx3b (Figure 2-figure supplement 5D). Previous studies of *Egr4* deficient mice have shown reduced fertility, but the activities of Bhlhe23 and Pbx3b in GSC development and function have not been reported (Tourtellotte et al., 1999). We performed smRNA *in situ* hybridization to confirm the expression of *bhlhe23* and found expression specifically in a subset of *nanos2*-expressing GSCs (Figure 2-figure supplement 5E). Finally, the motif enrichment analysis also identified multiple genes with known functions in germ cell regulation that contain Bhlhe23 binding motifs, such as *cnot1*, which encoded a component of the CCR4/Not deadenylase complex that functions with Nanos proteins to regulate translation of select targets in GSCs (Suzuki et al., 2012) In addition, *dnmt3bb.1, tsmb4x* and *mcm6* are expressed in GSC and have putative Bhlhe23 sites within 2KB of their transcription start sites, suggesting that Bhlhe23 may regulate the expression of multiple genes (Figure 2-figure supplement 5F).

### Identification of three follicle cell subpopulations

Follicle cells form a single-layered epithelium that encases developing oocytes once they arrest at the diplotene stage of meiotic prophase I. Teleost follicle cells are homologous to mammalian granulosa cells as evidenced by shared functions and gene expression. For example, zebrafish follicle cells express orthologs of many of the core granulosa cell-expressed genes, such as *forkhead-box protein L2* (*foxl2a* and *foxl2b*), *anti-Mullerian hormone* (*amh*), *follicle stimulating hormone receptor* (*fshr*), *aromatase*/*cytochrome P450 19a1a* (*cyp19a1a*), and *notch receptor 3* (*notch3*) (Figure 4B and Figure 4-figure supplement 1; (Crespo et al., 2013; Dranow et al., 2016; Kwok et al., 2005; Prasasya and Mayo, 2018; Rodriguez-Mari et al., 2005). Oocyte and follicle cell development is coordinated through bidirectional oocyte-follicle cell-cell interactions (Kidder and Vanderhyden, 2010). Soon after an oocyte has formed, it is surrounded by follicle cells and becomes dependent on these cells for factors it cannot produce, such as pyruvate, cholesterol, and select amino acids. In some cases, these compounds are transported to the oocyte via connexin-mediated gap junctions (Su et al., 2008). Mammalian granulosa cells are divided into two major cell types, the steroidogenic mural cells that form an outer layer and produce estradiol while cumulus cells that are in direct contact with the oocyte and provide nutrients through gap junctions (Gilchrist et al., 2008). By contrast, teleost follicle cells form only a single cell layer and are the likely source of both estradiol production and select nutrients required for oocyte development (Devlin and Nagahama, 2002). It can be presumed that zebrafish follicle cells also divide and mature along with the oocyte, however, little is known about stage-specific transcriptional changes that may occur in follicle cells in coordination with oocyte developmental progression. Another major difference between mammals and teleosts is that in mammals new oocytes are only formed during embryogenesis, while in many teleosts, including zebrafish, new oocytes are formed continuously from a population of GSCs. Thus, the need to produce new follicle cells extends throughout the reproductive life of the fish. However, pre-follicle cells have not been molecularly identified in any teleost (Beer and Draper, 2013; Cao et al., 2019b).

**Figure 4.**
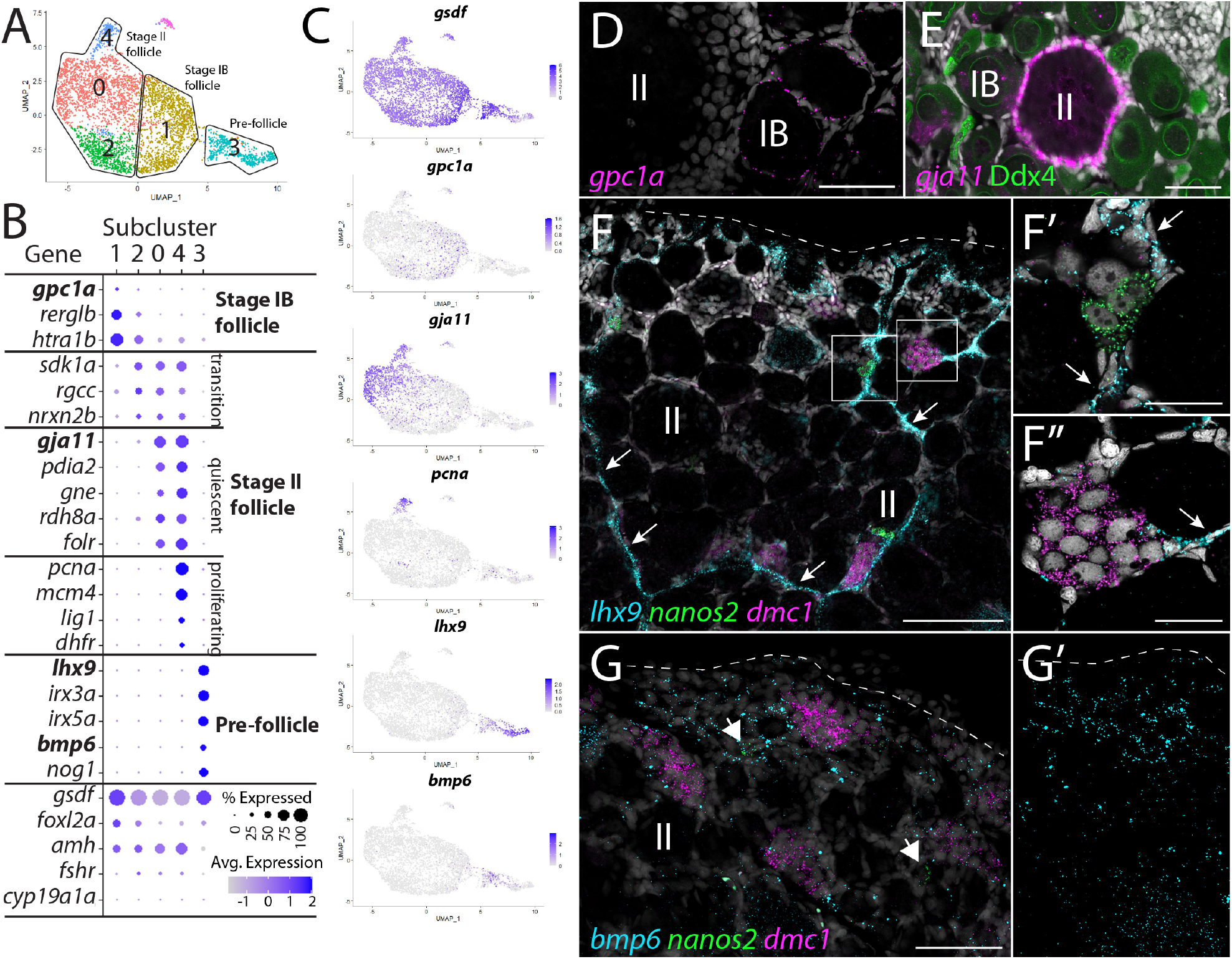
Follicle cell sub-cluster analysis reveals three main cell subtypes. **A.** Follicle cell sub-cluster UMAP plot, with cells color-coded by computationally determined cell subtypes. The three main subtypes are outlined. **B.** Dot-plot showing the relative expression of select genes in the follicle cell sub-clusters. Some genes, like *gsdf*, are expressed in all follicle cells, while others, such as *lhx9*, are only expressed in distinct subclusters. **C.** Gene expression UMAP plots of select genes. Cells expressing the indicated gene are colored purple, and the relative intensity indicates relative expression levels (intensity scale for each plot is on the right). **D-G.** smFISH on whole-mount 40 dpf ovaries reveals the location of cell subtypes. In all panels, DNA is gray. **D.** *gpc1a* expression (pink) is detected in follicle cells surrounding stage IB oocytes, but not stage II oocytes. **E.** *gja11* expression (pink) is detected in follicle cells surrounding stage II oocytes, but not stage IB oocytes. Ddx4 indirect immunofluorescence (green) labels all germ cells. **F.** Triple smFISH shows *lhx9* expressing cells (blue) form tracts on the surface of the ovary (arrows) that colocalize with *nanos2* (green) and *dmc1* (pink) expressing germline stem cells and early meiotic cells, respectively. Lateral edge of the ovary is indicated with a dashed line. **F’** and **F”** Higher magnification views of regions boxed in **F**. **G.** Triple smFISH shows *bmp6* expressing cells (blue) are concentrated near the lateral edge of the ovary, a region that contains *nanos2* (green) and *dmc1* (pink) expressing germline stem cells and early meiotic cells, respectively. **G’**. *bmp6* channel only. Scale bars in D, E and G9 for G and G’), 50 *μ*m; F, 100*μ*m; F’ and F”, 20*μ*m. IB, stage IB oocyte; II, stage II oocyte.

We performed sub-cluster analysis on the *gsdf+* cells to further characterize follicle cells (Figure 4). Because we used 40 dpf ovaries as our source of cells, the follicle cells in our data set likely represent pre-follicle cells and those associated with Stage IB or early-Stage II oocytes. Consistent with this, *cyp19a1a,* a gene known to be expressed in follicle cells surrounding mid-stage II and older oocytes, is only detected in a very small number of follicle cells (Figure 4B and Figure 4-figure supplement 1; (Dranow et al., 2016). We found that the majority of genes detected in follicle cells are expressed uniformly throughout the cluster (e.g. *gsdf*, *foxl2a/b* and *amh*; Figure 4A,B and Figure 4-figure supplement 1). However, our sub-cluster analysis distinguished five cell sub-clusters based on a subset of genes that had non-uniform expression (Figure 4A and Table 3). Upon further analysis we concluded that these sub-clusters likely represent three developmentally relevant cell populations (e.g. sub-cluster 4 represents a mitotic subpopulation of otherwise developmentally similar cells in subclusters 0 and 2; Figure 4A).

#### Stage IB follicle cells (Sub-cluster 1)

Cells within sub-cluster 1 have enriched expression of the heparan sulfate proteoglycan-encoding gene, *glypican 1a* (*gpc1a*), *RERG/RAS-like b* (*rerglb*) and *Htra serine protease 1b (htra1b;* Figures 4A-C). Using *gpc1a* as a representative gene of sub-cluster 1, we performed smFISH and determined that *gpc1a+* cells are found only surrounding stage IB oocytes (Figure 4D; (Selman et al., 1993)), and thus represent the primary follicle cells that first enclose oocytes to form the functional follicle. We did not observe significant expression of genes required for cell division in sub-cluster 1, such as *proliferating cell nuclear antigen* (*pcna*) and *dihydrofolate reductase* (*dhfr*), suggesting that Stage IB follicle cells are not highly proliferative (Figures 4B,C).

#### Stage II follicle cells (Sub-clusters 0, 2 and 4)

Sub-cluster 0 contains the majority of the follicle cells in our data set, which are characterized by the enriched expression of several genes, including *gap junctional protein alpha 11* (*gja11*; formerly *cx43.5*; Figures 4A-C). To determine the location of this cell population within the ovary, we performed smFISH for *gja11*. These results show that follicle cells expressing *gja11* are restricted to Stage II follicles (Figure 4E). Gene expression analysis for sub-cluster 2 cells suggests that these cells are a transitional state between from Stage IB to Stage II follicle cell, as they express several genes, such as s*idekick cell adhesion molecule 1a* (*sdk1a*), that are expressed in both populations (Figure 4B). Gene expression analysis argues that cells within sub-cluster 4 are a mitotic subpopulation of sub-cluster 0 cells, as these cells have nearly identical gene expression with cells in sub-cluster 0 but also express high levels of genes necessary for mitosis, such as *pcna* (Figures 4B,C). Follicle cell proliferation is not surprising given that the oocyte surface area increases 1,400 fold during oogenesis. In mammals there is evidence that granulosa cell proliferation is regulated by pituitary-produced follicle stimulating hormone (Fsh; (Goldenberg et al., 1972)), and consistent with this, we found the gene encoding the Fsh receptor, called *fshr*, is expressed in zebrafish follicle cells including those in sub-cluster 4 (Figure 4-figure supplement 1). Finally, we detect only a small percentage of cells expressing *cyp19a1a* in sub-cluster 0, arguing that cells in this sub-cluster is predominantly composed of follicle cells from early-Stage II oocytes, which do not express *cyp19a1a* (Figure 4B and Figure 4-figure supplement 1; (Dranow et al., 2016).

#### Pre-follicle cells (Sub-cluster 3)

Sub-cluster 3 cells are part of the follicle cell lineage because they express *gsdf* (Gautier et al., 2011; Figures 4A-C), yet these cells also express many genes that are unique to this subpopulation, arguing these cells are distinct from the Stage IB and II follicle cells identified above. Interestingly, these cells express orthologs of the core genes associated with early undifferentiated somatic gonad cells in mammals, including the transcription factors *LIM homeobox 9* (*lhx9*), *Wilms tumor 1a* (*wt1a*), *empty spiracles homeobox 2* (*emx2*), *nuclear receptor 5a1b* (*nr5a1b; formerly sf1b*; Figure 4B, C and Figure 4-figure supplement 2C,D; (Birk et al., 2000; Kreidberg et al., 1993; Luo et al., 1994; Miyamoto et al., 1997). Notably, sub-cluster 3 cells also uniquely express *iroquois homeobox 3a and 5a (irx3a and irx5a*) whose orthologs in mammals are required for pre-granulosa cell development in the embryonic mouse ovary (Figure 4B, C and Figure 4-figure supplement 2C,D; (Kim et al., 2011). Sub-cluster 3 cells do not express *gata4*, a gene required for the earliest stage of somatic gonad cell specification in mice and that is also expressed in early somatic gonad cells during the bipotential phase in zebrafish (10 dpf; (Hu et al., 2013; Leerberg et al., 2017). This argues that these cells are not developmentally similar to the somatic gonad precursors present in the bipotential gonad. Finally, sub-cluster 3 cells express lower levels of genes associated with follicle cell differentiation and function, such as *foxl2a/b*, *amh, fshr* and *notch3*, then do the Stage 1B and Stage II follicle cells (Figure 4B, C and Figure 4-figure supplement 1). Together these results argue strongly that sub-cluster 3 cells are pre-follicle cells.

We noticed that many of the genes with enriched expression in sub-cluster 3 did not appear uniformly expressed in all cells within this sub-cluster. For example, *lhx9* appeared to have higher expression in cells distal to the sub-cluster 1 in the UMAP while *gsdf* and *bmp6* appeared higher in cells proximal to sub-cluster 1 (Fig. 4A, C). To explore this further, we performed additional sub-cluster analysis using only those cells derived from follicle cell sub-clusters 1 and 3 (Figure 4-figure supplement 2A,B and Table 4). This analysis confirmed that sub-cluster 3 cells can be partitioned into two distinct sub-clusters which, for simplicity, we have designated sub-clusters 3.1 and 3.2 (Figure 4-figure supplement 2B). While many genes, such as *emx2*, *irx3a* and *irx5a*, appeared to be expressed uniformly in sub-clusters 3.1 and 3.2, our analysis confirmed that *lhx9* and *wt1a* have enriched expression in sub-cluster 3.1, while *gsdf* and *bmp6* appear enriched in sub-cluster 3.2 (Figure 4-figure supplement 2C,D). Another intriguing difference between these two cell populations is that sub-cluster 3.2 cells have enriched expression of genes that encode cell signaling components, such as *bmp6, fibroblast growth factor receptors 3* and *4* (*fgfr3* and *fgfr4*) and the Fgf-responsive genes *etv4* and *spry4*, while subcluster 3.1 cells have enriched expression of genes that encode cell signaling attenuators, such as the the Bmp ligand antagonists *noggin1 and 3* (*nog1 and nog3*), the Wnt ligand antagonist *dikkopf 1b* (*dkk1b*), and the retinoic acid degrading enzyme encoded by *cytochrome P450 26a1* (*cyp26a1*). In mammals retinoic acid signaling promotes germ cells to enter meiosis (Bowles et al., 2006; Koubova et al., 2006). Therefore, it is possible that inhibition of Bmp, Wnt, and retinoic acid signaling is necessary to keep these cells in an undifferentiated, stem cell-like state, or to influence other surrounding cells.

To determine where these cells reside in the ovary, we performed smFISH, using *lhx9* and *bmp6* as representative genes for clusters 3.1 and 3.2, respectively. Consistent with our hypothesis, we found that *lhx9*+ cells do not localize around oocytes, but instead formed cords-like structures that are located on the surface of the ovary (arrows in Figures 4F,F’,F”). We next asked if these cells were associated with pre-follicle stage germ cells, which would include pre-meiotic and/or early meiotic germ cells. We performed triple smFISH for *lhx9*, GSC-expressed *nanos2* and early meiosis-expressed *dmc1*. Remarkably, we found that early-stage germ cells were always contacting *lhx9*+ pre-follicle cells (*n*=31 clusters of *nanos2* or *dmc1*-expressing cells; N=4). In the medaka ovary, *sox9b*-expressing cells have been identified as pre-follicle cells and similarly formed cord-like structures on the ovarian surface that associate with early-stage germ cells, including GSC’s (Nakamura et al., 2008; Nakamura et al., 2010). Indeed, we found the zebrafish *sox9b* ortholog is expressed in a subset of *lhx9*-expression cells indicating that these cells are likely functional orthologs of the medaka *sox9b*-expressing cells (Figure 4-figure supplement 2). Finally, we determined the location of *bmp6* expressing cells together with *nos2* and *dmc1*. *bmp6* expressing cells also appear to colocalize to regions containing early-stage germ cells, but unlike *lhx9*, the *bmp6* staining was more diffuse, though concentrated towards the lateral edge of the ovary (Figures 4G,G’).

In mammals, retinoic acid signaling is required to induce germ cell entry into meiosis (Bowles et al., 2006; Koubova et al., 2006). Previous studies have argued that retinoic acid is likely produced in the zebrafish ovary by both follicle cells and interstitial cells as these cells express *aldh1a2*/*neckless*, which encodes the enzyme that converts retinaldehyde to retinoic acid (Figure 4-figure supplement 3A; (Rodriguez-Mari et al., 2013)). Given that the retinoic acid degrading enzyme Cyp26a1 appears to be produced by the *lhx9*-expressing pre-follicle cells (Figure 4-figure supplement 2C,D and Figure 4-figure supplement 3B), and the retinoic acid receptor-encoding genes *rxrba* and *rxrbb* are expressed in premeiotic germ cells (Figure 4-figure supplement 3C), we speculate that the close association of premeiotic germ cells with these cells may be important for regulating germline stem cell maintenance.

### Follicle cell-expressed *wnt9b* is required for female sexual development

In addition to their role in support and regulation of oocyte development during oogenesis, follicle cells also play key roles in promoting female sex determination and differentiation in vertebrates. Our lab previously showed that the follicle cell-expressed gene encoding the Wnt4a ligand, was required for female sex differentiation, similar to the role of Wnt4 in mammals (Kossack et al., 2019; Vainio et al., 1999). However, while 95% of *wnt4a* mutants develop as males, we found that ∼5% can develop as females. We reasoned that the partial penetrance of the phenotype could be due to the function of an additional Wnt ligand(s). To further demonstrate the usefulness of this scRNA-seq data set for functional gene discovery, we asked if the expression of other Wnt ligands were detected in the data set and if so, in what cell type(s)? Our scRNA-seq analysis confirmed that *wnt4a* is expressed in follicle cells in zebrafish ovary (Fig. 5A; (Kossack et al., 2019). Additionally, there is significant expression of *wnt8a* in oocytes, *wnt9a* in stromal cells, *wnt9b* in follicle cells, and *wnt11* in stromal and theca cells (Figure 5-figure supplement 1). We focused our attention on *wnt9b* because its expression in follicle cell clusters was strikingly similar to that of *wnt4a* (Figure 5A). To determine if *wnt9b* was involved in sex determination and/or differentiation, we produced *wnt9b* mutants using CRISPR/Cas9. The *wnt9b*(*fb207*) allele contains a 57bp deletion of exon1, which includes the splice donor sequence, and is therefore predicted to be a loss-of-function mutation (Figure 5B).

**Figure 5.**
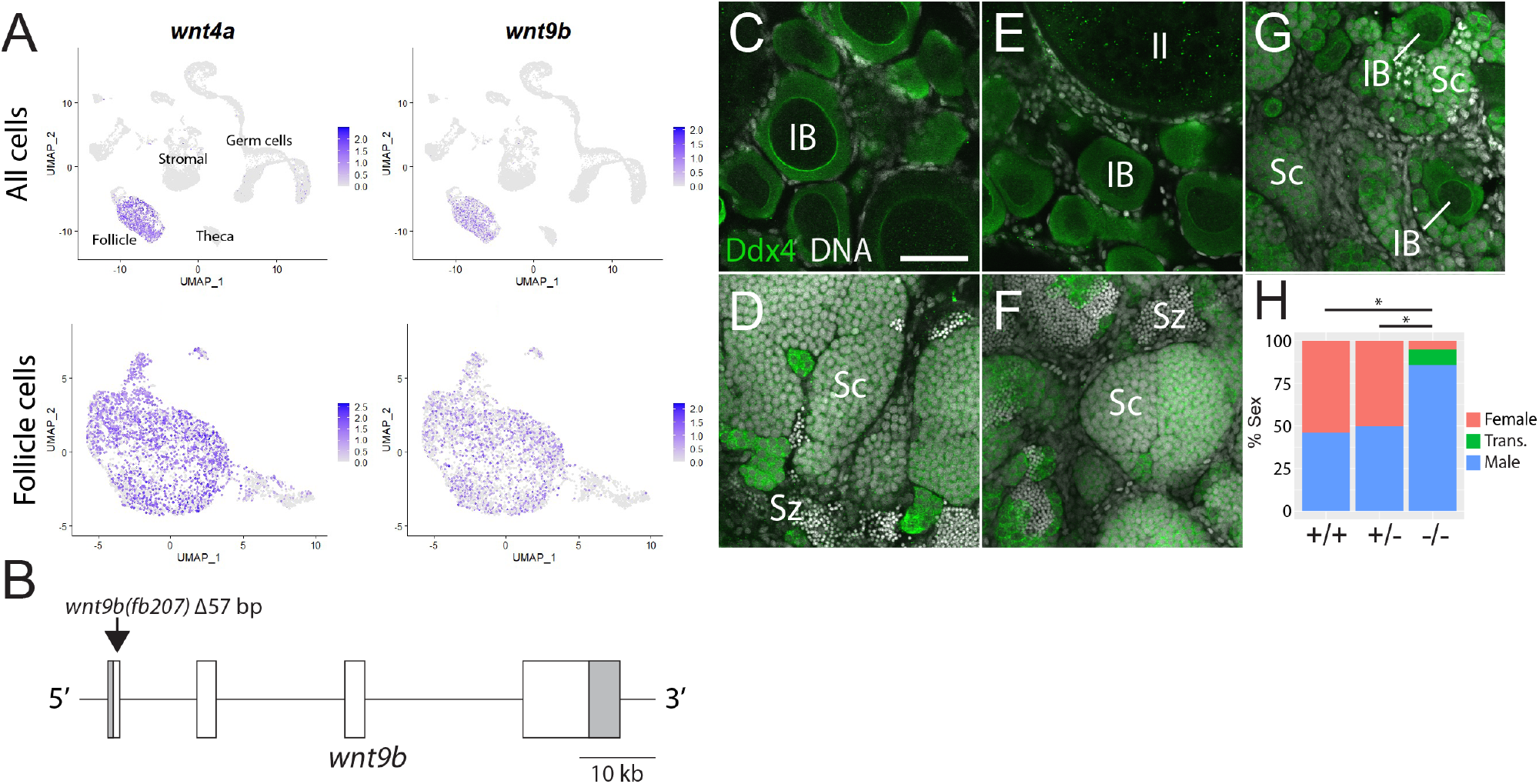
Expression and mutational analysis of *wnt9b*. **A.** Expression plots of *wnt4a* and *wnt9b* show that *wnt9b* is expressed only in follicle cells, in a pattern nearly identical to *wnt4a*. **B.** Schematic of the *wnt9b* genomic locus. Boxes are exons, UTR sequences are shaded. Arrow indicates approximate location of the 57 bp deletion in the *wnt9b*(*fb207*) allele. **C-G,** Representative regions of gonads stained for Ddx4 protein (green) to identify germ cells. *wnt9b*(*+/+*) ovaries (**C**) and testis (**D**). *wnt9b*(*-/-*) ovary (**E**) and testis (**F**). **G.** *wnt9b*(*-/-*) gonad that contains mostly germ cells that have characteristics of spermatogenesis, but also has a few stage IB oocytes. **H.** Sex ratios of offspring produced from *wnt9b(fb207)* heterozygous (+/−) parents (*n*=384, N=3, * p<0.05;). Trans., transitioning; IB, Stage IB oocyte; Sc, spermatocyte; Sz, spermatozoa. Scale bar in C for C-G, 50*μ*m.

Domesticated zebrafish do not have sex chromosomes, instead sex is determined by a combination of genetic and environmental factors (Kossack and Draper, 2019). Because sex ratios in any cross can vary, it is important to compare sex ratios in mutants to their wild-type siblings. We crossed *wnt9b*(*fb207*) heterozygous parents and determined the genotype of their offspring at 50 dpf. We then determined the phenotype by dissecting their gonads to determine whether they had ovaries (females) or testes (males; Figures 5C-G). We found wild-type animals developed similar numbers as male and female, while *wnt9b* mutants were majority male (Figure 5H). In addition, we found a subset of *wnt9b* mutants that had gonads that appeared to be transitioning from an ovary to a testis, as they contained a few stage IB oocytes surrounded by germ cells undergoing early spermatogenesis (Figure 5G). Because oocytes are never observed in wildtype testes at 50 dpf (Figure 5D), the presence of oocyte in these *wnt9b* mutants suggest that these animals initiated female development before sex reverting to males. Consistent with the hypothesis, wildtype testis at 50 dpf contained germ cells at all stages of spermatogenesis, including early spermatocytes and mature spermatozoa (Figure 5D), while the transitioning gonad contained only spermatocytes, but not mature spermatozoa (Figure 5G).

In addition to its role in sex differentiation, we previously found that both male and female *wnt4a* mutants had defects in the development of reproductive ducts that prevented release of mature gametes during spawning (Kossack et al., 2019). By contrast, 78.6% (n=14) of *wnt9b* mutants were able to spawn naturally, a number that is similar to wild-type controls (100%, n=3) arguing that *wnt9b* mutants have functional reproductive ducts. Thus, these data argue that *wnt9b* plays a similar role in female sex determination and/or maintenance to that of *wnt4a*, but unlike *wnt4a*, does not appear to have a role in the development of the reproductive ducts.

### Identification of five stromal cell subpopulations

Ovarian stromal cells can be broadly defined as all ovarian cell types other than germ, follicle, or theca cells. Given this broad definition, it is no surprise that stromal cells are the least characterized cell types in the ovary. Known stromal cell types include interstitial cells, which are collagen-producing connective tissue (fibroblast) that provide structure integrity to the ovary, surface epithelial cells that surround the ovary, vasculature and associated perivasculature cells, and resident blood cell types (Kinnear et al., 2020). To date few of these cell types have been characterized in the zebrafish ovary. From our initial clustering we identified a large population of cells that had enriched expression of several collagen encoding genes, such as *col1a1a* (Figure 1A) and reasoned that these were likely the stromal cell population. This is further supported by the expression of *decorin* (*dcn*), which encodes a proteoglycan component of the extracellular matrix, and *nuclear receptor subfamily 2, group F, member 2* (*nr2f2*, formerly *coup-tfII*) both of which are expressed in mammalian ovarian stromal cells (Figure 6A; (Pereira et al., 1995; Wagner et al., 2020). To further characterize this cell population, we performed sub-cluster analysis and identified five probable cell subpopulations, which are described below (Figure 6A and Table 5). Of the somatic ovarian cells, the stromal cell cluster is the most transcriptionally diverse. This is also supported by the Gene Ontology (GO) analysis.

**Figure 6.**
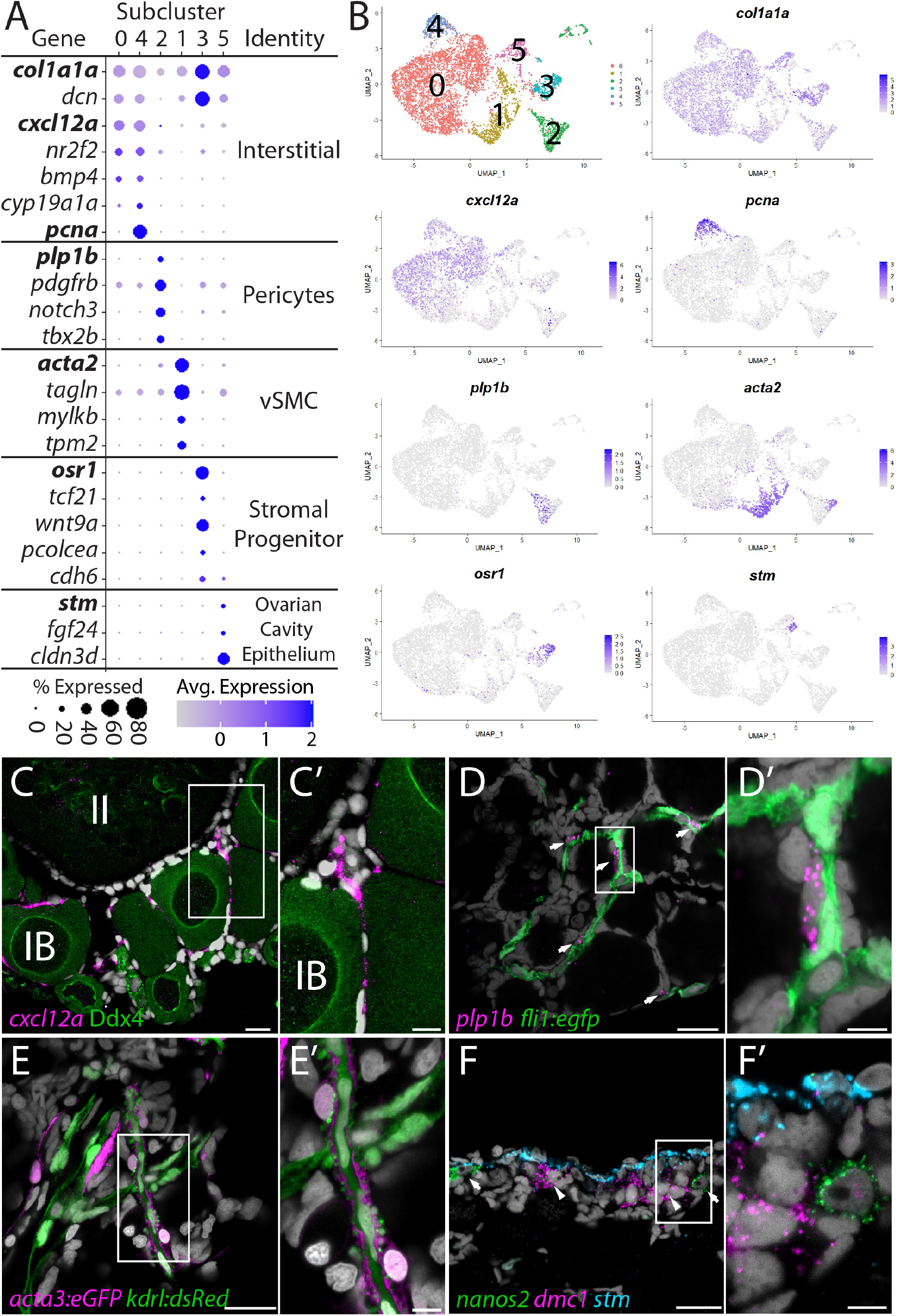
Stromal cell sub-cluster analysis reveals five main cell subtypes. **A.** Dot-plot showing the relative expression of select genes in the stromal cell sub-clusters. Some genes, like *col1a1a*, are expressed in all stromal cells, while others, such as *stm*, are only expressed in a specific subcluster. UMAP plots of genes in bold are shown in **B. B.** Gene expression UMAP plots of select genes. Top left panel shows cells color-coded by computationally determined cell subtype. Cells expressing the indicated gene are colored purple, and the relative intensity indicates relative expression levels (intensity scale for each plot is on the right). **C-G.** smFISH on whole-mount 40 dpf ovaries reveals the location of cell subtypes. In all panels, DNA is gray. **C.** *cxcl12a* expressing interstitial cells localize around early-stage oocytes (≤stage IB), but not around stage II oocytes. Ddx4 indirect immunofluorescence (green) labels all germ cells. **C’.** higher magnification of the region boxed in **C. D.** *plp1b*- expression pericytes co-localize with *fli1:eg*fp expressing blood vessels (green). **D’.** Higher magnification of region boxed in **D. E.** *acta3:egfp* expression vascular smooth muscle cells (red) surround *kdrl:dsRed* expression blood vessels (green). **E’.** Higher magnification of the region boxed in **E. F.** *stm* (blue) expressing cells localize to the lateral margin of the ovary and colocalize with *nanos2* (green) and *dmc1* (pink) germline stem cells and early meiotic cells, respectively. **F’.** Higher magnification of the region boxed in **F.**Scale bar in C, D, and E 20*μ*m; C’, 10*μ*m, D’, E’ and F’, 5 *μ*m. IB, stage IB oocyte; II, stage II oocyte.

#### Interstitial cells (Cluster 0 and 4)

Sub-cluster 0 is the largest of the stromal cell population and likely represents the collagen-producing fibroblast-like connective tissue (Figure 6B). In addition to the expression of the collagen-encoding genes, like *col1a1a*, notable genes expressed in these cells include the *chemokine (C-X-C motif) ligand 12a* (*cxcl12a*, formerly known as *sdf1a*), the nuclear receptor *nr2f2,* the cell signaling molecule *bmp4,* and *cyp19a1a* (Figure 6A,B). Cells within sub-clusters 0 and 4 have nearly identical gene expression profiles, with the exception that sub-cluster 4 cells also have strong expression of genes associated with cell cycle progression, such as *pcna* (Figure 6A,B). Given that the ovary continues to increase in size as the fish grows, it is likely that sub-cluster 4 cells serve as a source for new interstitial cells similar to follicle cell sub-cluster 4 cells. Previous studies have shown that *cxcl12a* is essential for directing migrating primordial germ cells to the forming gonad during embryogenesis (Doitsidou et al., 2002). However, we did not detect expression of the *cxcl12a* receptor, *cxcr4a*, in any cells of the 40 dpf ovary data set, therefore the relevant receptor or functional significance of *cxcl12a* expression at this stage is not apparent. Regardless, this gene provides a convenient marker for determining the location of this cell population in the ovary. Using fluorescent *in situ* hybridization, we found that *cxcl12a* expressing cells are enriched in the interstitial regions between early follicles, but not late follicles (<Stage II; Figures 6C-C’). Together these data argue that sub-clusters 4 and 0 are proliferative and non-proliferative interstitial cells, respectively.

#### Perivascular mural cells: vascular smooth muscle (sub-cluster 1) and pericytes (sub-cluster 2)

Stromal cell Sub-clusters 1 and 2 represent the perivascular mural cells which coat the endothelial-derived blood vessels and provide stability and integrity to the vasculature. Similar to their function in establishment of the blood-brain barrier, these cells have also been implicated in formation of a blood-follicle barrier (Siu et al., 2012). Pericytes are generally solitary cells associated with small diameter blood vessels, (arterioles, capillaries, and venules) while vascular smooth muscle cells (vSMC) are associated with large blood vessels where they form a continuous coating (Gaengel et al., 2009). In zebrafish, pericytes express *platelet-derived growth factor receptor ß* (*pdgfrb*) and *notch3*, two genes whose expression are enriched in our stromal cell sub-cluster 2 (Figure 6A and Figure 6-figure supplement 2; (Wang et al., 2014). In addition, we find that sub-cluster 2 cells are enriched for *proteolipid protein 1b* (*plp1b*; Figures 6A,B), *T-box transcription factor 2* (*tbx2*), *melanoma cell adhesion molecule B* (*mcamb*), and *regulator of G protein signaling 5a* and *5b* (*rgs5a* and *rgs5b*), all of which have orthologs expressed in human ovarian pericytes (Figure 6A and Figure 6-figure supplement 2A; (Wagner et al., 2020). vSMC are characterized by expression of *α-smooth muscle actin* (*acta2*) and *transgelin* (*tagln*, formally known as SM22α; (Bahrami and Childs, 2018). We found that both *acta2* and *tagln* are enriched in stromal cell sub-cluster 1 (Figures 6A,B and Figure 6-figure supplement 2B; Table 5). In addition, we found that sub-cluster 1 has enriched expression of *myosin light chain kinase b* (*mylkb*), *tropomyosin 2* (*tpm2*) and *cysteine and glycine-rich protein 1b* (*csrp1b*), further supporting the conclusion that these cells are vascular smooth muscle (Figure 6-figure supplement 2B).

To determine the location of pericytes in the ovary, we used *plp1b* as a marker because, in contrast to *notch3* and *pdgfrb*, our data indicate that *plp1b* is specific to pericytes. We performed smFISH for *plp1b* using ovaries isolated from Tg(*fli1:egfp*) which express eGFP in vascular endothelial cells (Lawson and Weinstein, 2002). We found that *plp1b+* cells were generally solitary, but always adjacent to blood vessels, consistent with pericytes (Figures 6D-D’). To determine the localization of vSMC, we imaged ovaries from Tg(*kdrl:dsRed*); Tg(*acta2:eGFP*) double transgenic animals. Tg(*kdrl:dsRed*) is expressed in vascular endothelial cells, while *acta2:egfp* is expressed in vSMC (Kikuchi et al., 2011; Whitesell et al., 2014). We found that the majority of the *acta2+* vSMC were either adjacent to or wrapped around the *kdrl+* vascular endothelial cells (Figures 6E-E’). Together our data argue strongly that sub-cluster 2 and 1 represent pericytes and vSMC, respectively.

#### Stromal progenitor cells (sub-cluster 3)

In addition to expressing high levels of the general stromal cell genes *col1a1a* and *dcn* (Figure 6A), sub-cluster 3 cells also express many genes that are known to play roles in early mesoderm and stromal cell development (Figures 6A,B and Figure 6-figure supplement 3B,C; Table 5). For example, the transcription factors *odd-skipped related 1* (*osr1*) and *transcription factor 21* (*tcf21*; formerly *pod1*) are expressed specifically in stromal cell sub-cluster 3 cells. In mice, *Ors1* is one of the earliest genes known to be expressed in lateral plate mesoderm, derivatives of which form the kidney and gonads, and *Ors1* mutants fail to form either of these organs (So and Danielian, 1999; Wang et al., 2005). *Tcf21*-expressing cells in the mouse ovary and testis were identified through lineage labeling as multipotent mesenchymal/stromal progenitor cells that can produce all somatic cell types during development and could contribute to somatic cell turnover during aging or cell replacement following injury in the testis (Cui et al., 2004; Shen et al., 2021). Additionally, sub-cluster 3 cells and mouse *Tcf21*-expressing mesenchymal progenitors also share common expression of *osr1*, *procollagen C-endopeptidase enhancer a (pcolcea), slit homolog 3* (*slit3*), *platelet derived growth factor receptor a* (*pdgfra*), *dcn*, *col1a1a*, *col1a2,* and *insulin-like growth factor 2* (*igf2a*; Figure 6-figure supplement 3B,C; (Shen et al., 2021). We therefore propose that these cells are stromal progenitor cells. Unfortunately, we have yet to identify where these cells reside in the ovary.

#### Ovarian Cavity Epithelium (sub-cluster 5)

Stromal cell sub-cluster 5 has enriched expression of *fgf24*, a gene that is expressed in the outer epithelial layer of the early bi-potential gonad (10-12 dpf), and is required for early gonad development in zebrafish (Leerberg et al., 2017). Sub-cluster 5 cells also express the cell adhesion protein encoded by *claudin 3d* (*cldn3d;* formerly called *cldnc*) and *starmaker* (*stm*), a gene required for otolith formation in the ear (Figures 6A,B; (Söllner et al., 2003). Given the expression of *fgf24* in surface epithelial cells of the early gonad (Leerberg et al., 2017) and the role of claudin proteins in forming tight junctions between epithelial cells (Tsukita et al., 2019), it was plausible that sub-cluster 5 cells corresponded to the ovarian surface epithelium. To test this, we performed smFISH using a probe for *stm.* We chose *stm* because it is not expressed in any other cell type in the ovary and would therefore allow us to specifically identify this population of cells. We found *stm* was specifically expressed in cells that localize to the medial and lateral edges of the 40dpf ovary. a region that also contained *nanos2*+ GSCs and *dmc1*+ meiotic germ cells (Figures 6F-F’ and Figure 6-figure supplement 4). The colocalization of *stm*-expressing somatic cells with early-stage germ cells, including GSC’s raises the interesting possibility that the lateral and medial edges of the 40dpf ovary are the precursors to the germinal zones in the adult ovary.

To explore the relationship between the lateral and medial edges of the juvenile ovary and the germinal zone in the adult ovary, we performed smFISH for *stm* on transverse sections from 3-month-old adult ovaries. We found *stm-*expressing cells localize to the membrane that forms the ovarian cavity, a fluid filled space that forms on the dorsal side of the ovary and into which mature eggs are ovulated (Figure 7; (Takahashi, 1977). We therefore designate this the ovarian cavity epithelium (OCE). The OCE is contiguous with the reproductive duct located at the posterior end of the ovary and that functions as the conduit between the OCE and the genital papilla (Kossack et al., 2019). Interestingly, the OCE appears to adhere to the medial and lateral edges of the ovary at a location that correlates with that of the germinal zone (Figure 7B, C and H). To investigate this further we prepared transverse histological sections from 3-month-old adult ovaries. We found that premeiotic and early meiotic germ cells are adjacent to where the OCE adheres to the ovary at both the lateral and medial sides (Figures 7D,E and 7F,G, respectively). Finally, we found that in addition to *stm* and *fgf24*, sub-cluster 5 cells also express the *fgf24-*related ligand encoded by *fgf18b*, transcription factors *paired box 2b* (*pax2b*) and *empty spiracles homeobox 2* (*emx2*), and *wnt7aa* (Figure 7A). Orthologs of *pax2*, *emx2* and *wnt7aa* are expressed in the Müllerian ducts in mammals (Roly et al., 2020), raising the intriguing possibility that the OCE and mammalian Müllerian ducts share a common evolutionary origin. All together these data argue that *stm*-expressing cells form the ovarian cavity epithelium and may also play a role in germline stem cell niche formation and/or function.

**Figure 7.**
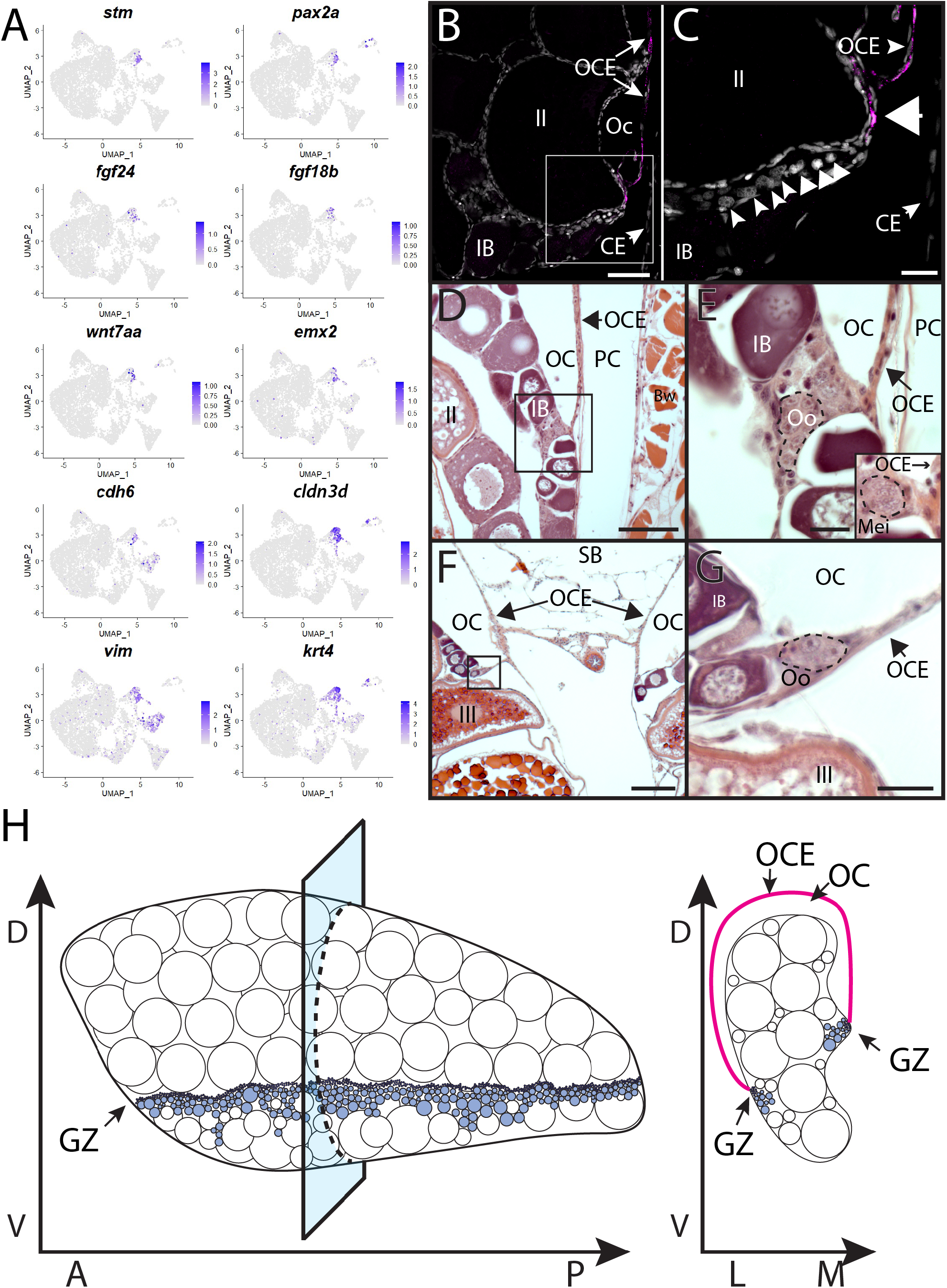
Stromal cell subcluster 3: Ovarian cavity epithelium (OCE). **A.** Gene expression UMAP plots for select genes whose expression is enriched in stromal cell subcluster 3. ligand-encoding genes. **B. s**mFISH on transverse sections from a 3-month-old ovary showing that *stm* (magenta) is expressed in the epithelium that lines the ovarian cavity. DNA is grey. **C.** Higher magnification of region boxed in **B** showing that early-stage germ cells localize to the region subjacent to where the ovarian cavity epithelium is attached to the lateral side of the ovary (arrowhead). **D-E.** Histological transverse sections from a 3-month-old ovary showing correlation between where the ovarian cavity epithelium attaches to the ovary at the lateral (**D** and **E**) and medial (**F** and **G**) sides, and the presence of pre-meiotic germ cells, characterized by large, dark staining nucleoli, and early meiotic germ cells, characterized by condensed chromosomes (inset in E). E and F are higher magnification views of regions boxed in **D** and **F**, respectively. OCE, ovarian epithelium; OC, ovarian cavity; PC, peritoneal cavity; CE, coelomic epithelium; SB, swim bladder; Oo, premeiotic oogonia; Mei, early meiotic germ cell; IB, stage IB oocyte; III, stage III oocyte. Scale bar in C, E, and G 10 *μ*m; B, 100 *μ*m; D, 200*μ*m, F, 250*μ*m.

### Theca cells and production of the 17ß-Estradiol precursor, Androstenedione

In addition to producing mature gametes, the other major role of the ovary is to produce the sex steroid hormone 17ß-Estradiol (E2) which induces female-specific sex characteristics throughout the body. E2 is the major female sex hormone in vertebrates, including zebrafish (Devlin and Nagahama, 2002). In mammals, the production of E2 requires two cell types: theca and granulosa cells (Ryan, 1979). The primary role of theca cells is to produce the intermediates for estrogen production, using cholesterol as the starting substrate and ending with the precursor androstenedione. Androstenedione is then transferred to follicle cells for conversion to E2 by two additional enzymes, most notably the Cyp19a1 aromatase (Miller and Auchus, 2011). Although many studies have investigated steroid hormone production in zebrafish, there are still significant gaps in our understanding of the biosynthetic pathway that led to E2 production. For example, while enzyme orthologs necessary for E2 production in mammals have been identified in zebrafish, the specific cell type(s) that produce E2 in zebrafish have not been determined, nor have theca cells been definitively identified.

The first, and rate limiting, step in E2 synthesis in mammals, is the transfer of cholesterol from the outer to inner mitochondrial membrane by steroidogenic acute regulatory protein, StAR, where it is converted to pregnenolone by the cytochrome P450 cholesterol side chain-cleaving enzyme, Cyp11a1 (Miller and Auchus, 2011). The expression of both *star* and *cyp11a1* have been detected in zebrafish ovaries (Bauer et al., 2000; Ings and Van Der Kraak, 2006). Unlike other vertebrates, teleost have two paralogs of *cyp11a1*, called *cyp11a1* and *cyp11a2*, with *cyp11a2* sharing the highest degree of similarity with other *Cyp11a1* orthologs (Parajes et al., 2013). Our scRNA-seq data set demonstrates that *cyp11a1* and *cyp11a2* are both expressed in a distinct population of somatic cells, thus identifying these as theca cells (Figure 1 and Figure 8-figure supplement 1). Additionally, we found that *cyp11a2* is expressed at higher levels than *cyp11a1* and therefore may encode the primary cholesterol side chain-cleaving enzyme required for E2 synthesis in the ovary (Figure 8-figure supplement 1). By contrast, the majority of *cyp11a1* expression cells in the dataset are oocytes (Stage IA; Figure 8-figure supplement 1), consistent with previous results showing that in zebrafish *cyp11a2* is maternally expressed (Parajes et al., 2013). Finally, expression of the three remaining genes encoding enzyme orthologs required for androstenedione production by theca cells, *star*, *cyp17a1* and *hydroxy-delta-5-steroid dehydrogenase, 3 beta- and steroid delta-isomerase 1* (*hsd3b1*), are also detected in theca cells (Figure 8-figure supplement 1 and Table 1). As theca cells had only been previously localized by their morphology, we performed smFISH using *cyp17a1* to determine the location of theca cells relative to oocytes. As expected, we found that *cyp17a1*-expressing cells localize to the interstitial spaces between developing oocytes (Figures 8B,C). Thus, we have molecularly characterized the theca cell population in the zebrafish ovary and confirmed that, as in mammals, theca cells are the probable source of the E2 precursor, androstenedione.

**Figure 8.**
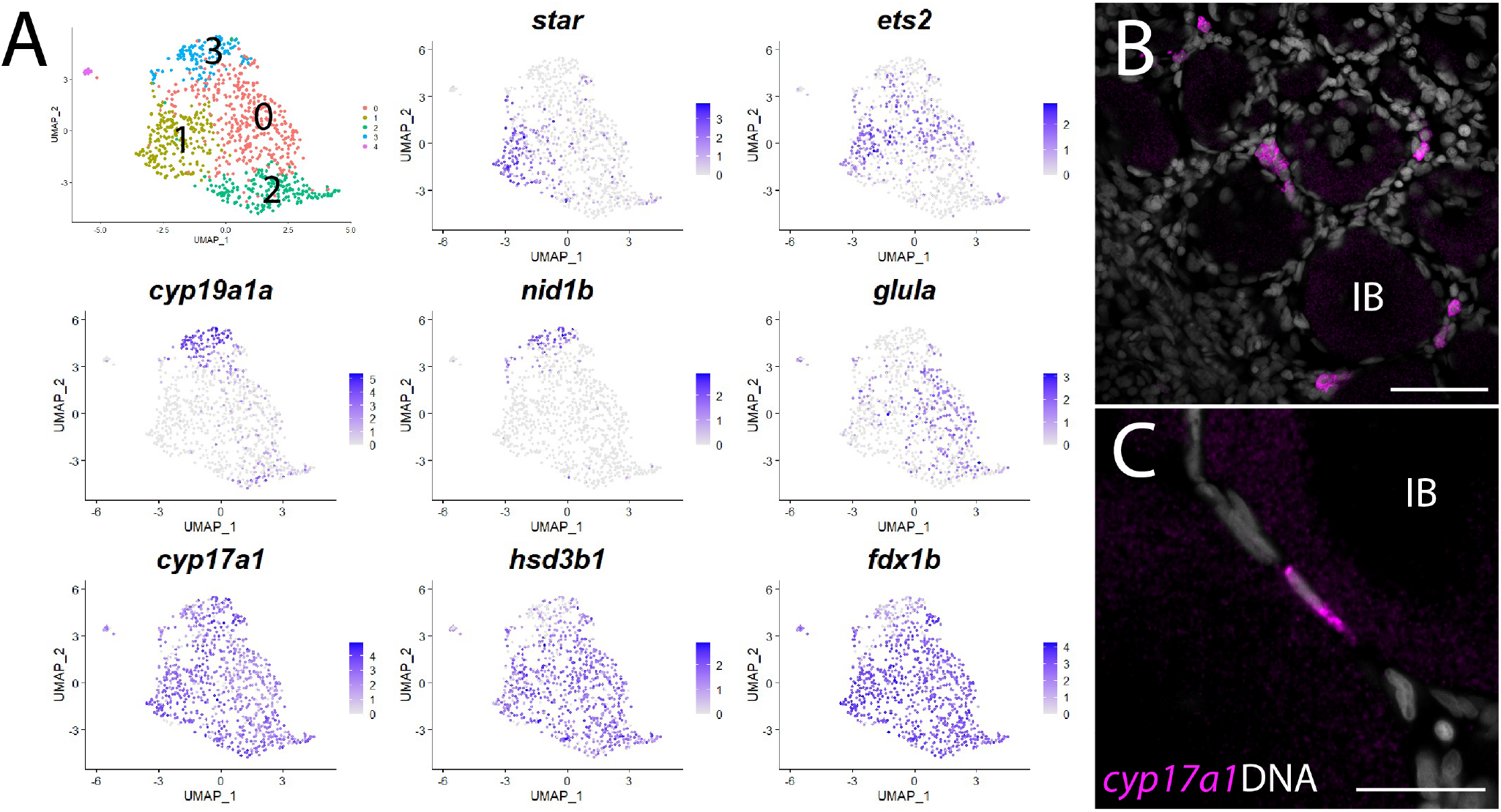
Theca cell sub-cluster analysis. **A.** Gene expression UMAP plots of select genes. Top left panel shows ells color-coded by computationally determined cell subtype. Cells expressing the indicated gene are colored purple, and the relative intensity indicates relative expression levels (intensity scale for each plot is on the right). **B.** smFISH on whole-mount 40 dpf ovaries reveals the location of *cyp17a1* expressing theca cells. DNA is gray. **C.** Higher magnification of a *cyp17a1* expressing theca cells. IB, stage IB oocyte. Scale bars in B, 50*μ*m and C, 20 *μ*m.

Further analysis of gene expression suggests that theca cells are not a homogeneous cell population (Figure 8A and Table 6). We found several genes that had non-uniform expression within theca cells in zebrafish, leading to the identification of four theca cell sub-populations (Figure 8A and Table 6). Examples of genes expressed non-uniformly in theca cells include *star* (sub-cluster 1), *cyp19a1a* and *nidogen 1b* (*nid1b*), which encodes a basement membrane-associated protein (sub-cluster 3), *v-ets avian erythroblastosis virus E26 oncogene homolog 2* (*ets2*; sub-clusters 0 and 1), and *glutamate ammonia ligase A* (*glula*), which encodes an enzyme involved in glutamine synthesis (sub-cluster 0; Figure 8A). Non-uniformity of theca cells has been previously noted in medaka based on the expression of select genes (Nakamura et al., 2009), providing support that some of these subpopulations have biological relevance that warrants further study.

### Identification of the probable pathway for estrogen production in zebrafish ovary

There are two possible biosynthetic pathways for the conversion of androstenedione to E2, which we will refer to as pathway 1 and pathway 2 (Figure 9A). Both pathways require two enzymatic steps, and the function of the Cyp19a1a aromatase. However, the pathways differ in the intermediate produced and at what step Cyp19a1a functions. Specifically, pathway 1 produces testosterone as an intermediate while pathway 2 produces estrone (E1; Figure 9A). Pathway choice is therefore not dictated solely by the expression of Cyp19a1a, but instead by the presence or absence of the two additional enzymes encoded by *17ß-hydroxysteroid dehydrogenase types 1 and 3*, called Hsd17b1 and Hsd17b3. While these enzymes catalyze similar reactions, in zebrafish Hsd17b3 prefers androstenedione as a substrate while Hsd17b1 prefers estrone (Mindnich et al., 2004; Mindnich et al., 2005); Figure 9A, Figure 9-figure supplement 1). By contrast, current evidence argues that the aromatase Cyp19a1a does not have preference for androstenedione over testosterone (Guiguen et al., 2010); Figure 9A). It therefore follows that the expression of Hsd17b1 vs. Hsd17b3 will determine pathway preference. Though it has been proposed that testosterone is the normal intermediate for E2 production in fish (Devlin and Nagahama, 2002), the pathway by which E2 is produced in the zebrafish ovary remains unknown. Thus, knowing which genes are expressed, and in what cells is key to understanding how E2 production is regulated in the zebrafish ovary.

**Figure 9.**
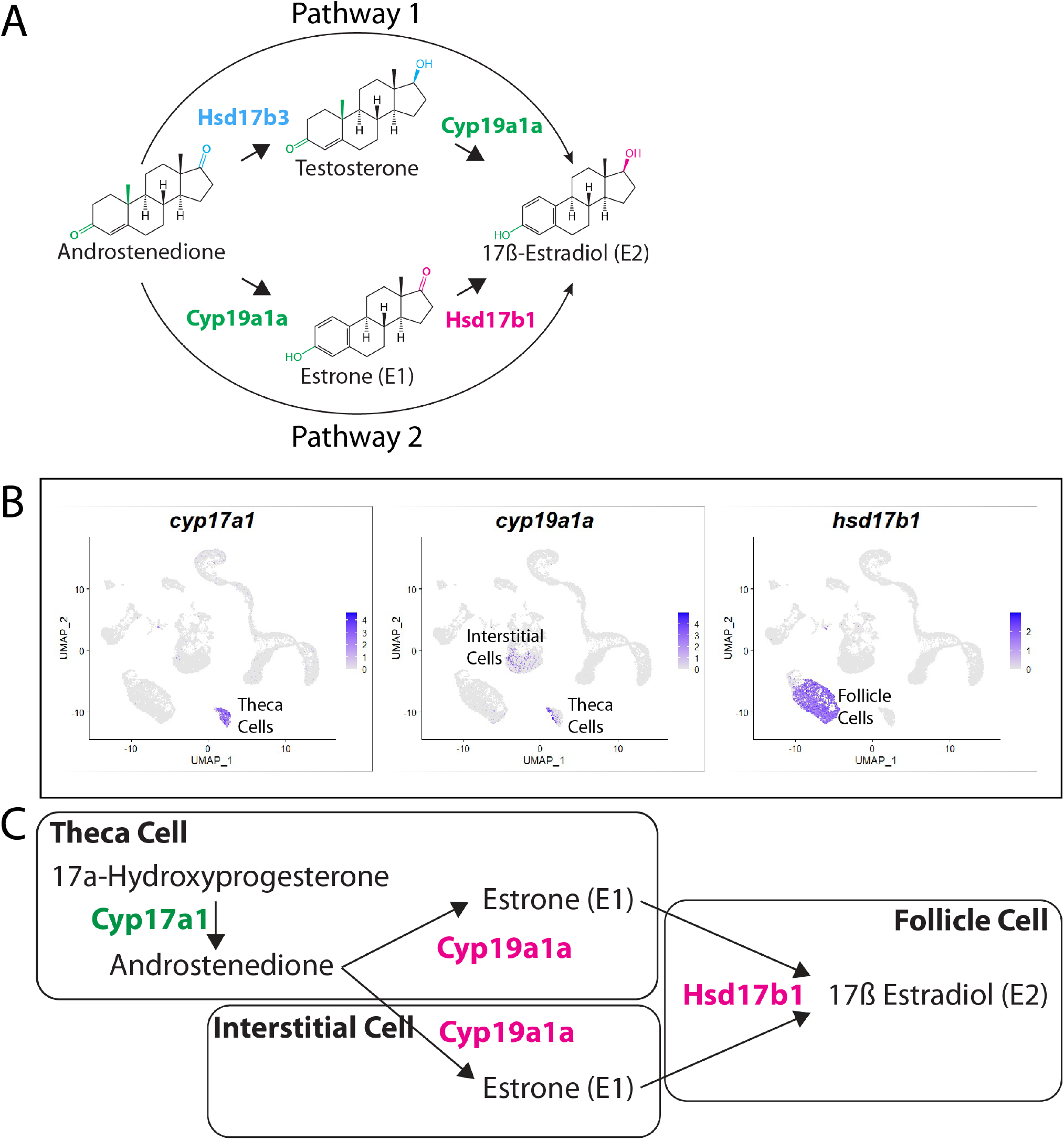
Pathway for 17ß-estradiol (E2) synthesis in the zebrafish ovary. **A.** Two possible pathways for E2 synthesis starting with androstenedione. Colors correspond to the region of the molecules being modified, and to the enzyme that catalyzes the modification. **B.** Gene expression UMAP plots of select genes. Top left panel shows cells color-coded by computationally determined cell subtype. Cells expressing the indicated gene are colored purple, and the relative intensity indicates relative expression levels (intensity scale for each plot is on the right). **C.** Proposed pathway for E2 synthesis in the zebrafish ovary, starting with the 17a-Hydroxyprogesterone intermediate precursor, together with the cell types where each reaction occurs.

Our lab previously reported that *cyp19a1a* was likely expressed in theca cells in the early ovary and then was later upregulated in follicle cells surrounding oocytes that had progressed past mid-stage II, a stage that is underrepresented in 40dpf ovaries (Dranow et al., 2016). Our scRNA-seq data verified that *cyp19a1a* is expressed in theca cells (Figure 9B) but not in early follicle cells (Figure 9B, see also Figure 4B and Figure 4-figure supplement 4). Unexpectedly we found significant expression of *cyp19a1a* in interstitial stromal cells. Thus it is likely that three district ovarian cell populations contribute to E2 production in zebrafish: theca cells, follicle cells, and stromal cells (Figure 9B,C). Finally, we determined what cell types *hsd17b1* and *hsd17b3* were expressed. We found that *hsd17b1* is expressed strongly in follicle cells while *hsd17b3* expression appears to be only weakly expressed in premeiotic germ cells (Figure 9-figure supplement 1). Three major conclusions can be drawn from this analysis: (1) The major pathway for E2 biosynthesis in the zebrafish ovary occurs via pathway 2, (2) Hsd17b1, not Cyp19a1a, catalyzes the final enzymatic step in E2 synthesis, and (3) follicle cells are the major source of E2 production in the zebrafish ovary (Figure 9).

## DISCUSSION

We have presented here the transcriptome of the 40 dpf zebrafish ovary at single cell resolution, analysis of which has allowed us to define the major cell types present in the juvenile ovary. Further sub-cluster analysis, supported by molecular and mutational evidence, revealed an unexpected level of cell subtype heterogeneity within the somatic cells that likely play important roles in regulating germ cell development as well as the development of female sexual characteristics through the production of estrogen. Our data provides strong support that orthologs of genes involved in mammalian ovary development and function likely play conserved roles in zebrafish, further validating zebrafish as a relevant model for understanding vertebrate ovarian development and function. Similarly, the major cell types required for mammalian ovary development and function have clearly identifiable homologs in the zebrafish ovary. However, unlike mammals, zebrafish females can produce new follicles throughout adult life from a population of self-renewing GSCs and pre-follicle progenitor cells, thus allowing the study of these unique cell types in a tractable vertebrate research animal. Importantly, this reference data set will facilitate the use of zebrafish to study female reproductive diseases and disorders of sexual development.

### High resolution transcriptomics of pre-meiotic, meiotic and early-stage oocytes

We transcriptionally profiled over 10,000 germ cells that represent all developmental stages from germline stem cells to early-stage oocytes (Stage IA). Though previous work had identified genes expressed in most of these developmental stages (e.g. *nanos2* in GSCs and *dmc1* in early meiosis; Beer and Draper, 2013), our data set allowed us to identify markers for oocyte progenitor cells and cells that are undergoing pre-meiotic S-phase (*foxl2l* and *rec8a*, respectively). The ability to identify specific germ cell stages in the ovaries will allow us to more accurately define the structure and function of the GSC niche and to better characterize germline developmental defects that result from experimental manipulations (e.g. mutant analysis or chemical perturbation). In addition to identifying new stage-specific markers, the sequencing depth has allowed us to also identify gene modules that may represent co-regulated genes (e.g. novel markers of GSCs), as well as candidate transcriptional regulators. This information will allow us to build more accurate gene regulatory networks between zebrafish and humans, which can be experimentally tested in the genetically tractable zebrafish. Finally, we provided genetic evidence that the oocyte progenitor cell-expressed transcription factor *foxl2l* is likely required for oocyte development in zebrafish, similar to the role of the *foxl2l* ortholog in medaka (Nishimura et al., 2015). It is well established in zebrafish that animals that cannot produce oocytes invariably develop as males (Anderson et al., 2012; Rodriguez-Mari et al., 2010; Shive et al., 2010) and we found that all *foxl2l* mutants developed as males, consistent with a role for *foxl2l* in oocyte development.

### Follicle cells and pre-follicle cells

We identified three distinct populations of follicle cells. Two of these were populations of differentiated follicle cells that surrounded Stage 1B and early-Stage II oocytes. This is consistent with the 40 dpf ovaries we used for the analysis, as they contain few oocytes that are more advanced than early stage II. The third population we identified are likely pre-follicle cells. Coupled with the ability to produce new oocytes for GSCs is the need to produce new follicle cells to support them. In mammals, new follicle cells are produced from a transient population of pre-follicle cells only during embryogenesis (McLaren, 1991), however given the presence of GSCs in the adult zebrafish ovary we expected zebrafish to maintain a population of pre-follicle cells throughout life. Indeed, we identified a subpopulation of follicle cells that expressed both the canonical teleost follicle cell marker *gsdf* and transcription factor *lhx9*. Our analysis argues strongly that these cells are pre-follicle cells. First, *lhx9*-expressing cells do not localize around oocytes, but instead associate with pre-follicle stage germ cells. Second, these cells uniquely expressed *irx3a* and *irx5a*, homologs of which are expressed in, and required for, mammalian pre-follicle cells (Fu et al., 2018). Third, these cells express orthologs of genes associated with early-stage somatic gonad development in mammals, such as *wt1a* and *emx2* (Kreidberg et al., 1993; Miyamoto et al., 1997) but do not express high levels of genes that are associated with follicle cell function, such as *wnt4a* and *amh* (Kossack et al., 2019; Yan et al., 2019). Finally, a subset of these cells express *sox9b*, a gene previously shown to be expressed in pre-follicle in the medaka ovary (Nakamura et al., 2008).

The pre-follicle cell sub-cluster appears to contain two cell subtypes that represent early (sub-cluster 3.1) and transitional (sub-cluster 3.2) developmental stages, as suggested by their clustering position relative to the differentiated follicle cells (sub-cluster 1). Both subtypes appear to express equivalent levels of *irx3a* and *irx5a*, establishing them as per-follicle cells, but sub-cluster 3.1 expresses higher levels of both *lhx9* and *wt1a* while the sub-cluster 3.2 expresses higher levels of *gsdf*. It is interesting that the *lhx9*-high sub-population also expresses higher levels of genes encoding cell signaling attenuators, the Bmp antagonists Nog1 and Nog3, the Wnt antagonist Dkk3b and the retinoic acid degrading enzyme Cyp26a1, which suggest these cells create a local microenvironment that is devoid of these signals. It is possible that inhibition of Bmp, Wnt, and retinoic acid signaling is necessary to keep pre-follicle cells in an undifferentiated, stem cell-like state. Alternatively, these cells could produce a microenvironment that influences other surrounding cells, such as GSC. In this regard, *sox9b*-expressing pre-follicle cells in medaka have been proposed to function as the GSC niche (Nakamura et al., 2010). By contrast, the *gsdf*-high expression subpopulation expresses genes encoding the Bmp ligand Bmp6 and the Fgf receptors Fgfr3 and Fgfr4. These *gsdf*-high cells appear to be responding to Fgf signaling as they express *spry4* and *etv4*, genes known to be upregulated in response to Fgf signaling (Fürthauer et al., 2001; Raible and Brand, 2001; Roehl and Nüsslein-Volhard, 2001). We did not detect significant expression of Fgf ligands in any of the follicle cell subpopulations profiled suggesting that the source of Fgf is a non-follicle cell subtype. Interestingly, we had previously published a role for the Fgf ligand encoded by *fgf24* in the development of the bipotential somatic gonad in zebrafish. We found that *fgf24*-expressing surface epithelium in the 10 dpf gonad promoted *etv4* expression in the underlying mesenchymal cells but was not required for the expression of *wt1a* (Leerberg et al., 2017). This raises the possibility that these two cell subpopulations are present in the early bipotential gonad and that Fgf signaling is important for pre-follicle cells to transition to differentiated follicle cells.

### Stromal cell populations

The ovarian stromal cells in zebrafish were arguably the least characterized cell type in the zebrafish ovary. Our analysis identified five distinct stromal cell subtypes present in the 40 dpf ovary: interstitial cells, vascular smooth muscle cells, pericytes, ovarian cavity epithelial cells, and probable stromal progenitor cells. The interstitial cells likely play a structural role, consistent with their expression of many collagen-encoding genes. We also noted that expression of the cytokine ligand *cxcl12* is enriched in the interstitial cells, this ligand plays an essential role in guiding germ cell migration to the gonad during embryogenesis (Doitsidou et al., 2002). We did not, however, detect significant expression of the Cxcl12 receptor, encoded by *cxcr4a,* in any of the 40 dpf germ cells (Knaut et al., 2003). Instead, we found that vascular endothelial cells express *cxcr4a* (Supplemental Table 1) raising the possibility that Cxcl12/Cxcr4a signaling axis may instead be involved in regulating ovarian angiogenesis (Petit et al., 2007).

The second interesting stromal cell type we identified express *stm* and that form the ovarian cavity epithelium (OCE). Teleost ovaries can be categorized as two general types-gymnovarian and cystovarian. Gymnovarian ovaries ovulate mature eggs directly into the coelomic cavity whereas cystovarians, like those of zebrafish, ovulate eggs into a fluid filled ovarian cavity that forms between the dorsal side of the ovary and the OCE (Takahashi, 1977). The OCE is contiguous with the reproductive duct at the posterior end of the ovary (Kossack et al., 2019). The presence or absence of an OCE is therefore a defining feature of ovarian function, yet little is known about the development of the OCE in any teleost. Our analysis has identified *stm* as a specific marker of the OCE. *stm* encodes a protein that is predicted to be intrinsically disordered and that is a probable homolog of human Dentin Sialophosphoprotein (DSPP)(Söllner et al., 2003). Mutational analysis in zebrafish has shown it is involved in regulating the formation of otoliths in the ear, which are composed of crystallized calcium carbonate and function in balance and hearing (Söllner et al., 2003). It is not clear what role *stm* may play in the OCE development or function. Studies in medaka have shown that the transcription factor Pax2 is a probable direct regulator of the *stm* ortholog, *starmaker-like* (Bajoghli et al., 2009). We found that *pax2a* expression is nearly identical to that of *stm* in these stromal cells, raising the possibility that Pax2a may also regulate the expression of *stm* in OCE cells.

The OCE attaches to the lateral and medial sides of the adult ovary precisely where the germinal zone and GSC niche is located. It is an exciting possibility that this asymmetry on the surface of the ovary plays an important role, either directly or indirectly, in the formation and/or function of the ovarian GSC niche in zebrafish. In addition to *stm*, the OCE also expresses *fgf24,* a gene we have previously shown was expressed in the surface epithelium of the bipotential gland and was required for the development of the early somatic gonad cells, as well as its close homolog *fgf18b* (Jovelin et al., 2007; Leerberg et al., 2017). Though *stm*-expressing cells may not play a direct role in niche function, the identification of any cell type that localizes near the niche is nevertheless an important element towards defining the niche environment.

Finally, we note that the OCE expresses orthologs of several genes that are known to be expressed in the developing Müllerian ducts of mammals and birds. These include *wnt7aa*, one of two zebrafish orthologs of mammalian *Wnt7a*, *pax2a* and *emx2* (Mullen and Behringer, 2014; Roly et al., 2020). We had argued previously that the reproductive duct, which connects the ovarian cavity to the genital papilla in zebrafish, may have a common developmental origin with Müllerian ducts in mammals as these structures fail to form in both mammalian *Wnt4* and zebrafish *wnt4a* mutants (Kossack et al., 2019; Vainio et al., 1999). However, we also noted that formation of the ovarian cavity was normal in *wnt4a* mutants (Kossack et al., 2019). Regardless, these findings raise the intriguing possibility that the OCE and the reproductive ducts in fish may have a common developmental origin to Müllerian ducts in mammals.

### Stromal progenitor cells

The third stromal cell subpopulations that warrants discussion are those contained within subpopulation 3 and that express *osr1* and *tcf21*. In mice *Osr1* is the earliest marker of lateral plate mesoderm, which is the embryonic territory from which the urogenital system is derived (So and Danielian, 1999). *Tcf21*-expressing cells have been shown via lineage tracing to produce all known somatic cell types of the ovary and testis during development or in the adult following wounding, suggesting that this cell population has stem cell-like properties (Shen et al., 2021). We do not yet know if these cells are a stable population that persist into adulthood or if instead, they are a transient population present only during the juvenile and early adult stages while the ovary is increasing in size. It will be important to determine when these cells are first detected during gonad development, how long they persist, and whether they can function as precursors for multiple somatic gonadal cell lineages.

### Role of Wnt9b and Wnt4a in female sex determination and/or differentiation

The mechanisms of sex determination in zebrafish is not well understood, as domesticated zebrafish do not have sex chromosomes, and the sex determining gene that appears to exist in wild populations has yet to be identified (Wilson et al., 2014). Regardless, recent work has shown that orthologs of key genes involved in mammalian sex differentiation play conserved roles in zebrafish, arguing that the general pathway for sex differentiation in vertebrates is conserved (e.g. Webster (Webster et al., 2017; Yang et al., 2017). Our lab has previously published that loss of *wnt4a* resulted in female-to-male sex reversal, as it does in mammals, though the phenotype was only 95% penetrant (Kossack et al., 2019). Our scRNA-seq data allowed us to identify *wnt9b* as an additional Wnt ligand-encoding gene that is co-expressed with *wnt4a* in follicle cells. Similar to *wnt4a*, *wnt9b* mutants have an incompletely penetrant female-to-male sex reversal phenotype. If Wnt4a and Wnt9b function in parallel, it is not clear why loss of one ligand would have such a strong but incompletely penetrant phenotype. However, our observation that some *wnt9b* mutants appear to initiate female development before sex reverting to males may indicate that the primary role of Wnt9b is in maintenance of female sex determination, while Wtn4a may function primarily during sex determination. Resolving this question will require a more detailed analysis of the *wnt9b* phenotype. Interestingly, Wnt9b has previously been shown to be expressed in trout follicle cells, suggesting that it may have a similar function in sex determination and/or differentiation in other teleost (Nicol and Guiguen, 2011). Finally, Wnt9b in mammals is not expressed in granulosa cells and there is no evidence that it plays a role in sex determination or differentiation. Instead, Wnt9b is involved in formation of the male and female reproductive ducts, the Wolffian and Müllerian ducts, respectively (Carroll et al., 2005). Our preliminary analysis of *wnt9b* mutant zebrafish indicates that the reproductive ducts in males are functional, suggesting that in zebrafish *wnt9b* is not required for reproductive duct formation.

### Defining the pathway for estrogen synthesis in the juvenile zebrafish ovary

A key downstream component of any sex determination and differentiation pathway is the production of the appropriate sex hormone by the gonad. For female vertebrates, 17ß-estrodiol (E2) is the major bioactive sex hormone and in zebrafish and is essential for female development (Devlin and Nagahama, 2002; Dranow et al., 2016). However, there are significant gaps in our knowledge about the regulation of E2 synthesis in zebrafish. In mammals, E2 production requires both granulosa cells and theca cells. Theca cells produce the precursor androstenedione, which is then converted to E2 in granulosa cells. A key granulosa cell-produced enzyme required for the conversion of androstenedione to E2 is aromatase encoded by the *Cyp19A1* gene. The zebrafish genome encodes two *Cyp19a1* orthologs, called *cyp19a1a* and *cyp19a1b*, though only *cyp19a1a* is expressed in the ovary. We had previously shown that during the period of sex determination through early ovary development (∼10-40dpf) *cyp19a1a* expression was not detected in follicle cells, as would be expected based on the expression in mammals, but was instead expressed in theca cells (Dranow et al., 2016). Our current scRNA-seq data confirms our previous results and additionally shows that *cyp19a1a* is also expressed in interstitial cells. This expression raised the possibility that cells other than follicle cells were the source of E2 production during the period of sex determination and early ovary development. However, there are two possible pathways for E2 synthesis from androstenedione, and while both require aromatase, its position within each pathway differs (Tokarz et al., 2013). In the first pathway, androstenedione is converted to testosterone by Hsd17b3, which is then converted to E2 by Aromatase (Mindnich et al., 2005). By contrast, aromatase can also convert androstenedione to estrone (E1), which can then be converted to E2 by Hsd17b1 (Mindnich et al., 2004). The key to determining the pathway used in the early ovary therefore lies in determining if and where Hsd17b1 and Hsd17b3 are produced. Our scRNA-seq data revealed that expression of *hsd17b1*, but not *hsd17b3*, is detected only in the follicle cells of the 40 dpf ovary. This leads to two conclusions. First, E2 in the 40dpf ovary is produced from an E1 intermediate, not from testosterone. Second, because *hsd17b1* is the last enzyme in this pathway and its expression is limited to follicle cells, E2 is produced by only follicle cells in the zebrafish ovary, as it is in mammals. Thus, *hsd17b1* should be considered as a candidate regulatory node for E2 production when considering pathways for sex determination, differentiation and maintenance.

## CONCLUSIONS

Our single cell analysis has identified the major cell populations in the juvenile zebrafish ovary. Using smFISH, we have validated the data set, thus providing confidence that it will serve as a valuable resource to the community going forward. We also provided several examples to demonstrate the utility of these data to quickly identify relevant candidate genes for further analysis. First, we identified *foxl2l* as a gene specifically expressed in oocyte progenitor cells and, using mutational analysis, demonstrated that it was necessary for oocyte production. Second, we identified *wnt9b* as an additional Wnt ligand-encoding gene that promotes female sex differentiation, a hypothesis that we validated using mutational analysis. Through defining the various somatic cell subtypes and the identification of marker genes that allowed their location in the ovary to be determined, we have identified candidate cell populations that could function as the niche for germline stem cell maintenance. Finally, through analysis of genes that encode the enzymes necessary for estrogen production, we have identified the most likely pathway for estrogen synthesis in the zebrafish ovary, and the specific somatic cells that are likely to be involved.

## DATA AVAILIBILITY

The raw data reported in this paper are archived at NCBI GEO (accession number TBD) and in processed and interactively browsable forms in the Broad Single-Cell Portal (https://singlecell.broadinstitute.org/single_cell/study/SCP928/). Analysis code and objects are archived at github (https://github.com/yulongliu68/zeb_ov_ssRNAseq). Gene expression tables for the cell clusters identified are archived at Dryad: (https://datadryad.org/stash/share/CEd0Zs4oZKdinTWeJPKbWYjBq6hYq4QhVacQcFjf37E)

## ACKNOWLEDGMENTS

The authors wish to thank Jack Cazet for his help with NMF analysis; Members of the Juliano and Burgess labs, Blanche Capel and Jennifer McKey for thoughtful discussions; Florence Marlow and Blanche Capel for critical reading of the manuscript; Jeffery Essner for GeneWeld plasmids; UC Davis Genome Center Sequencing Core; MCB Light Microscope Facility

## Funding

R01 HD-081551 and NSF/IOS-1456737 to B.W.D.; R35 GM133689 to C.E.J.; I.A.D. and C.N.K. were supported by NIH grants DK126021 and DK107372 to I.A.D.; M.E.F and S.R.W. were supported in part by the NIH T32 predoctoral training program in Molecular and Cellular Biology (GM007377); M.E.K was supported in part by the NIH T32 predoctoral training program in Environmental Health Science (ES-0070599); S.R.W was supported in part by the NSF graduate research fellowship program (2036201); Y.L was supported in part by the UC Davis Dissertation Year Fellowship.

## Author contributions

Y.L. and B.W.D conceived the study; Y.L. and B.W.D. wrote the paper with revisions by M.E.K, M.E.M., S.S., I.D., C.K. and C.E.J.; Y.L., M.E.K, M.E.M and B.W.D. collected single-cell transcriptomes; Y.L., M.E.M., S.H. and N.A. performed ISH validation experiments; M.E.K. performed transgenic validation experiments; Y.L., M.E.K., S.R.W., and B.W.D. performed imaging; M.E.M., L.C., S.R.W. and C.K. performed gene mutant generation and analysis; Y.L. performed regulatory module analysis.

## Competing interests

The authors declare no competing interests.

## METHODS

### Animals

Zebrafish used for scRNAseq libraries are Tg(*piwil1:egfp*)*^uc02^* transgenics (formerly *ziwi:egfp*; Leu and Draper, 2010) in the AB strain. Dissected ovaries for whole-mount immunochemistry, RNA *in situ* hybridization, FISH, and smRNA FISH were from the wild-type AB strain. Zebrafish husbandry was performed as previously described (Westerfield, 2000). The following transgenic lines were also used: Tg(*fli1:egfp*)*^y1^*, (Lawson and Weinstein, 2002); Tg(*kdrl:dsRed*)*^pd27^*,(Kikuchi et al., 2011); Tg(*acta2:eGFP*)*^ca7^*, (Whitesell et al., 2014). All animal work was carried out with approval of the UC Davis IACUC.

### Zebrafish ovary dissociation for single-cell RNA sequencing

#### Germ cell library cell dissociation

Germ cells were dissociated from the somatic gonad using a modification of Blokhina et al., (2019). 40 pairs of 40 dpf ovaries were dissected from Tg(*piwil1:egfp*)*^uc02^* transgenic fish (Leu and Draper, 2010) and stored in a LoBind tube (Cat.No. 0030108302; Eppendorf) containing 2 mL of L15 medium(Cat.No. L5520; Sigma-Aldrich). The tissue was minced with small scissors into <1 mm pieces. 200 *μ*L of 20 mg/mL of type 2 collagenase in L15 (Cat.No. NC9870009; Worthington) were added and incubated on an orbital rotator at 28°C for 35 min. The cell suspension was then gently passed through a 23g needle five times to break up large cell clumps. 200 *μ*L of 7 mg/mL trypsin (Cat.No. LS003708; Worthington) in L15 were added and incubated on an orbital rotator for 10 min or until a minimal amount of cell clumps was observed. The trypsin reaction was stopped by adding 500 *μ*L of 20 mg/mL trypsin Inhibitor (Cat.No. 100612; MP Biomedicals) in L15. The cells were centrifuge for 3 min at 300 x g and the supernatant carefully removed. The cell pellet was then resuspended and washed two times with 5 mL of L15 using a P1000 pipette tip and subsequently centrifuged for 3 min at 300 x g. Cells were resuspended in 1 mL L15 and filtered first through a 100 *μ*m nylon filter (Cat.No. 431752; Corning) and then through a 40 *μ*m nylon filter (Cat.No. 431750; Corning) to remove cell clumps. The filtrate was centrifuged for 3 min at 300 x g and resuspended in 1 mL of 50 mg/mL BSA (Cat.No. A8806; Sigma-Aldrich) in phosphate buffered saline (PBS). Cell viability and number were determined using propidium iodine (Cat. No. P1304MP; Thermo Fisher) and Hoechst 33342 (Cat.No. H3570; Thermo Fisher) staining on a Fuchs-Rosenthal hemocytometer (Cat.No. DHC-F01; Incyto).

#### Whole ovary library cell dissociation

Somatic ovary cells were dissociated using a modification of Elkouby and Mullins (2017). 40 pairs of 40 dpf ovaries were dissected from Tg(*piwil1:egfp*) transgenic fish (Leu and Draper, 2010) were dissected and stored in a LoBind tube containing 2mL of L15. Tissues were minced with microdissection scissors into < 1 mm pieces. After the tissue had settled to the bottom of the tube, the media was replaced with 5mL of digestive enzyme mixture (3 mg/mL collagenase I (Cat.No. C0130; Sigma-Aldrich), 3 mg/mL collagenase II (Cat.No. C6885; Sigma-Aldrich), and 1.6 mg/mL hyaluronidase (Cat.No. H4272; Sigma-Aldrich) in L15 and incubated on an orbital rotator at room temperature. The suspension was monitored every 10 mins until no or a minimal number of cell clumps were observed (∼30 min.). Cells were centrifuged for 3 mins at 300 x g and resuspended in 5 mL of 5x TrypLE (Cat.No. A1217701; Thermo Fisher) in L15 and incubated on an orbital rotator at room temperature for 15 min. The trypsin reaction was stopped by adding 500 *μ*L of 2.8 mg/mL trypsin inhibitor in L15 and incubated on an orbital rotator at room temperature for 1 min. The cell suspension was then added to 25 ml of L15 in a 50 ml conical tube to dilute the trypsin and centrifuged for 3 min at 300 x g. The cell pellet was resuspended and washed two times with 5 mL of L15 using a P1000 pipette and centrifuged for 3 mins at 300 x g. The cell pellet was then resuspended and filtered as described above.

### Isolation of germ cells by Fluorescence-Activated Cell Sorting (FACS)

Following cell dissociation of Tg(*piwil1*:egfp) transgenic ovaries, GFP+ germ cells were sorted using a MoFlo Astrios EQ Cell Sorter (Beckman Coulter) with a 70 *μ*m nozzle. Cells were sorted using side scatter and GFP purify to identify single germ cells.

### Single-cell RNA library preparation and sequencing

Single-cell RNA sequencing libraries were prepared by the UC Davis DNA Technologies core. Briefly, barcoded 3’ single-cell libraries were prepared from dissociated cell suspensions or sorted cells using the Chromium Single-Cell 3’ Library and Gel Bead kit V3 (10X Genomics) for sequencing according to the manufacturer’s recommendations. All libraries were targeted at 10,000 cell recovery and were amplified using 11 cycles. The cDNA and library fragment size distribution were verified via micro-capillary gel electrophoresis on a Bioanalyzer 2100 (Agilent). The libraries were quantified by fluorometry on a Qubit instrument (LifeTechnologies) and by qPCR with a Kapa Library Quant kit (Kapa Biosystems) prior to sequencing. Libraries were sequenced on a HiSeq 4000 sequencer (Illumina) with paired-end 100 bp reads.

### Cell Ranger genome reference and gene annotation file generation

A General Transfer Format (GTF) gene annotation file (release 96) for the GRCz11 zebrafish genome was downloaded from Ensembl Genome Browser and filtered using the “mkgtf” function in Cell Ranger (v3.0.2; 10x Genomic) to retain the following attributes: protein_coding, lincRNA, and antisense. A genome reference file was generated with Cell Ranger’s “mkref” function using the GRCz11 zebrafish genome obtained from the Ensembl Genome Browser with alternative loci scaffolds removed and the filtered GTF file.

### Count file generation

GRCz11 zebrafish genome FASTQ file from Ensembl Genome Browser with alternative loci scaffolds removed and the filtered GTF file described above were used to generate the count file using the “count” function in Cell Ranger (v3.0.2; 10x Genomic). “expect-cells” was set to 10000 based on estimated cell recovery.

### Cell cluster analysis

Analysis scripts and data for the cell cluster analysis are available on GitHub(https://github.com/yulongliu68/zeb_ov_ssRNAseq), and the final clustering can be explored online at the Single Cell Portal (The Broad Institute of MIT and Harvard, https://singlecell.broadinstitute.org/single_cell/study/SCP928). Briefly, the expression matrices generated from Cell Ranger (germ cell library zx1_40gc, whole ovary libraries zx2_40ov and zx4_40ov) were first processed with SoupX (v0.3.1; (Young and Behjati, 2020)) to remove ambient RNA. During dissociation and library generation, lysed cells can lead to ambient RNA levels that interfere with subsequent analyses. Despite several washing steps post dissociation, the initial data exploration revealed the presence of oocyte RNA across a variety of cell types. SoupX is designed to identify and remove ambient RNA contaminations. We used non-oocyte cells and inferred top oocyte-specific genes to quantify the extent of the contamination.

The adjusted datasets were then processed with DoubletFinder (v2.0.2; (McGinnis et al., 2019)) to determine and remove doublets that are expected in any large-scale single-cell RNA sequencing datasets. DoubletFinder uses cell expression proximity and artificially generated doublets to determine potential doublets. Prior to assessing doublets, we performed quality control on the dataset. For the sorted germ cell library, we retained cells with 200 to 6000 genes, less than 150,000 unique transcripts, and less than 5% mitochondrial transcripts. For whole ovary libraries, we retained the cells with 200 to 8000 genes, less than 200,000 unique transcripts, and less than 20% mitochondrial transcripts. These cutoffs were determined by the gene and transcript distributions of those libraries. Blood cells and a small number of germ cells were also removed from the whole ovary libraries. DoubletFinder was run using a conservative 5% estimated cell doublet cutoff (Figure1-figure supplement 1).

Clustering analyses were conducted using Seurat (v3.1.0; (Stuart et al., 2019)) and the “SCTransform” workflow to normalize, identify variable genes, and perform scaling of the data. Principal component analysis (PCA) was performed using the top 3000 variable genes, and principal components considered were chosen based on the standard deviation of the elbow plot, p-value of the jackstraw plot, and biological knowledge of the genes in individual components. UMAP analysis and plots are generated based on selected principal components. Gene expression tables for all cells, germ cell subset, pre-follicle cell subset, follicle cell subset, stromal cell subset and theca cell subset are available for downloaded at Dryad: (https://datadryad.org/stash/share/CEd0Zs4oZKdinTWeJPKbWYjBq6hYq4QhVacQcFjf37E)

### Trajectory analysis

Monocle3 (v0.1.3) was used for trajectory analysis (Cao et al., 2019a). The expression matrix was exported from the Seurat object and used as Monocle3 input. UMAP was used for dimensionality reduction. We used germline stem cell (GSC) specific *nanos2* expression to identify the root of the trajectory and measured pseudo-time.

### Gene module analysis and motif enrichment

Non-negative matrix factorization (NMF) as a dimensionality reduction strategy was used to identify groups of co-expressed genes as previously described (Brunet et al., 2004; Farrell et al., 2018; Siebert et al., 2019). Expression data for the top 3,000 variable genes as identified in the Seurat analyses were used as input to the NMF analysis. To identify the optimal number of gene modules that can describe the dataset we tested a broad range of K values from 10 to 100 in increments of 5 and then reduced to an increment of 1 in a range between 30-40. The optimal K was identified as 36 for the germ cell dataset. Identified gene modules were then filtered to remove low-quality modules if a reproducibility score was lower than 0.6 or the gene module consisted of less than 10 genes.

For motif enrichment within 5’ regions of co-expressed genes, we extracted 2kb upstream of the transcription start site of the top 20% of the genes within a module based on gene scores. Gene score is a metric describing how well a particular gene reflects the expression of the associated gene module. All sequences were from Ensembl and extracted using biomaRt (v2.40.4; (Cunningham et al., 2019)). We used MEME (v5.1.0; (Bailey et al., 2015; McLeay and Bailey, 2010) for motif enrichment analysis with the following parameter: “--scoring avg --method fisher --hit-lo-fraction 0.25 --evalue-report-threshold 10.0 --control --shuffle-- --kmer 2 sequence”, and the following databases: jolma2013.meme, JASPAR2018_CORE_vertebrates_non-redundant.meme, and uniprobe_mouse.meme.

### Gene ontology analysis

The top 25% of the cluster genes (sorted by p-value) were used for GO analysis inputs. Go enrichment was assessed using g:Profiler2 (v0.2.0; (Raudvere et al., 2019)). The zebrafish biological process database was used, and enriched GO terms with the significance of <0.05 were considered.

### RNA *in situ* hybridization

All dissected ovary samples were fixed with 4% paraformaldehyde overnight at 4°C. Samples were dehydrated with 100% methanol for 10 mins at room temperature and stored at −20°C at a minimum overnight with fresh 100% methanol.

HCR fluorescence RNA *in situ* hybridization probes were ordered from Molecular Instruments. Accession numbers supplied to the manufacturer for designing of the probes, and the lot numbers for reordering are listed in table below. Hybridization procedures followed previous instructions and the official Molecular Instruments protocol (MI-Protocol-HCRv3-Zebrafish version 5; Molecular Instruments) with the following modification: tissue preparation followed the procedure above, tissues were permeabilized with proteinase K at 50 *μ*g/mL for 15 minutes, samples were cleared before mounting with 30%, 50%, and 70% glycerol-PBS gradient for 1 hour in each step, and they were mounted with ProLong™ Diamond Antifade Mountant (Cat.No P36961, Invitrogen)

Probes used for HCR fluorescence RNA *in situ* hybridization

| Target gene | Organism | Lot Number | Order Number | Amplifier |
| --- | --- | --- | --- | --- |
| *nanos2* | Zebrafish (*Danio rerio*) | PRB905 | d6f5fd8c-b888-4bf7-b9a1-d6d95db66d24 | B2 |
| *chgb* | Zebrafish (*Danio rerio*) | PRB908 | d6f5fd8c-b888-4bf7-b9a1-d6d95db66d24 | B1 |
| *id4* | Zebrafish (*Danio rerio*) | PRB909 | d6f5fd8c-b888-4bf7-b9a1-d6d95db66d24 | B1 |
| *foxl2l* (zgc:19418) | Zebrafish (*Danio rerio*) | PRD331 | 165f08fe-24a8-40e3-acb3-02767bf38a37 | B1 |
| *lhx9* | Zebrafish (*Danio rerio*) | PRD333 | 165f08fe-24a8-40e3-acb3-02767bf38a37 | B3 |
| *dmc1* | Zebrafish (*Danio rerio*) | PRD334 | 165f08fe-24a8-40e3-acb3-02767bf38a37 | B1 |
| *rec8a* | Zebrafish (*Danio rerio*) | PRE326 | 20502418-92db-49c1-801c-fc808aa3940f | B3 |
| *bhlhe23* | Zebrafish (*Danio rerio*) | PRE327 | 20502418-92db-49c1-801c-fc808aa3940f | B1 |
| *gpc1a* | Zebrafish (*Danio rerio*) | PRF573 | 984cd5dd-9cdc-4496-94eb-c722f6c027ae | B3 |
| *plp1b* | Zebrafish (*Danio rerio*) | PRF574 | 984cd5dd-9cdc-4496-94eb-c722f6c027ae | B3 |
| *cyp17a1* | Zebrafish (*Danio rerio*) | PRF576 | 984cd5dd-9cdc-4496-94eb-c722f6c027ae | B3 |
| *stm* | Zebrafish (*Danio rerio*) | PRF578 | 984cd5dd-9cdc-4496-94eb-c722f6c027ae | B3 |
| *bmp6* | Zebrafish (*Danio rerio*) | PRG855 | db0a04b4-ef3d-4944-9061-690ec2d1c956 | B3 |

RNA in situ hybridization was performed essentially as described (Thisse and Thisse, 2008). Template DNA for production of a *gja11* RNA *in situ* probe was generated by RT-PCR. mRNA was first isolated from 40 dpf wild-type AB strain ovaries using TRI reagent (Cat. No. T9424; Sigma-Aldrich) and then cDNAs were synthesized with RETROScript Reverse Transcription Kit (Cat. No. AM1710; Thermo Fisher). A *gja11-*specific template was then generated by PCR using Phusion polymerase (Cat. No. M0530L; New England BioLabs) and the following primer pair: fwd, 5’- CCCTGAGCAGTCTTTTCGAGCCT-3’; rev, 5’- taattaatacgactcactataggGTGCTTAAAGCCAGGCGGTCA-3’. Reverse primers contained a T7 RNA polymerase promoter shown in the lower-case letters. An RNA *in situ* probe for *cxcl12a* was generated as previously described (Knaut et al., 2005). For both probes, DNA templates were transcribed with T7 RNA polymerase (Cat. No. 10881775001; Roche) to yield Digoxigenin labeled antisense RNA probes. Probes were G-50 column purified (Cat. No. 45-001-398; GE Healthcare), and then diluted to 0.5-2 mg/ml in the hybridization solution with 5% dextran sulfate (Cat. No. S4030; Thermo Fisher). Fluorescence RNA *in situ* hybridization procedure was performed following Thisse and Thisse (2008) with the following modifications: 40 dpf ovaries were permeabilized with proteinase K at 50 *μ*g/mL for 15 minutes, and the fluorescence *in situs* was developed with FastRed (Cat.No. F4648; Sigma-Aldrich).

### Immunofluorescence staining

Whole ovary immunofluorescence stainings were performed as previously published (Leerberg et al., 2017). Primary chicken anti-Ddx4 antibody (Blokhina et al., 2019) or rabbit anti-Ddx4 antibody (Knaut et al., 2000) were used at 1:1,500. Secondary goat anti-chicken IgG Alexa Fluor 488 (Cat. No. A-11039; Thermo Fisher) or goat anti-rabbit IgG Alexa Fluor 488 (Cat. No. A-11008; Thermo Fisher) were used at 1:300.

### Imaging

Fluorescence RNA *in situ* hybridization, HCR fluorescence RNA *in situ* hybridization, and immunohistochemical staining samples were mounted with ProLong Diamond antifade mountant and imaged with either Olympus FV1000 laser scanning confocal microscope or Zeiss LSM 880 Airyscan microscope.

### Germ cell quantification

Numbers of early GCs expressing *nanos2*, *nanos2* + *foxl2l*, *foxl2l*, or *foxl2l* + *rec8a* in individual clusters were counted in Z-stack images with 1*μ*m steps through the entire cluster. Individual cells must have more than 20 RNA molecules/puncta after HCR fluorescence RNA in situ to be counted as an expressing cell. 70 germ cell clusters were counted from 3 independent experiments with 3 different 40dpf ovaries. The cell count graph was generated with ggplot2 (v3.3.2; Villanueva and Chen, 2019).

### Foxl2l Phylogenetic analysis

Phylogenetic analysis of Foxl proteins was performed using Clustal Omega with the following protein sequences: Carp (Cyprinus carpio): Foxl2l XP_018932368.1; Foxl3 XP_018951720.1; Chicken (*Gallus gallus*): FOXL1 XP_040537252.1, FOXL2 NP_001012630.1; Coelacanth (*Latimeria chalumnae*): Foxl1 XP_005995948.1, Foxl2 XP_006001344.1, Foxl2l XP_005986027.1, Foxl3 XP_005997033.1; Elephant shark (*Callorhinchus milii*): Foxl1 XP_007887429.1; Foxl2l XP_007900370.2; Human (*Homo sapiens*): FOXL1 NP_005241.1, FOXL2 NP_075555.1, FOXL3 NP_001361767.1; Killifish (*Nothobranchius furzeri*): Foxl2l XP_015822944.1; Medaka (*Oryzias latipes*): Foxl1 NP_001116391.1, Foxl2 NP_001098358.1, Foxl2l XP_004070713, Foxl3 XP_011486175.1; Mexican tetra (*Astyanax mexicanus*): Foxl1 XP_015461274.2, Foxl2a XP_007232357.2, Foxl2b XP_007241719.1, Foxl3 XP_007251255.2; Mouse (*Mus musculus*): FOXL1 NP_032050.2;, FOXL2 NP_036150.1, FOXL3 NP_001182057.1; Spotted gar (*Lepisosteus ocula*tus): Foxl1 XP_015223442.1, Foxl2 XP_006637658.1, Foxl2l XP_015192516.1, Foxl3 XP_006637427.1; Sterlet sturgeon (*Acipenser ruthenus*): Foxl2l XP_033909703.2; *Xenopus tropicalis*: Foxl1 XP_012817795.1, Foxl2 XP_004917868.1; Zebrafish (*Danio rerio*): Foxl1 NP_957278.1, Foxl2a, NP_001038717.1, Foxl2b, NP_001304690.1, Foxl2l NP_001122282, Foxl3 NP_001182055.1.

### Mutant production

#### Generation of *foxl2l:egfp* knock-in mutant

We used the Geneweld homology directed repair technique to generate an in frame insertion of a viral 2A peptide-eGFP fusion casset downstream of amino acid 222 (Wierson et al., 2020). A pGTag- *foxl2l-* eGFP-B-actin targeted integration plasmid was designed using the pGTag vector series (Wierson et al., 2020). 48-bp *foxl2l*-specific homology arms were cloned into the pGTag-eGFP-B-actin vector. Upstream homology arm 5’- GCGGgggAAACGCTCTAGTGCCTCTGAGCGGCATGACTCCGCCGGTGAGCCCGGGcGGAT-3’ Downstream homology arm 5’- AAGCGGAAGCTCCATCTCCACCTGCAGTTACGCGCCGCAGAACAGTCACCCcccCCG-3’. Targeted plasmid integration into the *foxl2l* locus was done using CRISPR/Cas9 using a CRISPRscan.org designed sgRNA: 5’-GGGAGATGGAGCTTCCGCCC-3’. Injected F0 fish were outcrossed to wildtype and a sample of F1 embryos were screened for germline transmission of precisely integrated plasmid by PCR, using the following primer sets: 5’-integration primers fwd 5’-- 3’, rev 5’--3’; 3’-integration primers fwd 5’--3’, rev 5’--3’. Precise 5’ integration of the construct was confirmed for the *foxl2l*(*uc91*) allele using Sanger sequencing, and expression of eGFP in germ cells was confirmed at 1.5-2 months post-fertilization. GFP-positive fish were then in-crossed to generate homozygous pGTag-*foxl2l*-eGFP-B-actin transgenic fish. *foxl2l*(uc91) heterozygous fish were in crossed and their offspring genotyped at 50 dpf using standard procedures. The wildtype and knock-in alleles were identified with independent PCR assays using the following primers: Universal_fwd: 5’-GCACATCTCCAGCTACATGC-3’; wild-type_rev: 5’-CACCGAGGTTTGCCATTAGT-3’, ampicon=217 bp; gfp_rev: 5’-CTTCTGCTTGTCGGCCATGATATAG-3’, amplicon=724 bp. Following genotyping, fish were phenotyped for sex using standard criteria (Kossack and Draper, 2019).

#### Generation of *wnt9b* mutant

One-celled zebrafish embryos were injected with Cas9 mRNA (1.4 ng/ embryo) and a gRNA (58 pg/embryo) (5′-<u>CGTGGGAGAGAAGGATGCAG</u>AGG-3′; target underlined) directed against the *wnt9b* locus. The target sequence was identified by using CHOPCHOP (Montague et al., 2014). Target site DNA oligos were annealed and cloned into the BsaI site of pDR274 (Hwang et al., 2013). Vector linearized with DraI was used as template for Maxi Script T7 RNA transcription (Cat. No. AM1312; Thermo Fisher) to generate gRNAs that were purified on a RNeasy Plus column (Cat. No. 74134; Qiagen). Mutagenesis was verified by sequencing individual injected embryo colony PCR products covering the targeted locus. Founder heterozygotes were isolated by outcrossing grown G0 fish to TuAB wild types and genotyping F1 offspring. Primers for genomic PCR: forward, 5′-CCCCTTTAAGAAGTTGCACTGT - 3′; reverse, 5′- CAATGAGGCATTTAGAGGCTTT-3′. An allele consisting of a 57bp deletion was identified which removed the 3’ end of the first exon and 5’ end of the first intron, leading to missplicing and a frameshift leading to a premature stop codon.

### Analysis of *wnt9b* mutants

To determine sex ratios of *wnt9b* mutant fish, 50dpf fish produced from *wnt9b*+/− incross were fixed in 4% paraformaldehyde-1X PBS overnight at 4°C and stored in methanol at −20°C at least overnight. Fish were rehydrated in a methanol:1X PBS gradient and washed three times in 1X PBS prior to dissection. Sex was determined by examining dissected gonads for ovary or testis characteristics. Tissue was collected from each fish during dissection for genotyping post-dissection. Genomic DNA from tissue collected during dissection was extracted and genotyped by PCR. The primers used for genotyping this allele are 5’-CCTCCCCAGGGGCTGAAATA-3’ (forward) and 5’- CAATGAGGCATTTAGAGGCTTT-3’ (reverse). Wildtype allele yields a 238-bp product, *wnt9b(fb209)* allele yields a 181-bp product.

## Supplemental Figures and Legends

**Figure 1-figure supplement 1. A.** Experimental pipeline for the production of the single cell RNA-seq library. Briely, 40 dpf ovaries were isolated from Tg(*piwil1:egfp*) zebrafish and dissociated to single cells using two dissociation methods. The whole ovary dissociation method favored dissociation of somatic cells but led to loss of germ cells. A less stringent method was used to dissociate germ cells from somatic cells, followed by purification of germ cells by fluorescent activated cell sorting (FACS). **B-D.** FACS pseudocolor scatter plots with gating overlays. **B.** The R2 gate selected for GFP+ single cells based on GFP fluorescence intensity (X-axis) and side scatter to measure cell size (Y-axis). **C**. The R3 gate selected for GFP signal (X-axis) relative to cell autofluorescence (Y-axis). The R4 and R5 gates selected for cell size (Y axis) and GFP signal (X-axis). The R5 gate contained smaller cells that were likely premeiotic, meiotic and early-stage oocytes, while R4 gate selected for larger cells that were likely more advanced oocytes. (**E-H**) Representative images of DAPI-stained nuclei from cells obtained from either the R5 (E-G) or R4 gates. E. The prominent single nucleolus (n) contained within this nucleus is indicative of a pre-meiotic germ cell. F-G The diffuse chromatin in these nuclei is characteristic of premeiotic oocyte progenitor cells. **H** The presence of condensed and synapse chromosomes is characteristic of cells that have entered meiosis. Note that the nucleus in H is 30*μ*m in diameter while those in E-F are between 7-10*μ*m in diameter. Scale bars 10*μ*m. (**I-I’**. Fluorescence (I) or bright-field (I’) micrographs of GFP+ cells obtained from gate 5. Scale bars, E-H 10*μ*m; I for I and I’ 250*μ*m.

**Figure 1-figure supplement 2.** Major cell type statistics. **A.** Single-cell UMAP plot of 40-day old zebrafish ovary with combined clusters based on major cell types. **B-D.** Violin plot of percent of mitochondria genes (**B**), number of genes per cell (**C**), and number of transcripts per cell (**D**), in each major cell type in the final dataset.

**Figure 1-figure supplement 3.** Differential expression heatmap and top markers of major cell-types. **A.** Gene expression heatmap of differentially expressed genes between major cell types. Yellow represents highly expressed genes, purple represents lowly expressed genes, and black represents no expression. **B.** Top 5 marker genes computationally identified in each major cell type based on statistical significance.

**Figure 2-figure supplement 1.** Developmental trajectory analysis of the germ cell library using Monocle 3 produced a similar trajectory to that apparent in the SURAT-based cluster analysis. **A.** Germ cell sub-cluster trajectory UMAP plot from Monocle 3. **B.** pseudotime gene expression plots from Monocle 3 for several germ cell relevant genes (see text for details).

**Figure 2-figure supplement 2.** Phylogenetic analysis of Foxl2, Foxl2l, Foxl3 and Foxl1 proteins. *denotes zebrafish Foxl2l (formerly annotated as zgc:194189) and **denotes medaka Foxl2l (formerly called Foxl3). Accession numbers of genes used in this analysis can be found in Methods.

**Figure 2-figure supplement 3.** Expression of *nanos* and *pumilio* orthologs in zebrafish germ cells. Gene expression UMAP plots of select genes. Cells expressing the indicated gene are colored blue, and the relative intensity indicates relative expression levels (intensity scale for each plot is on the right). The UMAP in the upper left corner shows cells color-coded by computationally determined cell subtypes.

**Figure 2-figure supplement 4.** Germ cell library statistics and novel zebrafish GSC markers **A**. Violin plot of the number of genes per cell, number of transcripts per cell, and percent of mitochondria genes in each germ cell sub-clusters. **B**. Gene expression UMAP plots of select GSC-enriched genes. Cells expressing the indicated gene are colored blue, and the relative intensity indicates relative expression levels (intensity scale for each plot is on the right). **C, D.** mFISH on whole-mount 40 dpf ovaries reveals the location confirms expression of *id4* (red in panel **C**; n=18/18 double positive, four replates) and *chga* (red in panel **D**; n=4/22 double positive, four replicates) in *nanos2* expressing germline stem cells (green). In all panels, DNA is gray. 5*μ*m scale bars.

**Figure 2-figure supplement 5.** Gene module analysis and motif enrichment identifies putative GSC-specific transcription factors. **A.** Dot plot of expression level and percent expressed of identified gene modules corresponding to germ cell clusters. **B.** Average expression of genes in selected gene modules plotted in germ cell subcluster. **C.** Gene expression plots and corresponding binding motifs of Figla (module 9). **D.** Gene expression plots and GSC-enriched selected transcription factors that were identified from motifs (module 25). **E.** mFISH on whole-mount 40 dpf ovaries confirms expression of *bhlhe23* (red) in nanos2-expressing GSC (green). DNA is grey. 5*μ*m scale bars.

**Figure 4-figure supplement 1** Gene expression UMAP plots of select follicle cell-enriched genes. Cells expressing the indicated gene in all cells (left column) or in follicle cell subcluster (right column) are colored blue, and the relative intensity indicates relative expression levels (intensity scale for each plot is on the right). Fol, follicle cells; Str, stromal cells; The, theca cells; GC, germ cells; IB, stage IB follicle cells; II, stage II follicle cells, PF, pre-follicle cells.

**Figure 4-figure supplement 2.** Subcluster analysis of pre-follicle cells (*lhx9+).* **A**. Follicle cell sub-cluster UMAP plot, with cells color-coded by computationally determined cell subtypes. The cells outlined, subclusters 1 and 3, were further sub-clustered, generating the UMAP plot shown in **B**. **B**. UMAP plot of the *lhx9* subcluster. For simplicity, the multiple sub-clusters have been combined into only three sub-clusters, which are referred to as SC3.1, SC3.2 and SC1, nomenclature that references their location in the original UMAP sub-cluster plot shown in **B**. **C**. Dot-plot showing the relative expression of select genes in the pre-follicle cell sub-clusters. **D**. Gene expression UMAP plots of genes listed in **B**.

**Figure 4-figure supplement 3.** Gene expression UMAP plots for genes that function in the retinoic acid signaling pathway. *aldh1a2* encodes the last enzyme needed for retinoic acid synthesis. *cyp26a1* encodes a retinoic acid degrading enzyme. *rxrba* and *rxrbb* encode retinoic acid nuclear receptors. Fol, follicle cells; Str, stromal cells; The, theca cells; GC, germ cells.

**Figure 5-figure supplement 1.** Gene expression UMAP plots for Wnt ligand-encoding genes in the 40 dpf ovary. Fol, follicle cells; Str, stromal cells; The, theca cells; Gc, germ cells.

**Figure 6-figure supplement 1.** GO terms associated with the follicle and stromal cell sub-clusters. A. Top 5 GO terms associated with individual follicle sub-clusters. B. Top GO terms associated with the stromal sub-clusters. In both A and B, blue indicates a high and yellow indicates low significance.

**Figure 6-figure supplement 2.** Stromal cell subclusters 1 and 2: pericytes and vascular smooth muscle cells. **A.** Gene expression UMAP plots for select genes whose expression is enriched in stromal cell subcluster 1 and are markers of pericytes. **B.** Gene expression UMAP plots for select genes whose expression is enriched in stromal cell subcluster 2 and are markers of vascular smooth muscle cells.

**Figure 6-figure supplement 3.** Stromal cell subcluster 3: stromal progenitor cells. **A**. Stromal cell sub-cluster UMAP plot, with cells color-coded by computationally determined cell subtypes. **B**. Dot-plot showing the relative expression of select genes in stromal cell subclusters that have enriched expression is sub-cluster 3. **C**. UMAP gene expression plot for genes listed in **B**.

**Figure 6-figure supplement 4.** Max projection view of a confocal stack showing *stm* expression (blue) and DNA (grey) in a 40 dpf ovary. *stm* expression is limited to cells that localize to the lateral and medial edges of the ovary. Scale bar, 200*μ*m.

**Figure 8-figure supplement 1.** Expression of genes involved in theca cell development and steroid synthesis. UMAP gene expression plot for orthologs of genes known to be involved in sex steroid synthesis or theca cell development, mapped onto all cells. Fc, follicle cell; Sc, stromal cell.

**Figure 9-figure supplement 1.** Pathway for sex hormone synthesis in the juvenile zebrafish. Diagram shows a likely pathway for the synthesis of the two main sex steroids in zebrafish, 17ß-estradiol (E2; female) and 11-ketotestosterone (male), starting from cholesterol. Enzymes that catalyze each step are shown, with color indicating sex specific expression. For example, Cyp17a1 is expressed in both males and females, while Cyp19a1a and Hsd17b1 are female-specific while Hsd17b3 and Cyp11c1 are male-specific.

## Notes

https://singlecell.broadinstitute.org/single_cell/study/SCP928

https://github.com/yulongliu68/zeb_ov_ssRNAseq

https://datadryad.org/stash/share/CEd0Zs4oZKdinTWeJPKbWYjBq6hYq4QhVacQcFjf37E

### Competing Interest Statement

The authors have declared no competing interest.

### Summary of Updates

Previous version did not include Figure 1.

## References

Anderson, J.L., Rodriguez Mari, A., Braasch, I., Amores, A., Hohenlohe, P., Batzel, P., and Postlethwait, J.H. (2012). Multiple sex-associated regions and a putative sex chromosome in zebrafish revealed by RAD mapping and population genomics. PLoS One 7, e40701.

Bahrami, N., and Childs, S.J. (2018). Pericyte Biology in Zebrafish. Adv Exp Med Biol 1109, 33–51.

Bailey, T.L., Johnson, J., Grant, C.E., and Noble, W.S. (2015). The MEME Suite. Nucleic Acids Res 43, W39–49.

Bajoghli, B., Ramialison, M., Aghaallaei, N., Czerny, T., and Wittbrodt, J. (2009). Identification of starmaker-like in medaka as a putative target gene of Pax2 in the otic vesicle. Dev Dyn 238, 2860–2866.

Bauer, M.P., Bridgham, J.T., Langenau, D.M., Johnson, A.L., and Goetz, F.W. (2000). Conservation of steroidogenic acute regulatory (StAR) protein structure and expression in vertebrates. Mol Cell Endocrinol 168, 119–125.

Beer, R.L., and Draper, B.W. (2013). nanos3 maintains germline stem cells and expression of the conserved germline stem cell gene nanos2 in the zebrafish ovary. Dev Biol 374, 308–318.

Birk, O.S., Casiano, D.E., Wassif, C.A., Cogliati, T., Zhao, L., Zhao, Y., Grinberg, A., Huang, S., Kreidberg, J.A., Parker, K.L., et al. (2000). The LIM homeobox gene Lhx9 is essential for mouse gonad formation. Nature 403, 909–913.

Blokhina, Y.P., Nguyen, A.D., Draper, B.W., and Burgess, S.M. (2019). The telomere bouquet is a hub where meiotic double-strand breaks, synapsis, and stable homolog juxtaposition are coordinated in the zebrafish, Danio rerio. PLoS Genet 15, e1007730.

Bowles, J., Knight, D., Smith, C., Wilhelm, D., Richman, J., Mamiya, S., Yashiro, K., Chawengsaksophak, K., Wilson, M.J., Rossant, J., et al. (2006). Retinoid signaling determines germ cell fate in mice. Science 312, 596–600.

Brown, L.A., Rodaway, A.R., Schilling, T.F., Jowett, T., Ingham, P.W., Patient, R.K., and Sharrocks, A.D. (2000). Insights into early vasculogenesis revealed by expression of the ETS-domain transcription factor Fli-1 in wild-type and mutant zebrafish embryos. Mech Dev 90, 237–252.

Brunet, J.P., Tamayo, P., Golub, T.R., and Mesirov, J.P. (2004). Metagenes and molecular pattern discovery using matrix factorization. Proc Natl Acad Sci U S A 101, 4164–4169.

Cao, J., Spielmann, M., Qiu, X., Huang, X., Ibrahim, D.M., Hill, A.J., Zhang, F., Mundlos, S., Christiansen, L., Steemers, F.J., et al. (2019a). The single-cell transcriptional landscape of mammalian organogenesis. Nature 566, 496–502.

Cao, Z., Mao, X., and Luo, L. (2019b). Germline Stem Cells Drive Ovary Regeneration in Zebrafish. Cell Rep 26, 1709–1717.e1703.

Carmona, S.J., Teichmann, S.A., Ferreira, L., Macaulay, I.C., Stubbington, M.J., Cvejic, A., and Gfeller, D. (2017). Single-cell transcriptome analysis of fish immune cells provides insight into the evolution of vertebrate immune cell types. Genome Res 27, 451–461.

Carroll, T.J., Park, J.S., Hayashi, S., Majumdar, A., and McMahon, A.P. (2005). Wnt9b plays a central role in the regulation of mesenchymal to epithelial transitions underlying organogenesis of the mammalian urogenital system. Dev Cell 9, 283–292.

Crespo, B., Lan-Chow-Wing, O., Rocha, A., Zanuy, S., and Gómez, A. (2013). foxl2 and foxl3 are two ancient paralogs that remain fully functional in teleosts. Gen Comp Endocrinol 194, 81–93.

Crespo, D., Assis, L.H.C., van de Kant, H.J.G., de Waard, S., Safian, D., Lemos, M.S., Bogerd, J., and Schulz, R.W. (2019). Endocrine and local signaling interact to regulate spermatogenesis in zebrafish: follicle-stimulating hormone, retinoic acid and androgens. Development 146.

Cui, S., Ross, A., Stallings, N., Parker, K.L., Capel, B., and Quaggin, S.E. (2004). Disrupted gonadogenesis and male-to-female sex reversal in Pod1 knockout mice. Development 131, 4095–4105.

Cunningham, F., Achuthan, P., Akanni, W., Allen, J., Amode, M.R., Armean, I.M., Bennett, R., Bhai, J., Billis, K., Boddu, S., et al. (2019). Ensembl 2019. Nucleic Acids Res 47, D745–d751.

De Keuckelaere, E., Hulpiau, P., Saeys, Y., Berx, G., and van Roy, F. (2018). Nanos genes and their role in development and beyond. Cell Mol Life Sci 75, 1929–1946.

Devlin, R.H., and Nagahama, Y. (2002). Sex determination and sex differentiation in fish: an overview of genetic, physiological, and environmental influences. Aquaculture 208, 191–364.

Doitsidou, M., Reichman-Fried, M., Stebler, J., Koprunner, M., Dorries, J., Meyer, D., Esguerra, C.V., Leung, T., and Raz, E. (2002). Guidance of primordial germ cell migration by the chemokine SDF-1. Cell 111, 647–659.

Dranow, D.B., Hu, K., Bird, A.M., Lawry, S.T., Adams, M.T., Sanchez, A., Amatruda, J.F., and Draper, B.W. (2016). Bmp15 Is an Oocyte-Produced Signal Required for Maintenance of the Adult Female Sexual Phenotype in Zebrafish. PLoS Genet 12, e1006323.

Draper, B.W. (2012). Identification of oocyte progenitor cells in the zebrafish ovary. Methods Mol Biol 916, 157–165.

Draper, B.W., McCallum, C.M., and Moens, C.B. (2007). nanos1 is required to maintain oocyte production in adult zebrafish. Dev Biol 305, 589–598.

Elkouby, Y.M., and Mullins, M.C. (2017). Methods for the analysis of early oogenesis in Zebrafish. Dev Biol 430, 310–324.

Farrell, J.A., Wang, Y., Riesenfeld, S.J., Shekhar, K., Regev, A., and Schier, A.F. (2018). Single-cell reconstruction of developmental trajectories during zebrafish embryogenesis. Science 360.

Forbes, A., and Lehmann, R. (1998). Nanos and Pumilio have critical roles in the development and function of Drosophila germline stem cells. Development 125, 679–690.

Fu, A., Oberholtzer, S.M., Bagheri-Fam, S., Rastetter, R.H., Holdreith, C., Caceres, V.L., John, S.V., Shaw, S.A., Krentz, K.J., Zhang, X., et al. (2018). Dynamic expression patterns of Irx3 and Irx5 during germline nest breakdown and primordial follicle formation promote follicle survival in mouse ovaries. PLoS Genet 14, e1007488.

Fürthauer, M., Reifers, F., Brand, M., Thisse, B., and Thisse, C. (2001). sprouty4 acts in vivo as a feedback-induced antagonist of FGF signaling in zebrafish. Development 128, 2175–2186.

Gaengel, K., Genové, G., Armulik, A., and Betsholtz, C. (2009). Endothelial-mural cell signaling in vascular development and angiogenesis. Arterioscler Thromb Vasc Biol 29, 630–638.

Gautier, A., Sohm, F., Joly, J.S., Le Gac, F., and Lareyre, J.J. (2011). The proximal promoter region of the zebrafish gsdf gene is sufficient to mimic the spatio-temporal expression pattern of the endogenous gene in Sertoli and granulosa cells. Biol Reprod 85, 1240–1251.

Gilchrist, R.B., Lane, M., and Thompson, J.G. (2008). Oocyte-secreted factors: regulators of cumulus cell function and oocyte quality. Hum Reprod Update 14, 159–177.

Goldenberg, R.L., Vaitukaitis, J.L., and Ross, G.T. (1972). Estrogen and follicle stimulation hormone interactions on follicle growth in rats. Endocrinology 90, 1492–1498.

Gore, A.V., Athans, B., Iben, J.R., Johnson, K., Russanova, V., Castranova, D., Pham, V.N., Butler, M.G., Williams-Simons, L., Nichols, J.T., et al. (2016). Epigenetic regulation of hematopoiesis by DNA methylation. Elife 5, e11813.

Guiguen, Y., Fostier, A., Piferrer, F., and Chang, C.F. (2010). Ovarian aromatase and estrogens: a pivotal role for gonadal sex differentiation and sex change in fish. Gen Comp Endocrinol 165, 352–366.

Haddon, C., Jiang, Y.J., Smithers, L., and Lewis, J. (1998). Delta-Notch signalling and the patterning of sensory cell differentiation in the zebrafish ear: evidence from the mind bomb mutant. Development 125, 4637–4644.

Hsu, R.J., Lin, C.Y., Hoi, H.S., Zheng, S.K., Lin, C.C., and Tsai, H.J. (2010). Novel intronic microRNA represses zebrafish myf5 promoter activity through silencing dickkopf-3 gene. Nucleic Acids Res 38, 4384–4393.

Hu, Y.C., Okumura, L.M., and Page, D.C. (2013). Gata4 is required for formation of the genital ridge in mice. PLoS Genet 9, e1003629.

Hwang, W.Y., Fu, Y., Reyon, D., Maeder, M.L., Tsai, S.Q., Sander, J.D., Peterson, R.T., Yeh, J.R., and Joung, J.K. (2013). Efficient genome editing in zebrafish using a CRISPR-Cas system. Nat Biotechnol 31, 227–229.

Ings, J.S., and Van Der Kraak, G.J. (2006). Characterization of the mRNA expression of StAR and steroidogenic enzymes in zebrafish ovarian follicles. Mol Reprod Dev 73, 943–954.

Jovelin, R., He, X., Amores, A., Yan, Y.L., Shi, R., Qin, B., Roe, B., Cresko, W.A., and Postlethwait, J.H. (2007). Duplication and divergence of fgf8 functions in teleost development and evolution. J Exp Zool B Mol Dev Evol 308, 730–743.

Kidder, G.M., and Vanderhyden, B.C. (2010). Bidirectional communication between oocytes and follicle cells: ensuring oocyte developmental competence. Can J Physiol Pharmacol 88, 399–413.

Kikuchi, K., Holdway, J.E., Major, R.J., Blum, N., Dahn, R.D., Begemann, G., and Poss, K.D. (2011). Retinoic acid production by endocardium and epicardium is an injury response essential for zebrafish heart regeneration. Dev Cell 20, 397–404.

Kim, B., Kim, Y., Cooke, P.S., Rüther, U., and Jorgensen, J.S. (2011). The fused toes locus is essential for somatic-germ cell interactions that foster germ cell maturation in developing gonads in mice. Biol Reprod 84, 1024–1032.

Kinnear, H.M., Tomaszewski, C.E., Chang, F.L., Moravek, M.B., Xu, M., Padmanabhan, V., and Shikanov, A. (2020). The ovarian stroma as a new frontier. Reproduction 160, R25–r39.

Knaut, H., Blader, P., Strähle, U., and Schier, A.F. (2005). Assembly of trigeminal sensory ganglia by chemokine signaling. Neuron 47, 653–666.

Knaut, H., Werz, C., Geisler, R., and Nusslein-Volhard, C. (2003). A zebrafish homologue of the chemokine receptor Cxcr4 is a germ-cell guidance receptor. Nature 421, 279–282.

Koprunner, M., Thisse, C., Thisse, B., and Raz, E. (2001). A zebrafish nanos-related gene is essential for the development of primordial germ cells. Genes Dev 15, 2877–2885.

Kossack, M.E., and Draper, B.W. (2019). Genetic regulation of sex determination and maintenance in zebrafish (Danio rerio). Curr Top Dev Biol 134, 119–149.

Kossack, M.E., High, S.K., Hopton, R.E., Yan, Y.L., Postlethwait, J.H., and Draper, B.W. (2019). Female Sex Development and Reproductive Duct Formation Depend on Wnt4a in Zebrafish. Genetics 211, 219–233.

Koubova, J., Menke, D.B., Zhou, Q., Capel, B., Griswold, M.D., and Page, D.C. (2006). Retinoic acid regulates sex-specific timing of meiotic initiation in mice. Proc Natl Acad Sci U S A 103, 2474–2479.

Kreidberg, J.A., Sariola, H., Loring, J.M., Maeda, M., Pelletier, J., Housman, D., and Jaenisch, R. (1993). WT-1 is required for early kidney development. Cell 74, 679–691.

Kwok, H.F., So, W.K., Wang, Y., and Ge, W. (2005). Zebrafish gonadotropins and their receptors: I. Cloning and characterization of zebrafish follicle-stimulating hormone and luteinizing hormone receptors--evidence for their distinct functions in follicle development. Biol Reprod 72, 1370–1381.

Lawson, N.D., and Weinstein, B.M. (2002). In vivo imaging of embryonic vascular development using transgenic zebrafish. Dev Biol 248, 307–318.

Leerberg, D.M., Sano, K., and Draper, B.W. (2017). Fibroblast growth factor signaling is required for early somatic gonad development in zebrafish. PLoS Genet 13, e1006993.

Leu, D.H., and Draper, B.W. (2010). The ziwi promoter drives germline-specific gene expression in zebrafish. Dev Dyn 239, 2714–2721.

Liang, L., Soyal, S.M., and Dean, J. (1997). FIGalpha, a germ cell specific transcription factor involved in the coordinate expression of the zona pellucida genes. Development 124, 4939–4947.

Lieschke, G.J., Oates, A.C., Crowhurst, M.O., Ward, A.C., and Layton, J.E. (2001). Morphologic and functional characterization of granulocytes and macrophages in embryonic and adult zebrafish. Blood 98, 3087–3096.

Luo, X., Ikeda, Y., and Parker, K.L. (1994). A cell-specific nuclear receptor is essential for adrenal and gonadal development and sexual differentiation. Cell 77, 481–490.

Marlow, F.L., and Mullins, M.C. (2008). Bucky ball functions in Balbiani body assembly and animal-vegetal polarity in the oocyte and follicle cell layer in zebrafish. Dev Biol 321, 40–50.

McGinnis, C.S., Murrow, L.M., and Gartner, Z.J. (2019). DoubletFinder: Doublet Detection in Single-Cell RNA Sequencing Data Using Artificial Nearest Neighbors. Cell Syst 8, 329–337.e324.

McLaren, A. (1991). Development of the mammalian gonad: the fate of the supporting cell lineage. Bioessays 13, 151–156.

McLeay, R.C., and Bailey, T.L. (2010). Motif Enrichment Analysis: a unified framework and an evaluation on ChIP data. BMC Bioinformatics 11, 165.

Miller, W.L., and Auchus, R.J. (2011). The molecular biology, biochemistry, and physiology of human steroidogenesis and its disorders. Endocr Rev 32, 81–151.

Mindnich, R., Deluca, D., and Adamski, J. (2004). Identification and characterization of 17 beta-hydroxysteroid dehydrogenases in the zebrafish, Danio rerio. Mol Cell Endocrinol 215, 19–30.

Mindnich, R., Haller, F., Halbach, F., Moeller, G., Hrabé de Angelis, M., and Adamski, J. (2005). Androgen metabolism via 17beta-hydroxysteroid dehydrogenase type 3 in mammalian and non-mammalian vertebrates: comparison of the human and the zebrafish enzyme. J Mol Endocrinol 35, 305–316.

Miyamoto, N., Yoshida, M., Kuratani, S., Matsuo, I., and Aizawa, S. (1997). Defects of urogenital development in mice lacking Emx2. Development 124, 1653–1664.

Montague, T.G., Cruz, J.M., Gagnon, J.A., Church, G.M., and Valen, E. (2014). CHOPCHOP: a CRISPR/Cas9 and TALEN web tool for genome editing. Nucleic Acids Res 42, W401–407.

Morvan-Dubois, G., Le Guellec, D., Garrone, R., Zylberberg, L., and Bonnaud, L. (2003). Phylogenetic analysis of vertebrate fibrillar collagen locates the position of zebrafish alpha3(I) and suggests an evolutionary link between collagen alpha chains and hox clusters. J Mol Evol 57, 501–514.

Mullen, R.D., and Behringer, R.R. (2014). Molecular genetics of Müllerian duct formation, regression and differentiation. Sex Dev 8, 281–296.

Nakamura, S., Aoki, Y., Saito, D., Kuroki, Y., Fujiyama, A., Naruse, K., and Tanaka, M. (2008). Sox9b/sox9a2-EGFP transgenic medaka reveals the morphological reorganization of the gonads and a common precursor of both the female and male supporting cells. Mol Reprod Dev 75, 472–476.

Nakamura, S., Kobayashi, K., Nishimura, T., Higashijima, S., and Tanaka, M. (2010). Identification of germline stem cells in the ovary of the teleost medaka. Science 328, 1561–1563.

Nicol, B., and Guiguen, Y. (2011). Expression profiling of Wnt signaling genes during gonadal differentiation and gametogenesis in rainbow trout. Sex Dev 5, 318–329.

Nishimura, T., Sato, T., Yamamoto, Y., Watakabe, I., Ohkawa, Y., Suyama, M., Kobayashi, S., and Tanaka, M. (2015). Sex determination. foxl3 is a germ cell-intrinsic factor involved in sperm-egg fate decision in medaka. Science 349, 328–331.

Oatley, M.J., Kaucher, A.V., Racicot, K.E., and Oatley, J.M. (2011). Inhibitor of DNA binding 4 is expressed selectively by single spermatogonia in the male germline and regulates the self-renewal of spermatogonial stem cells in mice. Biol Reprod 85, 347–356.

Onichtchouk, D., Aduroja, K., Belting, H.G., Gnügge, L., and Driever, W. (2003). Transgene driving GFP expression from the promoter of the zona pellucida gene zpc is expressed in oocytes and provides an early marker for gonad differentiation in zebrafish. Dev Dyn 228, 393–404.

Parajes, S., Griffin, A., Taylor, A.E., Rose, I.T., Miguel-Escalada, I., Hadzhiev, Y., Arlt, W., Shackleton, C., Müller, F., and Krone, N. (2013). Redefining the initiation and maintenance of zebrafish interrenal steroidogenesis by characterizing the key enzyme cyp11a2. Endocrinology 154, 2702–2711.

Pepling, M.E., de Cuevas, M., and Spradling, A.C. (1999). Germline cysts: a conserved phase of germ cell development? Trends Cell Biol 9, 257–262.

Pereira, F.A., Qiu, Y., Tsai, M.J., and Tsai, S.Y. (1995). Chicken ovalbumin upstream promoter transcription factor (COUP-TF): expression during mouse embryogenesis. J Steroid Biochem Mol Biol 53, 503–508.

Petit, I., Jin, D., and Rafii, S. (2007). The SDF-1-CXCR4 signaling pathway: a molecular hub modulating neo-angiogenesis. Trends Immunol 28, 299–307.

Prasasya, R.D., and Mayo, K.E. (2018). Notch Signaling Regulates Differentiation and Steroidogenesis in Female Mouse Ovarian Granulosa Cells. Endocrinology 159, 184–198.

Qin, M., Zhang, Z., Song, W., Wong, Q.W., Chen, W., Shirgaonkar, N., and Ge, W. (2018). Roles of Figla/figla in Juvenile Ovary Development and Follicle Formation During Zebrafish Gonadogenesis. Endocrinology 159, 3699–3722.

Raible, F., and Brand, M. (2001). Tight transcriptional control of the ETS domain factors Erm and Pea3 by Fgf signaling during early zebrafish development. Mech Dev 107, 105–117.

Raudvere, U., Kolberg, L., Kuzmin, I., Arak, T., Adler, P., Peterson, H., and Vilo, J. (2019). g:Profiler: a web server for functional enrichment analysis and conversions of gene lists (2019 update). Nucleic Acids Res 47, W191–w198.

Rodriguez-Mari, A., Canestro, C., BreMiller, R.A., Catchen, J.M., Yan, Y.L., and Postlethwait, J.H. (2013). Retinoic acid metabolic genes, meiosis, and gonadal sex differentiation in zebrafish. PLoS One 8, e73951.

Rodriguez-Mari, A., Canestro, C., Bremiller, R.A., Nguyen-Johnson, A., Asakawa, K., Kawakami, K., and Postlethwait, J.H. (2010). Sex reversal in zebrafish fancl mutants is caused by Tp53-mediated germ cell apoptosis. PLoS Genet 6, e1001034.

Rodriguez-Mari, A., Yan, Y.L., Bremiller, R.A., Wilson, C., Canestro, C., and Postlethwait, J.H. (2005). Characterization and expression pattern of zebrafish Anti-Mullerian hormone (Amh) relative to sox9a, sox9b, and cyp19a1a, during gonad development. Gene Expr Patterns 5, 655–667.

Roehl, H., and Nüsslein-Volhard, C. (2001). Zebrafish pea3 and erm are general targets of FGF8 signaling. Curr Biol 11, 503–507.

Roly, Z.Y., Godini, R., Estermann, M.A., Major, A.T., Pocock, R., and Smith, C.A. (2020). Transcriptional landscape of the embryonic chicken Müllerian duct. BMC Genomics 21, 688.

Ruzicka, L., Howe, D.G., Ramachandran, S., Toro, S., Van Slyke, C.E., Bradford, Y.M., Eagle, A., Fashena, D., Frazer, K., Kalita, P., et al. (2019). The Zebrafish Information Network: new support for non-coding genes, richer Gene Ontology annotations and the Alliance of Genome Resources. Nucleic Acids Res 47, D867–d873.

Ryan, K.J. (1979). Granulosa-thecal cell interaction in ovarian steroidogenesis. J Steroid Biochem 11, 799–800.

Selman, K., Wallace, R.A., Sarka, A., and Qi, X. (1993). Stages of oocyte development in the zebrafish, Brachydanio rerio. J Morphol 218, 203–224.

Shen, Y.C., Shami, A.N., Moritz, L., Larose, H., Manske, G.L., Ma, Q., Zheng, X., Sukhwani, M., Czerwinski, M., Sultan, C., et al. (2021). TCF21(+) mesenchymal cells contribute to testis somatic cell development, homeostasis, and regeneration in mice. Nat Commun 12, 3876.

Shive, H.R., West, R.R., Embree, L.J., Azuma, M., Sood, R., Liu, P., and Hickstein, D.D. (2010). brca2 in zebrafish ovarian development, spermatogenesis, and tumorigenesis. Proc Natl Acad Sci U S A 107, 19350–19355.

Siebert, S., Farrell, J.A., Cazet, J.F., Abeykoon, Y., Primack, A.S., Schnitzler, C.E., and Juliano, C.E. (2019). Stem cell differentiation trajectories in Hydra resolved at single-cell resolution. Science 365.

So, P.L., and Danielian, P.S. (1999). Cloning and expression analysis of a mouse gene related to Drosophila odd-skipped. Mech Dev 84, 157–160.

Söllner, C., Burghammer, M., Busch-Nentwich, E., Berger, J., Schwarz, H., Riekel, C., and Nicolson, T. (2003). Control of crystal size and lattice formation by starmaker in otolith biomineralization. Science 302, 282–286.

Soyal, S.M., Amleh, A., and Dean, J. (2000). FIGalpha, a germ cell-specific transcription factor required for ovarian follicle formation. Development 127, 4645–4654.

Stuart, T., Butler, A., Hoffman, P., Hafemeister, C., Papalexi, E., Mauck, W.M., 3rd, Hao, Y., Stoeckius, M., Smibert, P., and Satija, R. (2019). Comprehensive Integration of Single-Cell Data. Cell 177, 1888–1902.e1821.

Su, Y.Q., Sugiura, K., Wigglesworth, K., O’Brien, M.J., Affourtit, J.P., Pangas, S.A., Matzuk, M.M., and Eppig, J.J. (2008). Oocyte regulation of metabolic cooperativity between mouse cumulus cells and oocytes: BMP15 and GDF9 control cholesterol biosynthesis in cumulus cells. Development 135, 111–121.

Suzuki, A., Saba, R., Miyoshi, K., Morita, Y., and Saga, Y. (2012). Interaction between NANOS2 and the CCR4-NOT deadenylation complex is essential for male germ cell development in mouse. PLoS One 7, e33558.

Takahashi, H. (1977). Juvenile Hermaphroditism in the zebrafish, Brachydanio rerio. Bull Fac Fish Hokkaido Univ 28, 57–65.

Tang, Q., Iyer, S., Lobbardi, R., Moore, J.C., Chen, H., Lareau, C., Hebert, C., Shaw, M.L., Neftel, C., Suva, M.L., et al. (2017). Dissecting hematopoietic and renal cell heterogeneity in adult zebrafish at single-cell resolution using RNA sequencing. J Exp Med 214, 2875–2887.

Thisse, C., and Thisse, B. (2008). High-resolution in situ hybridization to whole-mount zebrafish embryos. Nat Protoc 3, 59–69.

Tokarz, J., Moller, G., de Angelis, M.H., and Adamski, J. (2013). Zebrafish and steroids: what do we know and what do we need to know? J Steroid Biochem Mol Biol 137, 165–173.

Tourtellotte, W.G., Nagarajan, R., Auyeung, A., Mueller, C., and Milbrandt, J. (1999). Infertility associated with incomplete spermatogenic arrest and oligozoospermia in Egr4-deficient mice. Development 126, 5061–5071.

Tsukita, S., Tanaka, H., and Tamura, A. (2019). The Claudins: From Tight Junctions to Biological Systems. Trends Biochem Sci 44, 141–152.

Vainio, S., Heikkilä, M., Kispert, A., Chin, N., and McMahon, A.P. (1999). Female development in mammals is regulated by Wnt-4 signalling. Nature 397, 405–409.

Wagner, M., Yoshihara, M., Douagi, I., Damdimopoulos, A., Panula, S., Petropoulos, S., Lu, H., Pettersson, K., Palm, K., Katayama, S., et al. (2020). Single-cell analysis of human ovarian cortex identifies distinct cell populations but no oogonial stem cells. Nat Commun 11, 1147.

Wang, Q., Lan, Y., Cho, E.S., Maltby, K.M., and Jiang, R. (2005). Odd-skipped related 1 (Odd 1) is an essential regulator of heart and urogenital development. Dev Biol 288, 582–594.

Wang, Y., Pan, L., Moens, C.B., and Appel, B. (2014). Notch3 establishes brain vascular integrity by regulating pericyte number. Development 141, 307–317.

Webster, K.A., Schach, U., Ordaz, A., Steinfeld, J.S., Draper, B.W., and Siegfried, K.R. (2017). Dmrt1 is necessary for male sexual development in zebrafish. Dev Biol 422, 33–46.

Westerfield, M. (2000). The zebrafish book. A guid to the laboratory use of zebrafish (Danio rerio). 4th edn (Univ. of Oregon Press, Eugene OR).

Whitesell, T.R., Kennedy, R.M., Carter, A.D., Rollins, E.L., Georgijevic, S., Santoro, M.M., and Childs, S.J. (2014). An α-smooth muscle actin (acta2/αsma) zebrafish transgenic line marking vascular mural cells and visceral smooth muscle cells. PLoS One 9, e90590.

Wierson, W.A., Welker, J.M., Almeida, M.P., Mann, C.M., Webster, D.A., Torrie, M.E., Weiss, T.J., Kambakam, S., Vollbrecht, M.K., Lan, M., et al. (2020). Efficient targeted integration directed by short homology in zebrafish and mammalian cells. Elife 9.

Wilson, C.A., High, S.K., McCluskey, B.M., Amores, A., Yan, Y.L., Titus, T.A., Anderson, J.L., Batzel, P., Carvan, M.J., 3rd, Schartl, M., et al. (2014). Wild sex in zebrafish: loss of the natural sex determinant in domesticated strains. Genetics 198, 1291–1308.

Xie, J., Wang, W.Q., Liu, T.X., Deng, M., and Ning, G. (2008). Spatio-temporal expression of chromogranin A during zebrafish embryogenesis. J Endocrinol 198, 451–458.

Yan, Y.L., Batzel, P., Titus, T., Sydes, J., Desvignes, T., Bremiller, R., Draper, B., and Postlethwait, J.H. (2019). A Hormone That Lost Its Receptor: Anti-Mullerian Hormone (AMH) in Zebrafish Gonad Development and Sex Determination. Genetics.

Yang, Y.J., Wang, Y., Li, Z., Zhou, L., and Gui, J.F. (2017). Sequential, Divergent, and Cooperative Requirements of Foxl2a and Foxl2b in Ovary Development and Maintenance of Zebrafish. Genetics 205, 1551–1572.

Yoon, C., Kawakami, K., and Hopkins, N. (1997). Zebrafish vasa homologue RNA is localized to the cleavage planes of 2- and 4-cell-stage embryos and is expressed in the primordial germ cells. Development 124, 3157–3165.

Yoshida, K., Kondoh, G., Matsuda, Y., Habu, T., Nishimune, Y., and Morita, T. (1998). The mouse RecA-like gene Dmc1 is required for homologous chromosome synapsis during meiosis. Mol Cell 1, 707–718.

Young, M.D., and Behjati, S. (2020). SoupX removes ambient RNA contamination from droplet-based single-cell RNA sequencing data. Gigascience 9.

Zakrzewska, A., Cui, C., Stockhammer, O.W., Benard, E.L., Spaink, H.P., and Meijer, A.H. (2010). Macrophage-specific gene functions in Spi1-directed innate immunity. Blood 116, e1–11.

