## Supplemental Figures and Legends for "Single-cell transcriptome reveals insights into the development and function of the zebrafish ovary"

**Figure 9-figure supplement 1.** Pathway for sex hormone synthesis in the juvenile zebrafish. Diagram shows a likely pathway for the synthesis of the two main sex steroids in zebrafish, 17ß-estradiol (E2; female) and 11-ketotestosterone (male), starting from cholesterol. Enzymes that catalyze each step are shown, with color indicating sex specific expression. For example, Cyp17a1 is expressed in both males and females, while Cyp19a1a and Hsd17b1 are female-specific while Hsd17b3 and Cyp11c1 are male-specific.

#### Figure 1- figure supplement 1

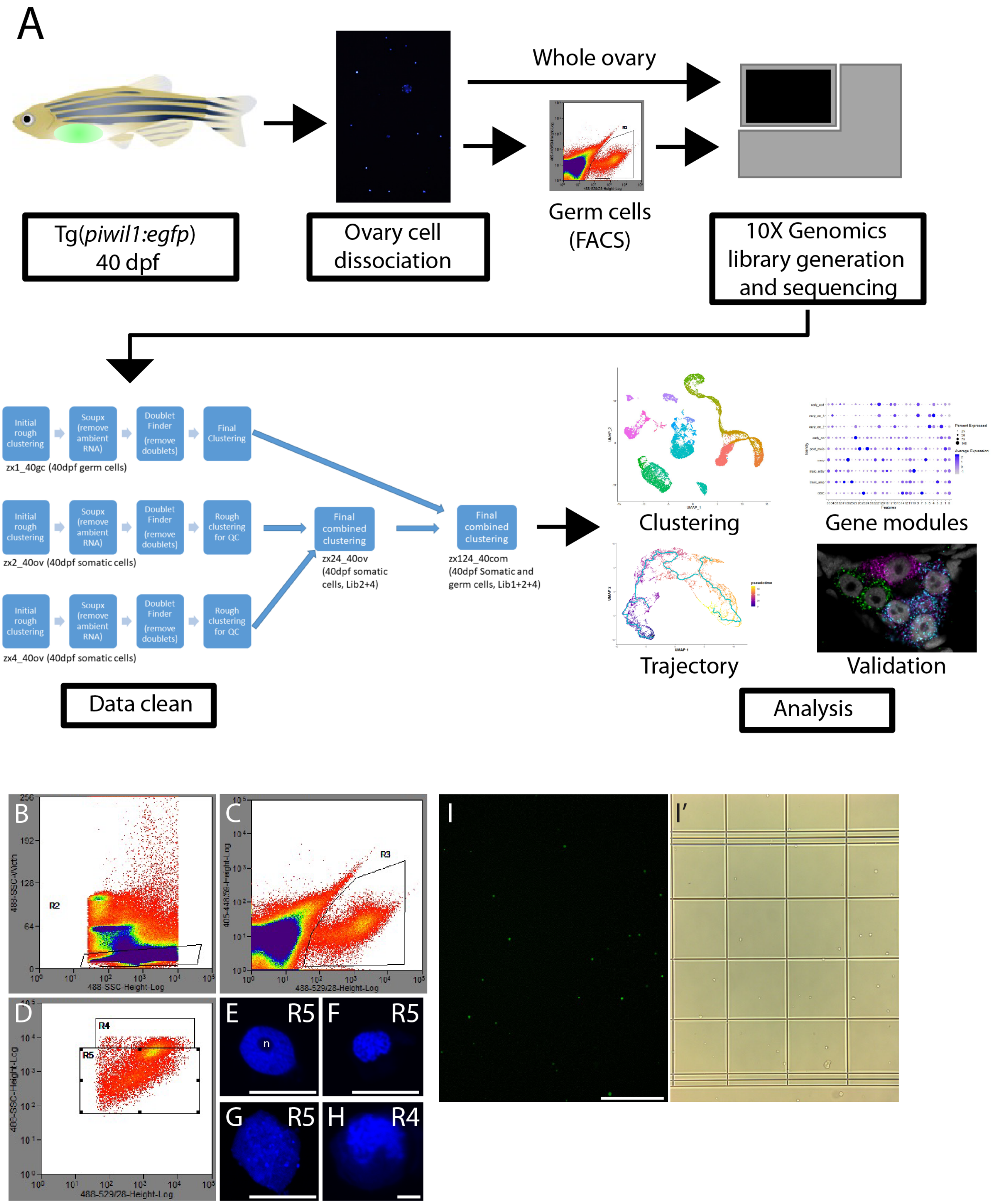

#### Figure 1- figure supplement 2

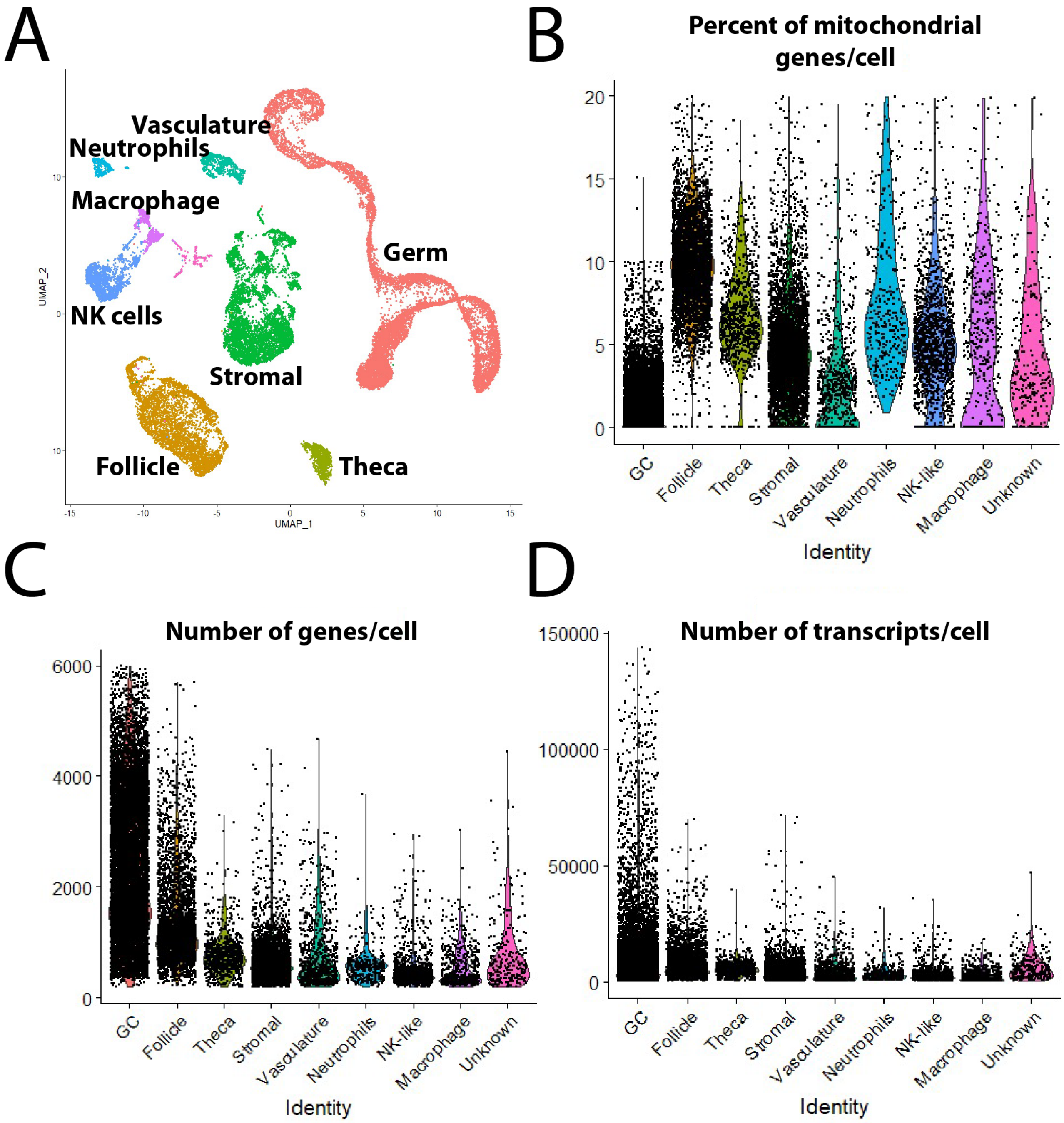

Figure 1- figure supplement 3

A.

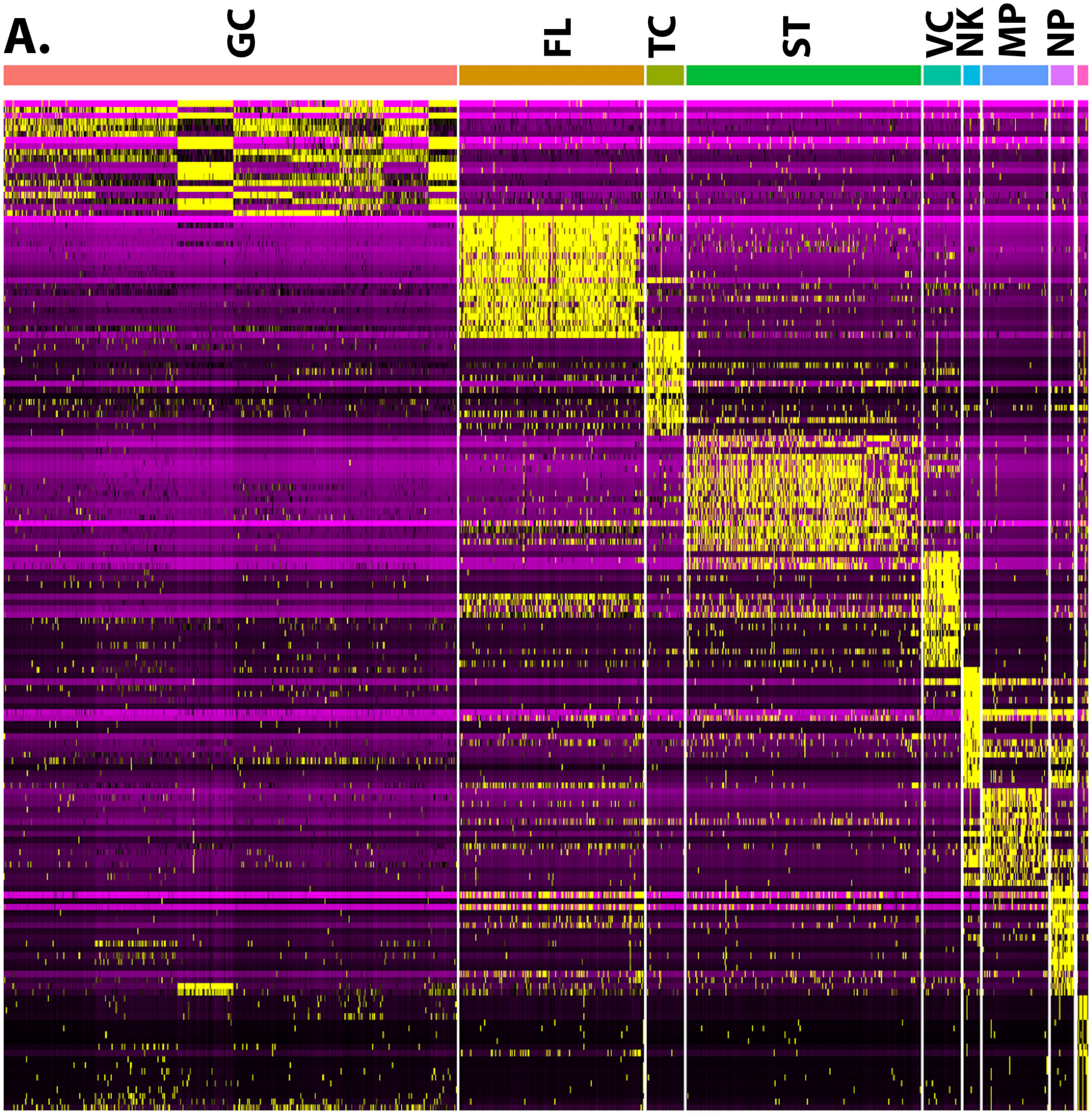


B.

| GC | FL | TC | ST | VC | NP | NK | MP |
|---|---|---|---|---|---|---|---|
| *ddx4* | *gsdf* | *cy917a1* | *col1a1a* | *cdh5* | *lyz* | *ccl36.1* | *grn1* |
| *sycp1* | *hsd19b1* | *star* | *nid1b* | *etv2* | *lect21* | *nkl.2* | *ccl35.1* |
| *zp3.2* | *her2* | *cyp11a2* | *krt15* | *sox7* | *mpx* | *wasb* | *ccr9a* |
| *piwil1* | *cldn11a* | *ddx1b* | *tagln* | *aoc2* | *cpa5* | *laptm5* | *c1qb* |
| *nasp* | *notch3* | *fgg* | *anxa1a* | *krt8* | *cxcr4b* | *arpc1b* | *lygl1* |

#### Figure 2- figure supplement 1

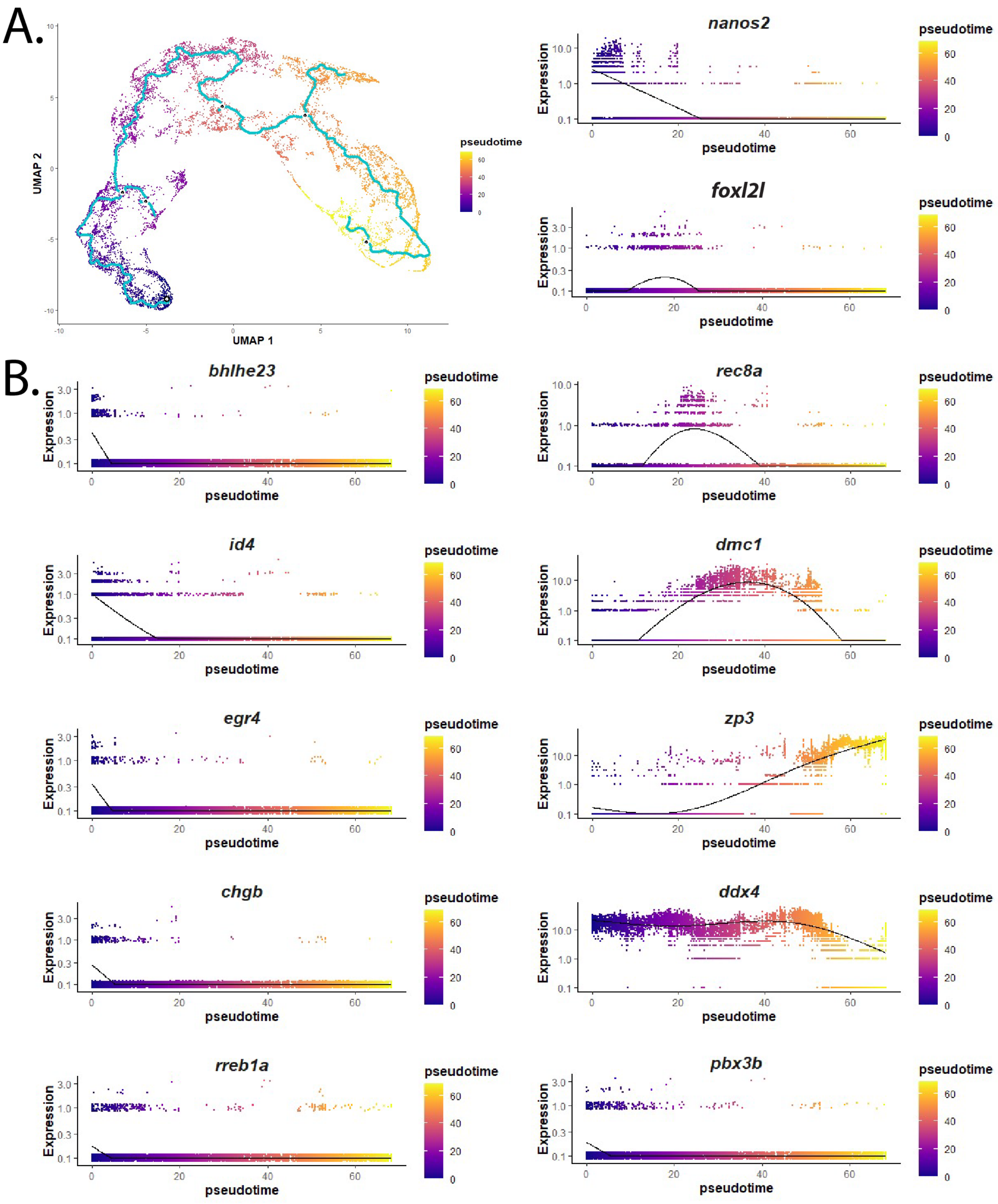

**Figure 2- figure supplement 2**

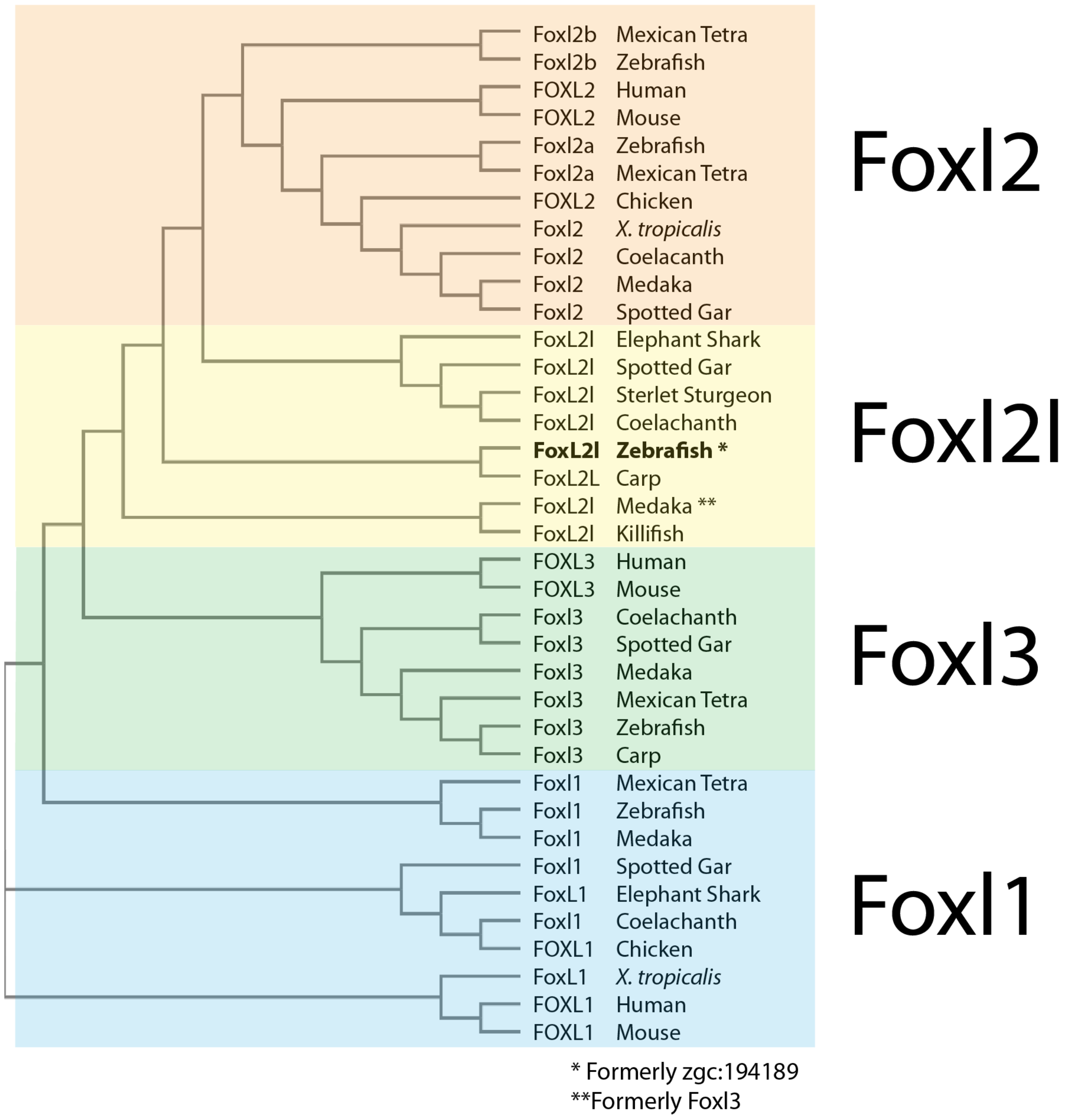

**Figure 2- figure supplement 3**

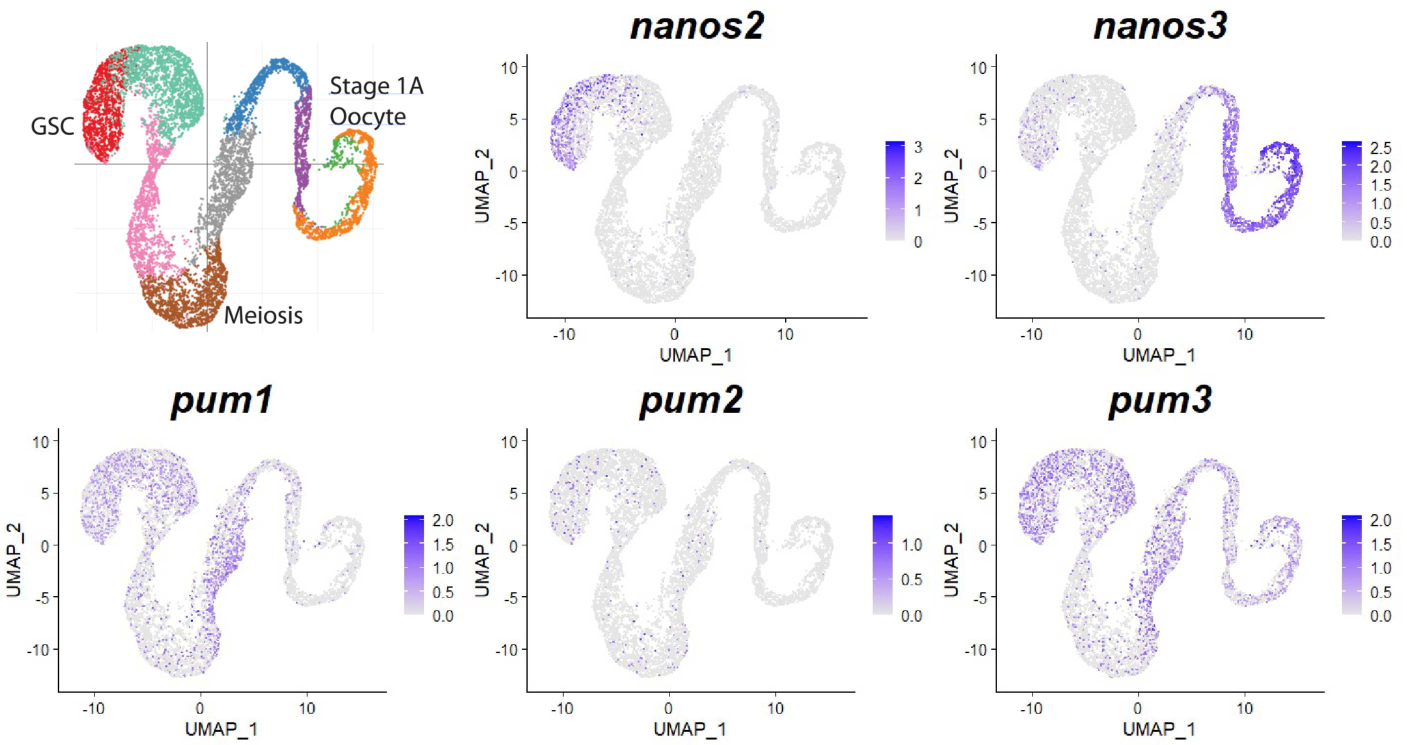

#### Figure 2- figure supplement 4

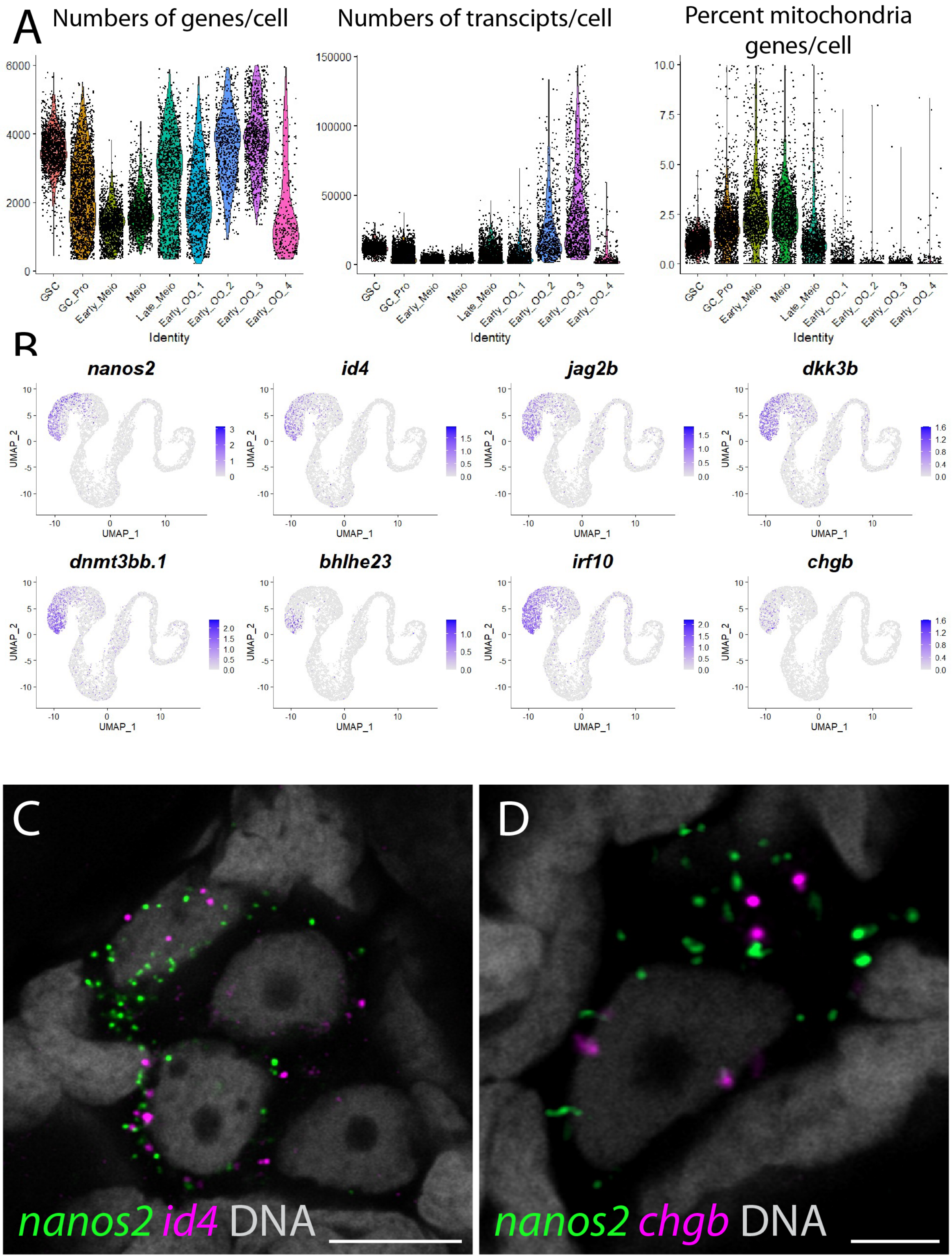

**Figure 2- figure supplement 5**

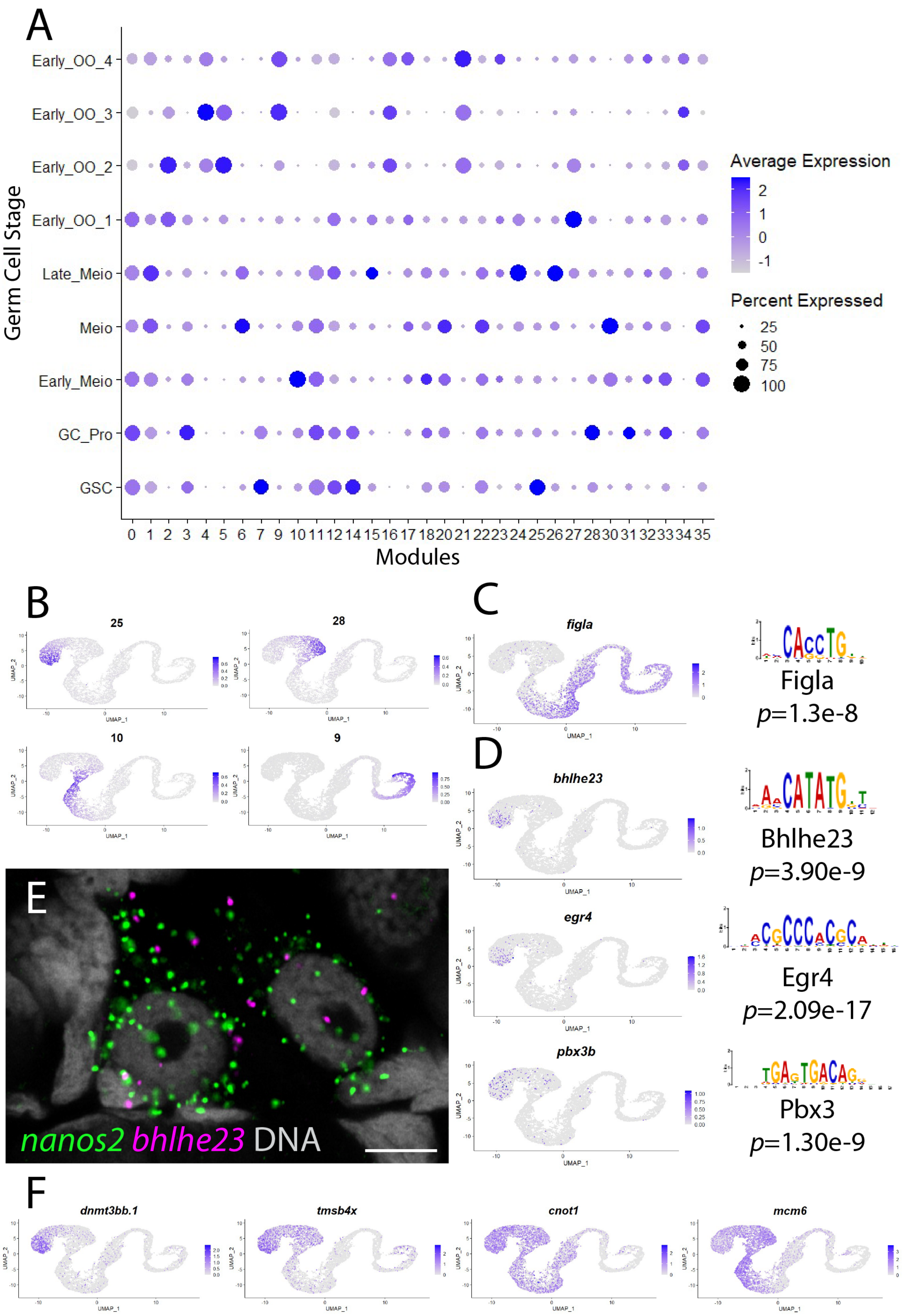

### Figure 4- figure supplement 1

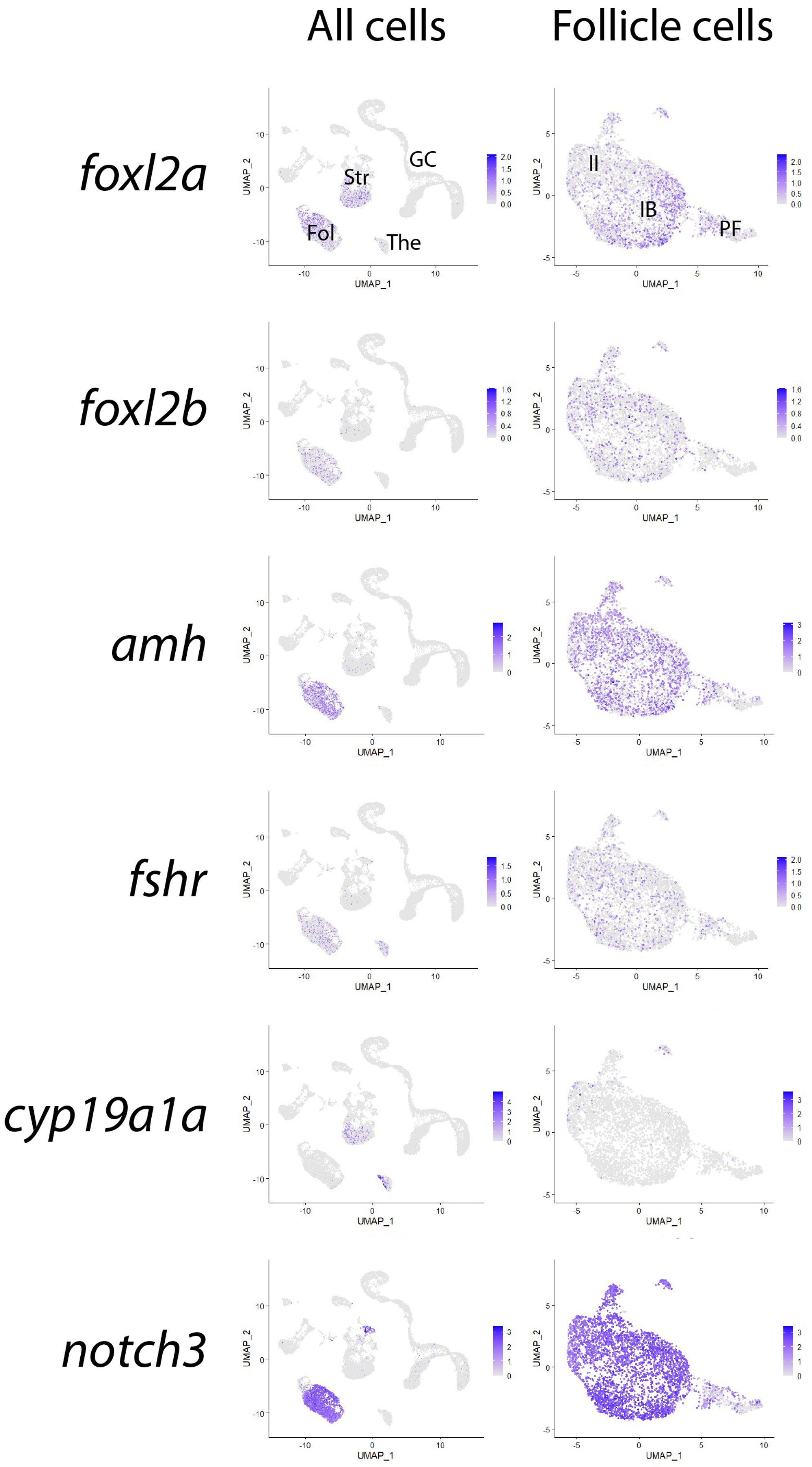

**Figure 4- figure supplement 2**

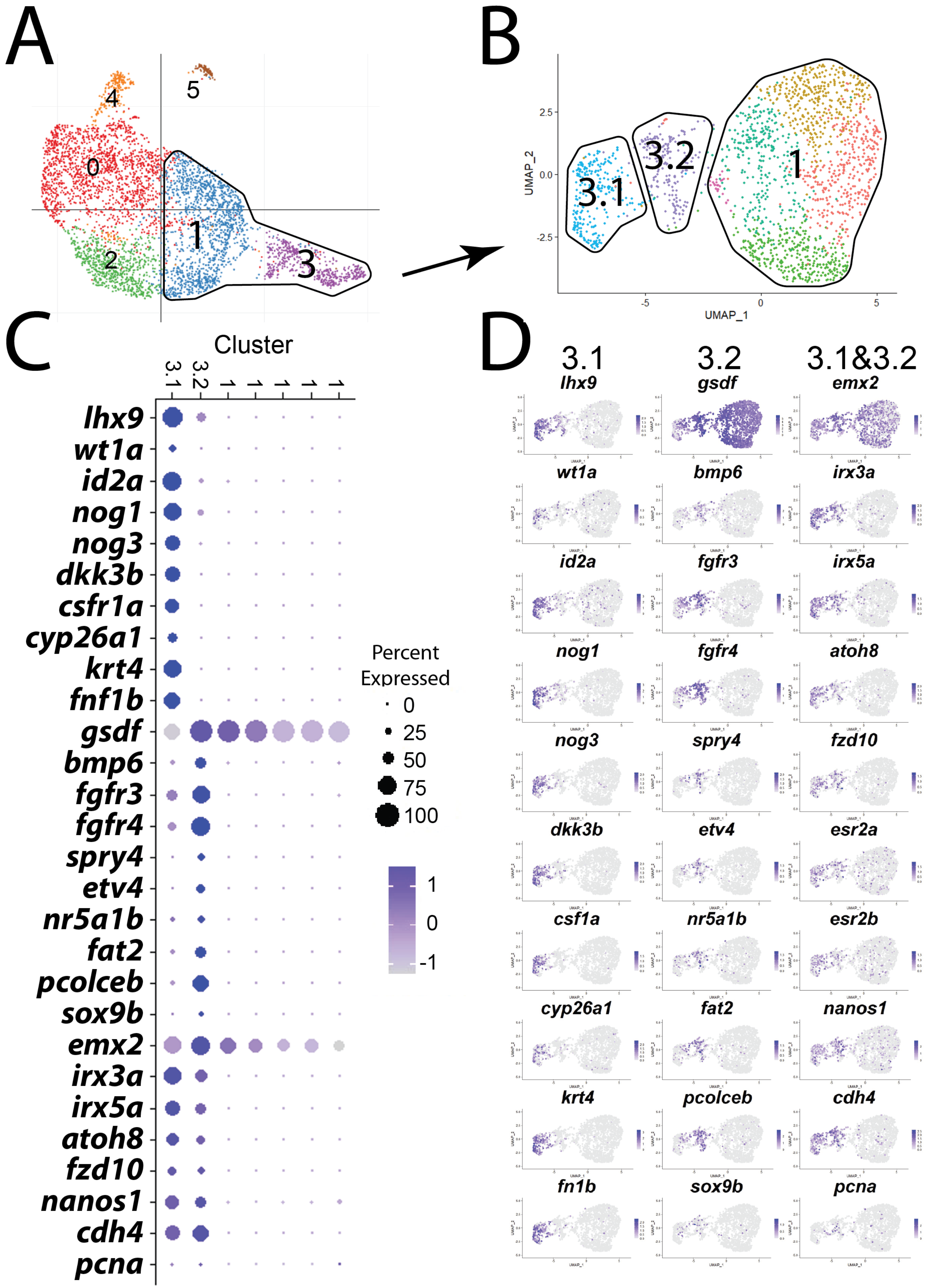

**Figure 4- figure supplement 3**

#### A. Retinoic Acid Production

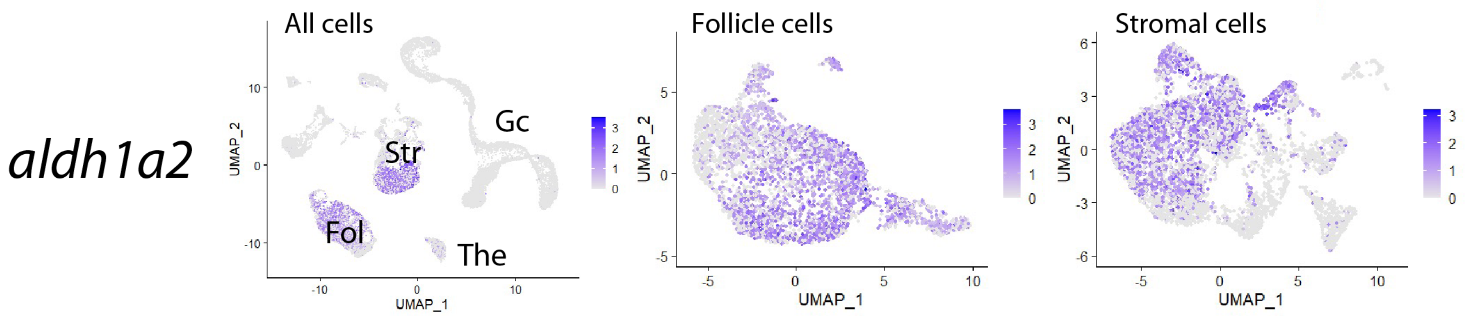


#### B. Retinoic Acid Degredation

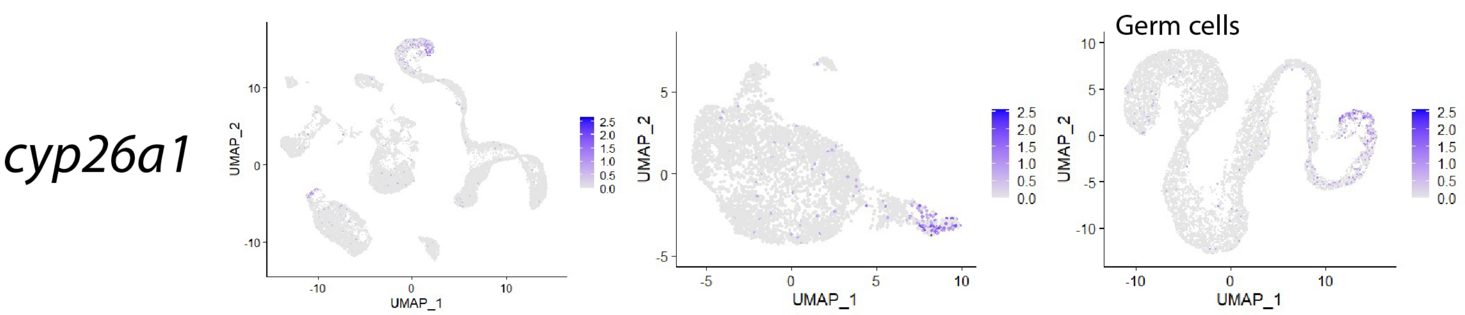


#### C. Retinoic Acid Receptors

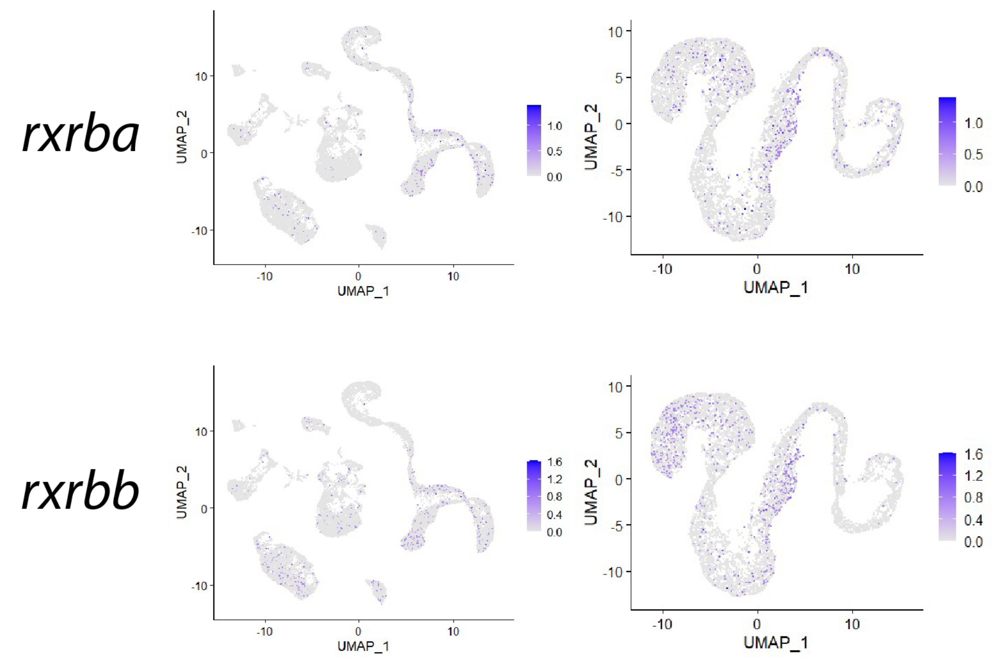

**Figure 4- figure supplement 4**

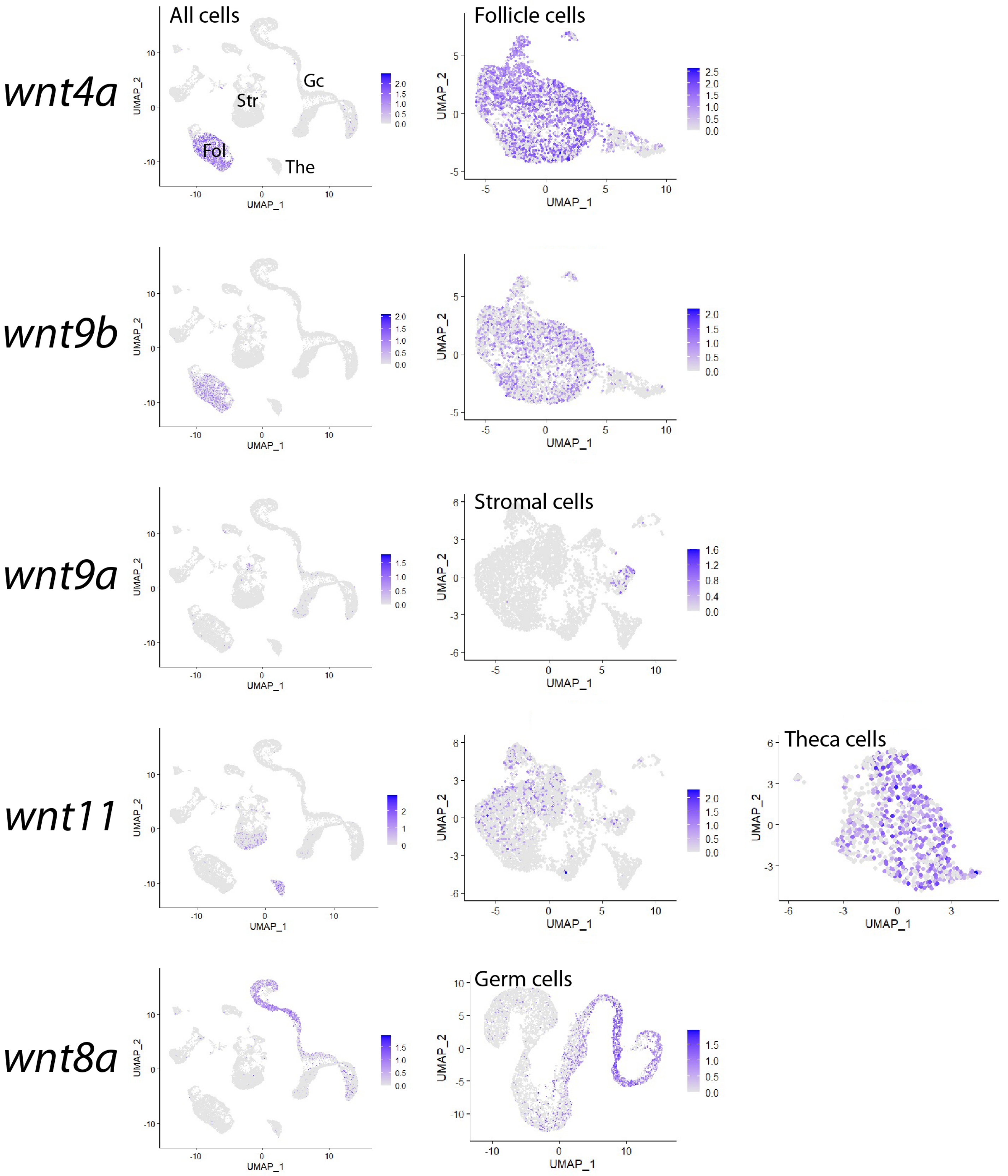

### Figure 6- figure supplement 1

#### A. Follicle cell subcluster GO terms

| Subcluster | id | source | term_id | term_name | term_size | p_value cluster.0 | p_value cluster.1 | p_value cluster.2 | p_value cluster.3 | p_value cluster.4 | p_value cluster.5 |
|---|---|---|---|---|---|---|---|---|---|---|---|
| 0 q-Stage II FC | 1 | GO:BP | GO:0006412 | translation | 519 | 3.0e-43 | NA | 2.7e-04 | NA | NA | NA |
| | 2 | GO:BP | GO:0043043 | peptide biosynthetic process | 526 | 5.6e-43 | NA | 3.0e-04 | NA | NA | NA |
| | 3 | GO:BP | GO:0043604 | amide biosynthetic process | 609 | 4.4e-40 | NA | 8.9e-04 | NA | NA | NA |
| | 4 | GO:BP | GO:0006518 | peptide metabolic process | 625 | 1.4e-39 | NA | 1.1e-03 | NA | NA | NA |
| | 5 | GO:BP | GO:0043603 | cellular amide metabolic process | 806 | 1.2e-34 | NA | 7.1e-03 | NA | NA | NA |
| 1 Stage I FC | 6 | GO:BP | GO:0019882 | antigen processing and presentation | 36 | NA | 1.1e-13 | NA | NA | NA | NA |
| | 7 | GO:BP | GO:0007423 | sensory organ development | 673 | NA | 1.8e-04 | NA | NA | 3.3e-02 | NA |
| | 8 | GO:BP | GO:0009653 | anatomical structure morphogenesis | 2275 | NA | 2.3e-03 | NA | NA | NA | 2.4e-03 |
| | 9 | GO:BP | GO:0048839 | inner ear development | 178 | NA | 2.3e-03 | NA | NA | NA | NA |
| | 10 | GO:BP | GO:0043583 | ear development | 178 | NA | 2.3e-03 | NA | NA | NA | NA |
| 2 Transitioning | 11 | GO:BP | GO:0042773 | ATP synthesis coupled electron transport | 56 | NA | NA | 3.9e-04 | NA | NA | NA |
| | 12 | GO:BP | GO:0046034 | ATP metabolic process | 149 | NA | NA | 6.5e-04 | NA | 9.0e-03 | NA |
| | 13 | GO:BP | GO:1902600 | proton transmembrane transport | 159 | NA | NA | 8.9e-04 | NA | NA | NA |
| | 14 | GO:BP | GO:0022904 | respiratory electron transport chain | 69 | NA | NA | 9.1e-04 | NA | NA | NA |
| | 15 | GO:BP | GO:0006119 | oxidative phosphorylation | 69 | NA | NA | 9.1e-04 | NA | 1.5e-02 | NA |
| 3 pre-FC | 16 | GO:BP | GO:0048856 | anatomical structure development | 4684 | 4.0e-02 | 2.8e-03 | NA | 6.4e-08 | NA | 2.8e-03 |
| | 17 | GO:BP | GO:0032502 | developmental process | 4900 | NA | 7.4e-03 | NA | 1.1e-07 | NA | 7.0e-03 |
| | 18 | GO:BP | GO:0009888 | tissue development | 1594 | NA | 5.2e-03 | NA | 4.0e-06 | NA | 1.9e-02 |
| | 19 | GO:BP | GO:0007275 | multicellular organism development | 4280 | 5.4e-03 | 1.4e-03 | NA | 2.0e-05 | NA | 1.7e-03 |
| | 20 | GO:BP | GO:0061448 | connective tissue development | 192 | NA | 1.4e-02 | NA | 1.2e-04 | NA | NA |
| 4 p-Stage II FC | 21 | GO:BP | GO:0006260 | DNA replication | 142 | NA | NA | NA | NA | 9.3e-35 | NA |
| | 22 | GO:BP | GO:0006259 | DNA metabolic process | 585 | NA | NA | NA | NA | 3.9e-27 | NA |
| | 23 | GO:BP | GO:0006261 | DNA-dependent DNA replication | 100 | NA | NA | NA | NA | 4.4e-25 | NA |
| | 24 | GO:BP | GO:0006281 | DNA repair | 363 | NA | NA | NA | NA | 2.0e-17 | NA |
| | 25 | GO:BP | GO:0034641 | cellular nitrogen compound metabolic process | 5168 | 9.9e-12 | NA | NA | NA | 2.5e-16 | NA |
| 5 | 26 | GO:BP | GO:0030198 | extracellular matrix organization | 161 | NA | NA | NA | NA | NA | 6.9e-06 |
| | 27 | GO:BP | GO:0043062 | extracellular structure organization | 161 | NA | NA | NA | NA | NA | 6.9e-06 |
| | 28 | GO:BP | GO:0048731 | system development | 3749 | NA | 1.3e-03 | NA | 1.9e-04 | NA | 1.0e-04 |
| | 29 | GO:BP | GO:0001501 | skeletal system development | 404 | NA | NA | NA | 2.6e-03 | NA | 7.0e-04 |
| | 30 | GO:BP | GO:0048589 | developmental growth | 401 | NA | NA | NA | NA | NA | 7.1e-03 |

#### B. Stromal cell subcluster GO terms

| Subcluster | id | term_id | term_name | p_value cluster.0 | p_value cluster.1 | p_value cluster.2 | p_value cluster.3 | p_value cluster.4 | p_value cluster.5 |
|---|---|---|---|---|---|---|---|---|---|
| 0 q-Interstitial | 1 | GO:0051085 | chaperone cofactor-dependent protein refolding | 1.1e-02 | NA | NA | NA | NA | NA |
| | 2 | GO:0051084 | 'de novo' posttranslational protein folding | 1.1e-02 | NA | NA | NA | NA | NA |
| | 3 | GO:0006458 | 'de novo' protein folding | 1.3e-02 | NA | NA | NA | NA | NA |
| | 4 | GO:0001945 | lymph vessel development | 1.4e-02 | NA | NA | NA | NA | NA |
| | 5 | GO:0061077 | chaperone-mediated protein folding | 2.8e-02 | NA | NA | NA | NA | NA |
| 1 pericytes | 6 | GO:0006412 | translation | NA | 7.0e-20 | NA | NA | NA | NA |
| | 7 | GO:0043043 | peptide biosynthetic process | NA | 9.3e-20 | NA | NA | NA | NA |
| | 8 | GO:0043604 | amide biosynthetic process | NA | 2.2e-18 | NA | NA | NA | NA |
| | 9 | GO:0006518 | peptide metabolic process | NA | 3.9e-18 | NA | NA | NA | NA |
| | 10 | GO:0043603 | cellular amide metabolic process | NA | 9.2e-16 | NA | NA | NA | NA |
| 2 VSM | 11 | GO:0050920 | regulation of chemotaxis | NA | NA | 4.1e-03 | NA | NA | NA |
| | 12 | GO:0031175 | neuron projection development | NA | NA | 2.9e-02 | NA | NA | NA |
| | 13 | GO:0010959 | regulation of metal ion transport | NA | NA | 4.2e-02 | NA | NA | NA |
| 3 Stromal progenitors | 14 | GO:0001501 | skeletal system development | NA | NA | NA | 2.5e-02 | NA | NA |
| | 15 | GO:0048385 | regulation of retinoic acid receptor signaling pathway | NA | NA | NA | 2.7e-02 | NA | NA |
| | 16 | GO:0030198 | extracellular matrix organization | NA | NA | NA | 3.4e-02 | NA | NA |
| | 17 | GO:0043062 | extracellular structure organization | NA | NA | NA | 3.4e-02 | NA | NA |
| 4 p-Interstitial | 18 | GO:0006259 | DNA metabolic process | NA | NA | NA | NA | 1.1e-27 | NA |
| | 19 | GO:0006260 | DNA replication | NA | NA | NA | NA | 1.5e-26 | NA |
| | 20 | GO:0006261 | DNA-dependent DNA replication | NA | NA | NA | NA | 8.1e-20 | NA |
| | 21 | GO:0051276 | chromosome organization | NA | NA | NA | NA | 9.7e-20 | NA |
| | 22 | GO:0006281 | DNA repair | NA | NA | NA | NA | 2.5e-18 | NA |
| 5 Epithelial | 23 | GO:0071696 | ectodermal placode development | NA | NA | NA | NA | NA | 6.9e-03 |
| | 24 | GO:0060114 | vestibular receptor cell differentiation | NA | NA | NA | NA | NA | 1.4e-02 |
| | 25 | GO:0060118 | vestibular receptor cell development | NA | NA | NA | NA | NA | 1.4e-02 |

**Figure 6- figure supplement 2**

#### A. Stromal subclsuter 1: Pericytes

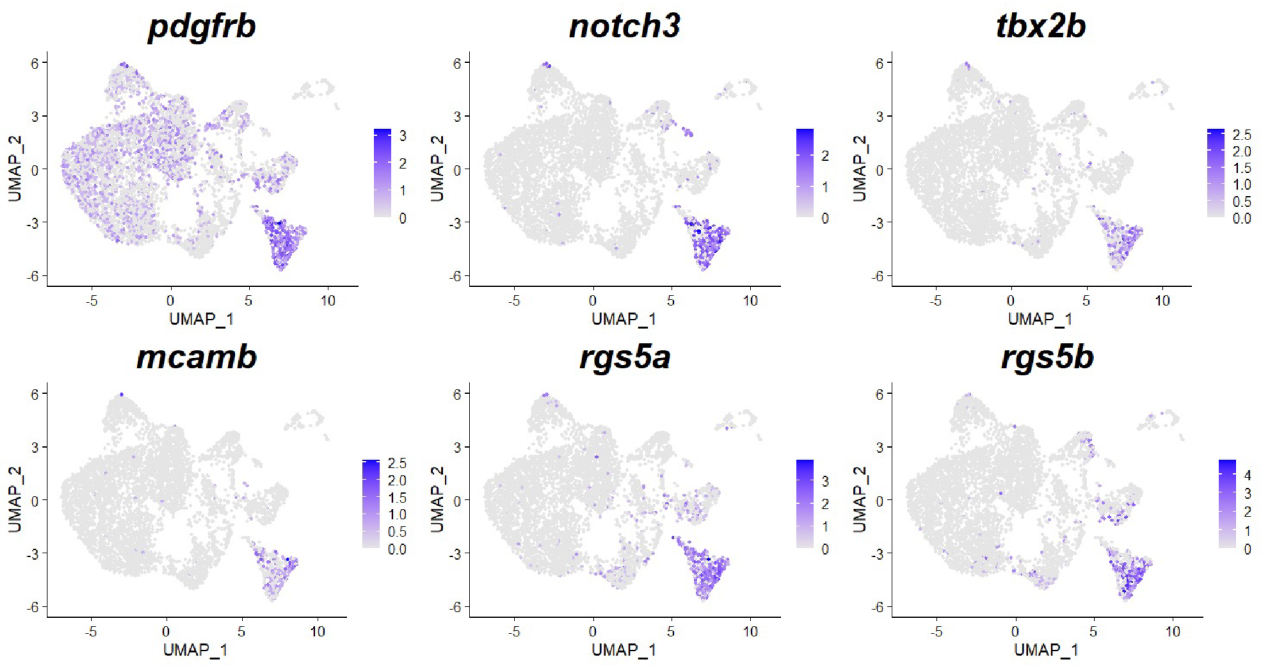


#### B. Stromal subcluster 2: Vascular smooth muscle

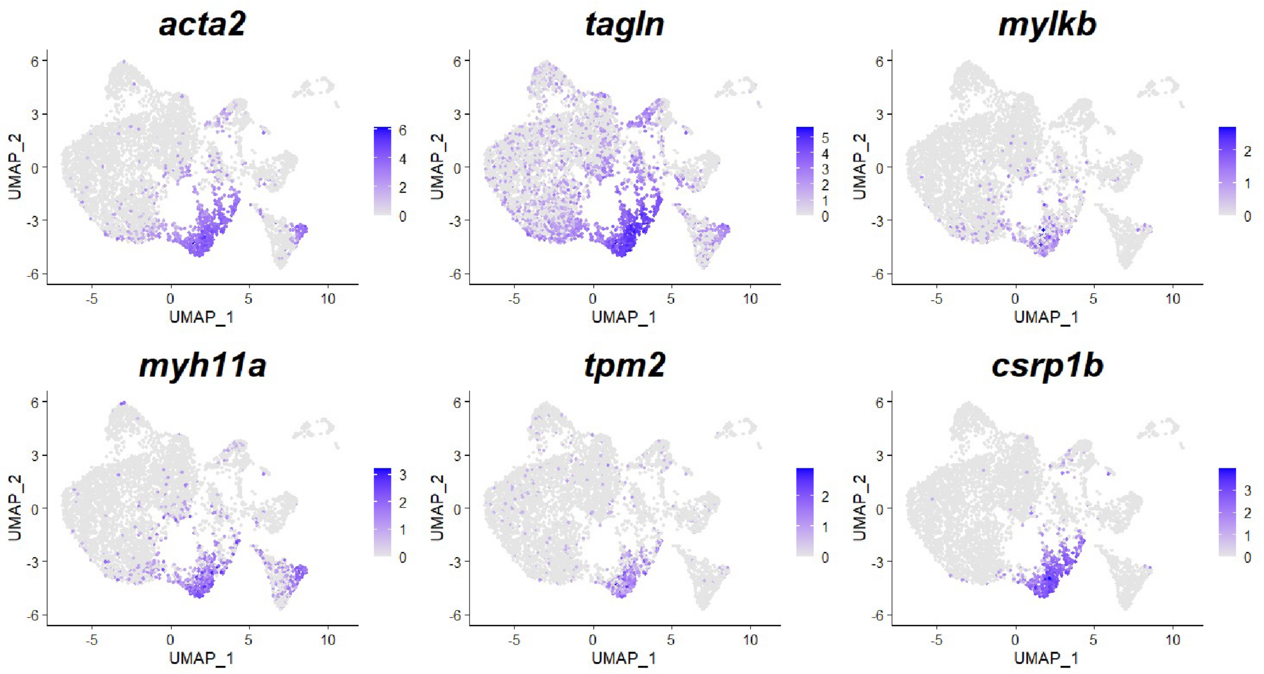

#### Figure 6- figure supplement 3

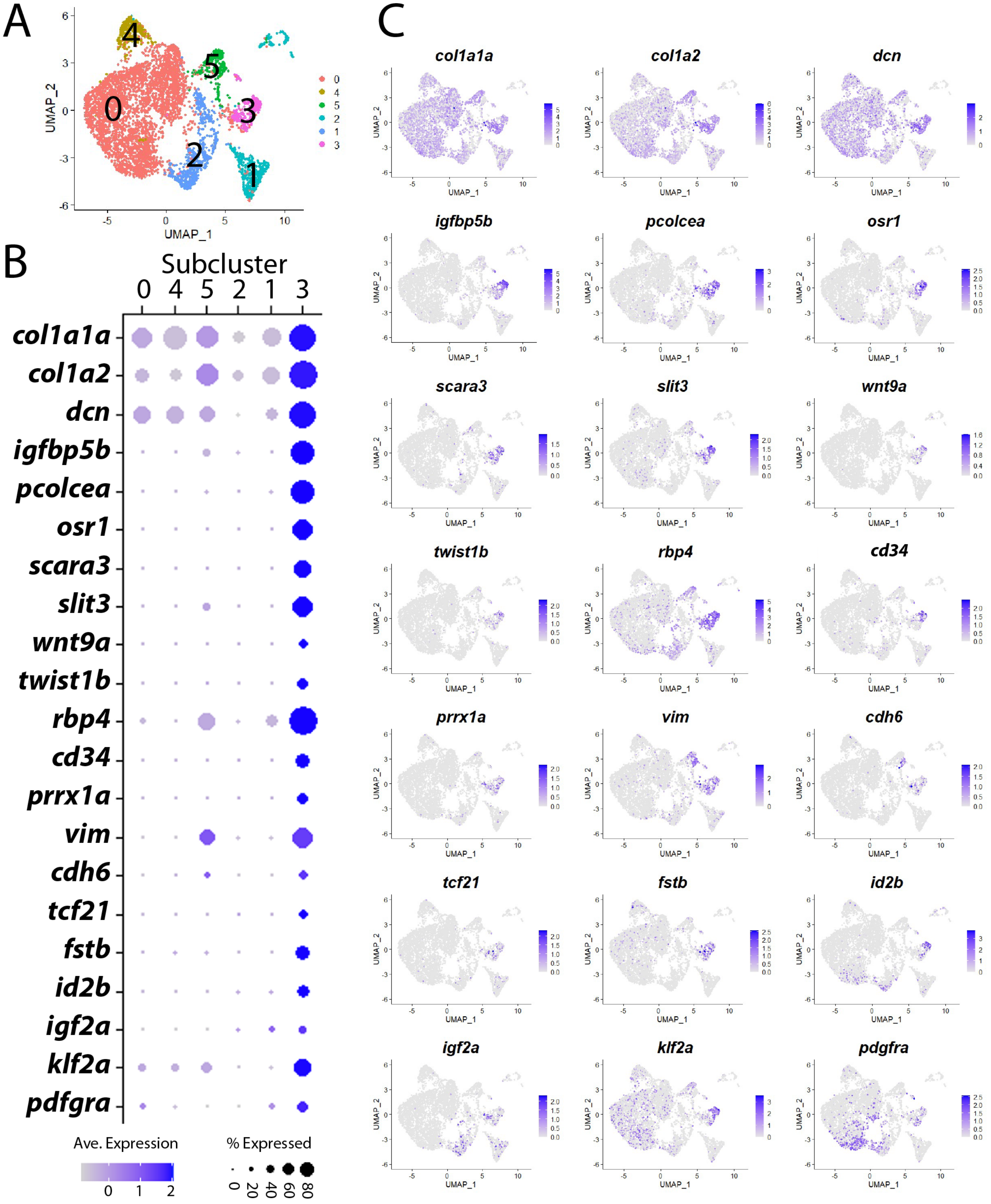

**Figure 7-figure supplament 1**

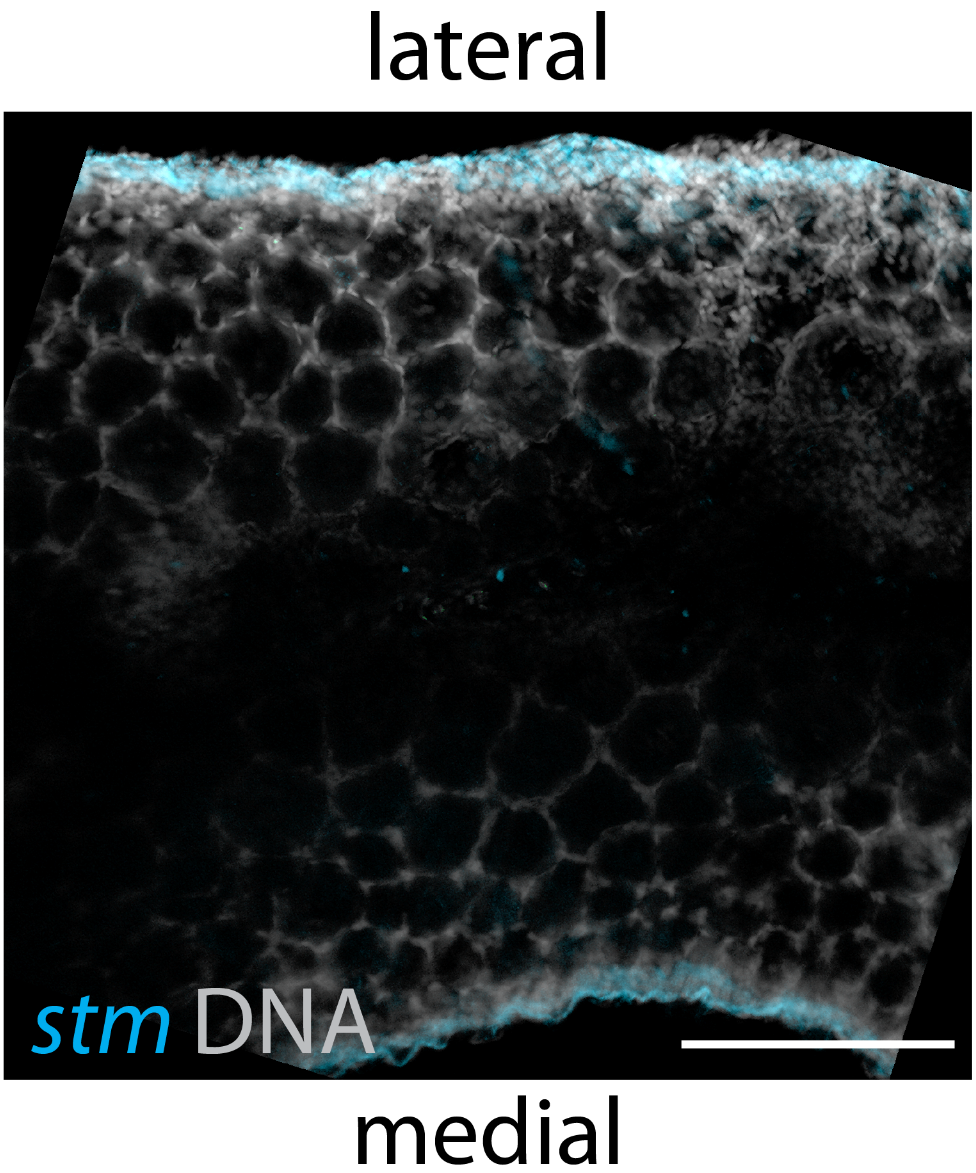

### Figure 8- figure supplement 1

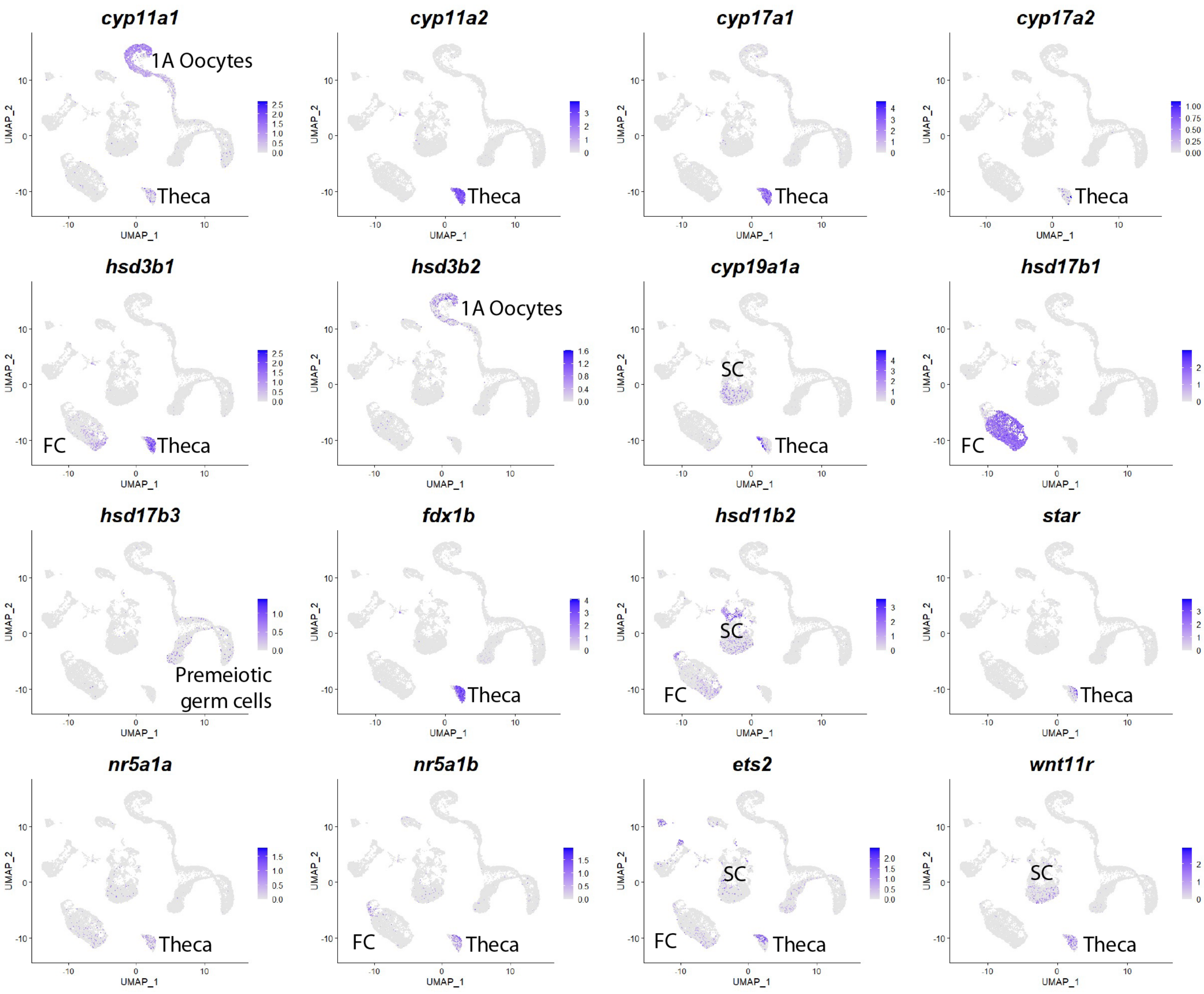

**Figure 9- figure supplement 1**

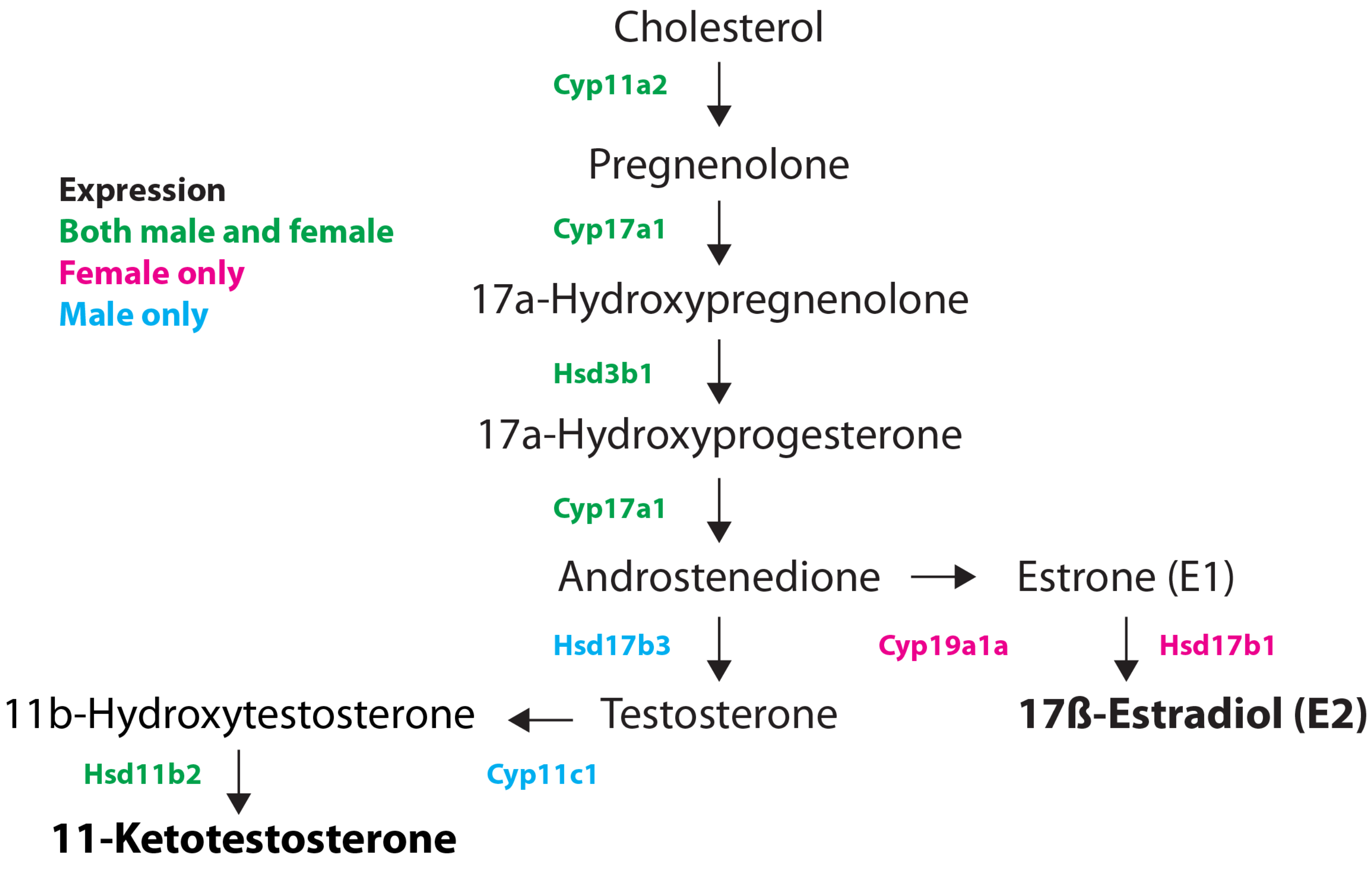
